# The Long Limb Bones of the StW 573 *Australopithecus* Skeleton from Sterkfontein Member 2: Descriptions and Proportions

**DOI:** 10.1101/497636

**Authors:** Jason L. Heaton, Travis Rayne Pickering, Kristian J. Carlson, Robin H. Crompton, Tea Jashashvili, Amelie Beaudet, Laurent Bruxelles, Kathleen Kuman, A.J. Heile, Dominic Stratford, Ronald J. Clarke

## Abstract

Due to its completeness, the A.L. 288-1 (“Lucy”) skeleton has long served as the archetypal bipedal *Australopithecus.* However, there remains considerable debate about its limb proportions. There are three competing, but not necessarily mutually exclusive, explanations for the high humerofemoral index of A.L. 288-1: (1) a retention of proportions from an *Ardipithecus-like* most recent common ancestor (MRCA); (2) indication of some degree of climbing ability; (3) allometry. Recent discoveries of other partial skeletons of *Australopithecus,* such as those of *A. sediba* (MH1 and MH2) and *A. afarensis* (KSD-VP-1/1 and DIK-1/1), have provided new opportunities to test hypotheses of early hominin body size and limb proportions. Yet, no early hominin is as complete (>90%), as is the ~3.67 Ma “Little Foot” (StW 573) specimen, from Sterkfontein Member 2. Here, we provide the first descriptions of that skeleton’s upper and lower long limb bones, as well as a comparative context of its limb proportions. As to the latter, we found that StW 573 possesses absolutely longer limb lengths than A.L. 288-1, but both skeletons show similar limb proportions. This finding seems to argue against a purely allometric explanation for A.L. 288-1’s limb proportions. In fact, our multivariate allometric analysis suggests that limb lengths of *Australopithecus,* as represented by StW 573 and A.L. 288-1, developed along a significantly different (p < 0.001) allometric scale than that which typifies modern humans and African apes. Our analyses also suggest, as have those of others, that hominin limb evolution occurred in two stages with: (1) a modest increase in lower limb length and a concurrent shortening of the antebrachium between *Ardipithecus* and *Australopithecus,* followed by (2) considerable lengthening of the lower limb along with a decrease of both upper limb elements occurring between *Australopithecus* and *Homo sapiens.*

## Introduction

Whether the earliest form of hominin terrestrial bipedalism was facultative or obligate, there is little debate that two-legged locomotion was a key adaptation of the earliest human ancestors, as indicated by their inferred functional anatomy (Lovejoy et al., 2009; Almécija et al., 2013), and, perhaps more crucially by ~3.66 Ma fossil footprints that occur at Laetoli G and S (Tanzania) (Leakey and Hay, 1979; Crompton et al., 2012; Hatala et al., 2016; Masao et al., 2016; Raichlen and Gordon, 2017). However, based on A.L. 288-1 (“Lucy”), a ~40 % complete skeleton of *Australopithecus afarensis,* the nature of early hominin bipedalism (e.g., efficiency and gait type) continues to be disputed (e.g., Lovejoy, 1979 versus Stern and Susman, 1983). Adding to the debate over ancestral locomotor patterns, the anatomy of the long limb bones of A.L. 288-1, and those of purported hominins of the terminal Miocene, such as *Ardipithecus* and *Orrorin,* as well as manual and pedal elements assigned to various species of early hominin, suggest to many that these ancient groups engaged in varying degrees of forelimb-powered climbing (Stern and Susman, 1983; Susman et al., 1984, 1985; Tuttle, 1985; Heinrich et al., 1993; Clarke and Tobias, 1995; McHenry and Berger, 1998; Richmond and Strait, 2000; Corruccini and McHenry, 2001; Dainton, 2001; Pickford et al., 2002; Galik et al., 2004; McHenry and Jones, 2006; Richmond and Jungers, 2008; Lovejoy et al., 2009b; Haile-Selassie et al., 2012; Pickering et al., 2018, 2019) Irrespective of this debate, energetic costs for animals with relatively long limbs and more extended joint postures are lower than those of animals with shorter limbs, because, for example, such a bauplan reduces the number of strides needed to cover a given distance (Jungers, 1982; Steudel-Numbers and Tilkens, 2004; Steudel-Numbers, 2006; Pontzer, 2007, 2012). By extension, it follows that a bent-knee, bent-hip gait, as commonly assumed by extant non-human apes when walking upright (Crompton et al., 2003, 2008, 2010), is far less efficient for walking, or even standing, than is a posture with an extended knee (Crompton et al., 1998; Sockol et al., 2007; Pontzer et al., 2009).

Against this background, it is now a prevailing view that morphometric indices – including especially relative limb element lengths (e.g., humerus-to-femur length) – can be used as proxies for assessing the effectiveness of bipedality (Hartwig-Scherer and Martin, 1991; McHenry and Berger, 1998; McHenry and Berger, 1998; Asfaw et al., 1999; Richmond et al., 2002; Haeusler and McHenry, 2004; Reno et al., 2005; Jungers, 2009; Ruff, 2009; Jungers et al., 2016). Such approaches are often confounded, however, by presumed intraspecific body size sexual dimorphism (BSSD), a prominent example being that of *A. afarensis* which has been reconstructed variably as displaying limited or moderate (Reno et al., 2003;; Reno and Lovejoy, 2015) to extreme BSSD (Gordon et al., 2008). Thus, Jungers et al. (2016) have suggested that the differences in body shape/size (e.g., the differences in the presumptive male *A. afarensis* A.L. 827-1 femur and that of the small female A.L. 288-1) may reflect differences in locomotor function, as is the case for some living forest cercopithecines.

Equally important, recent discoveries, such as KSD-VP-1/1, a partial skeleton from Woranso-Mille, Ethiopia (Haile-Selassie et al., 2010), demonstrate that some *A. afarensis* individuals possessed long limb bones (e.g., tibia) that are well within the length range of modern humans.

Unfortunately, there are very few early hominin skeletons that preserve complete or near-complete limb elements, and those that do often require reconstructions which are inherently unreliable (Korey, 1990; Richmond et al., 2002; Holliday, 2012). For example, various analyses of the associated, but fragmentary limb bones of the South African *Australopithecus* skeletons Sts 14 and StW 431 have led to disagreements about the degree to which *Australopithecus* and early *Homo* engaged in arboreal behaviors. Specifically, Robinson (1972) argued that *Australopithecus africanus* possessed a humerofemoral index that was like modern humans, and therefore, engaged in modern human-like bipedality. However, this conclusion was based upon an apparent overestimation of the femoral length of Sts 14, which was estimated (Lovejoy and Heiple, 1970) to be much shorter than originally reconstructed.

It is against this backdrop that the completeness of the StW 573 (“Little Foot”) *Australopithecus* skeleton, unmatched until the time (~1.5 Ma) of *Homo ergaster* KNM-WT 15000 Nariokotome (Kenya) skeleton, is of such great importance to our understanding of early hominin locomotion. Here, we present the full anatomical descriptions of the brachial (humerus), antebrachial (radius and ulna), thigh (femur), and leg (tibia and fibula) bones of StW 573, as well as calculations of standard osteological intra-and interlimb indices. The goal of this paper is to contribute to ongoing discussions of the evolution of hominin limb proportions. For the sake of brevity and clarity, detailed comparative interpretations of the functional morphology of these elements are presented in Heaton et al. (in prep.) and Crompton et al. (2018). Additionally, we did not seek to study the suite of variables that impact the relationship of size and morphological variation (i.e., allometry), but instead we sought, to determine if the limb lengths of *Australopithecus* could be attributed to similarities in allometric scaling. Therefore, we hypothesized that fossil *Australopithecus,* represented by StW 573 and A.L. 288–1, do not differ significantly from the allometric pattern observed in modern *H. sapiens.*

Our results indicate that the evolution of hominin limb proportions broadly occurred in two phases, with a modest increase in the lower limb length and concurrent shortening of the antebrachium between *Ardipithecus* and *Australopithecus* followed by considerable lengthening of the lower limb along with decreases of both upper limb elements occurring between *Australopithecus* and *H. sapiens.* The resulting differences in limb proportions, therefore, appear to be a result of different scales of allometry, as StW 573 (and A.L. 288-1) differ significantly from the modern human and ape allometric scales.

## Materials and Methods

The overall macroscopic condition of each specimen was recorded, including completeness, degree of subaerial weathering (Behrensmeyer, 1978), staining by manganese dioxide (a qualitative assessment: none, moderate, which indicates partial coating with manganese deposits, or heavy, which means near-totally or totally coating with deposits). Each specimen was examined carefully for bone surface modifications using 10× power magnification (e.g., Blumenschine et al., 1996). Fractured surfaces on all specimens were assessed concerning the “angle formed by the fracture surface and bone cortical surface” (Villa and Mahieu, 1991: 34). Typically, fracture angles on long limb bones that were created when the bone was “green” (i.e., before significant loss of a bone’s organic fraction and its desiccation) are usually either acute or obtuse, while angles created on dry long limb bones are usually right (Villa and Mahieu, 1991).

We used Mitutoyo™ digital calipers and Paleotech™ field osteometric boards to collect standard osteometric linear measurements and osteometric angular measurements with SPI^TM^ 0 – 180° protractors (Wilder, 1920; Martin, 1928; Martin and Saller, 1928; McHenry et al., 1976, 2007; Corruccini, 1978; Trinkaus, 1983; Buikstra and Ubelaker, 1994; Haile-Selassie et al., 2010). In most cases, due to the completeness of StW 573, these traditional methods could be used; however, we provide additional measurement details or explanations, as necessary, below.

Initially unsure of the impact of proximal damage on humeral length, we employed Pearson’s (2000) method utilizing the relative position of the deltoid tuberosity to predict length. We discovered that while the head and neck were damaged, this did not appear to affect humeral length, as measures of the fossil and predicted values were the same.

Traditional methods of predicting humeral torsion could not be used, due to this proximal damage. Therefore, we used the position of the bicipital groove following the method of Larson (1996) to predict humeral torsion

Neither femur preserves the greater trochanter; therefore, neck length was measured as the maximum distance between the lateral most edge of the femoral head to the estimated location of the intertrochanteric crest (McHenry and Corruccini, 1978). In StW 573, only a small inferior portion of the crest is preserved. As StW 573 does not possess a complete femur, femoral anteversion was estimated following Martin and Saller (1959) on the right femur. Biomechanical lengths of the femora and tibiae (Trinkaus et al., 1999), measures of the femoral condyles (Lague, 2002), indices of relative head width or relative length of the femoral neck (Napier, 1964), tibial arch angle (DeSilva and Throckmorton 2010), tibiotalar joint measures (DeSilva, 2009), and fibulotalar angles (Marchi 2015) were also measured. Moreover, while our aim is not to interpret or discuss all measures reported in this paper, we provide them, as a service to the scientific community.

Comparative limb lengths for African apes were taken from raw data (*Pan troglodytes*, n=66 and Gorilla gorilla, n=22) provided by Pontzer et al. (2010)^1^ or means (*Pan paniscus*) presented in Shea (1984). The limb lengths of modern *H. sapiens* (n=548) were collected by one of us (JLH) in the Terry Collection at the National Museum of Natural History (Smithsonian Institution).

For StW 573, limb lengths were obtained reliably on at least one of the antimeres, with the sole exception of the left femur (missing its proximal end), which required mirroring with the right. Limb indices, such as brachial (100 × radius length/humerus length), crural (100 × tibia length/femur length), humerofemoral (100 × humerus length/femur length) and intermembral, or IMI (100 × (humerus length + radius length)/(femur length + tibia length)) derive from Schultz (1937) and Aiello and Dean (1990). Abbreviations used include: AP = anteroposterior or anteroposteriorly; ML = mediolateral or mediolaterally; SI = superoinferior or superoinferiorly.

Within the fossil descriptions, individual elements have been assigned a letter designation (e.g., StW 573g) in order to facilitate ease of discussion. In some cases, individual limb elements (e.g., left femur) comprise articulating, and non-articulating, fragments; thus, numbers (e.g., /1) have been added to the specimen number/letter designations (e.g., StW 573m/1=left femoral head). However, the broader catalog number (e.g., StW 573m) may be used when generally talking about an element, as there is no question about the association between these fragments.

Our fossil comparisons are primarily limited to specimens assigned to pre-Homo *ergaster* taxa, and do not include specimens of more recent groups (e.g., *Homo heidelbergensis, Homo neanderthalensis* or *Homo naledi),* as modern proportions (Holliday, 2012) had primarily appeared by *H. ergaster* (e.g., KNM-WT 15000). In our analyses, we have chosen to focus on limb lengths relevant to the calculation of the various limb proportion indices (e.g., brachial, crural, humerofemoral and intermembral). Measurements of comparative fossil specimens were taken from the literature; data sources are provided throughout. Additionally, the taxonomic assignment used for each specimen follows that of the original source. The comparative context among extant groups^2^ and fossil elements is given for each element.

Last, we note that the antebrachium of StW 573 exhibits bilateral asymmetry, with significantly more longitudinally curved left than right forearm bones. We hypothesize that this condition is likely due to traumatic plastic deformation of the left radius and ulna during the life of the StW 573 individual (Heile et al., in preparation). Regardless, compared to their antimeres, the left antebrachial elements are also better preserved and less altered diagenetically. Thus, we calculated all relevant indices using measurements of the left radius and ulna.

Among most biological datasets, size is the dominant component of variation and allometry (Klingenberg, 2016). In our allometric analysis, we used Jolicoeur’s (1963) multivariate method to analyze the long bone lengths of *Gorilla, Pan* and *H. sapiens* samples. Within each extant sample, individuals were excluded from the allometric analysis of data if all four limb lengths were not known. Sylvester et al. (2008) have noted sexual variation in limb allometry among humans, but sex assignation among fossil hominin specimens is often problematic (Grine, 2013). Thus, we pooled sexes within our analyses. Further, our goal was not to refine the understanding of human (or ape) allometry, but rather to determine whether StW 573 and A.L. 288-1 deviate from the allometric axis of those populations.

For each group, a principal component analysis (PCA) of the covariance matrix was conducted on natural log (ln) transformed limb lengths (i.e., humerus, radius, femur, and tibia) as implemented in Paleontological Statistics, or PAST (Version 3.21; Hammer et al., 2001). If accounting for a large portion (e.g., > 80%; see Hammer and Harper, 2006) of the total variation within a sample, the first principal component (PC 1) can be used as a “size” axis for the sample (Klingenberg 2016). For all samples, the expected isometric value for each eigenvector coefficient was calculated following Jolicoeur (1963), as 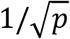, where *p* represents the number of variables. By dividing the PC 1 loading of each element (e.g., *ln* hum) by the expected isometric value (0.5), we were then able to obtain the allometric coefficients (*k* coefficient) for each element – *ln* hum, *ln* rad, *ln* fem, and *ln* tib. High values (*k* > 1.0) are indicative of positive allometry while low values (*k* < 1.0) denote negative allometry.

To test the statistical significance (p < 0.05) of the allometric coefficients, bootstrapping was used, as recommended by Kowalewski et al. (1997). To do this, we used an older version of PAST (Version 2.17: Hammer et al., 2001) in order to compute the 95% confidence intervals of each multivariate allometric coefficient using 2000 bootstrap replicates^3^. Significant deviations from isometry were accepted if the 95% confidence intervals did not include the value of one (1.0) within their range (Hammer and Harper, 2006).

By accounting for all traits together, PC 1 mean scores can be used as an index of size (Creighton and Strauss., 1986; Klingenberg and Spence., 1993; Klingenberg 1996a). Additionally, an individual’s position along this allometric axis (PC 1) can reveal information about its location within the range of the comparative sample’s size variation. For each extant group, the mean and standard deviation of PC 1 scores were computed.

Also, StW 573 and A.L. 288-1 were standardized to each population by first converting (*ln*-transforming) the raw limb measurements of StW 573 and A.L. 288-1. Then, principal component scores were computed for each fossil hominin. The principal component loadings (for each PC) were multiplied by their corresponding ln-transformed measurement (e.g., PC 1 humerus loading * *ln* humerus; where *ln* humerus represents the natural log transformation of the raw humerus measurement). For each component, the sum of these values (loadings * measurement) represents its principal component score, as shown in Equation 1 (for PC1). This process was carried out for the three remaining principal components (i.e., PC 2-4).

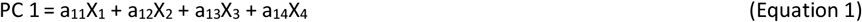

where a_11_ = first component loading for the humerus, a_12_ = first component loading of the radius, a_13_ = first component loading of the femur and a_14_ = first component loading of the tibia and while X_1_ = represents the measurement of the first variable (*ln* hum), X_2_ = measurement of the second variable (*ln* rad), X_3_ = measurement of the third variable (*ln* fem) and X_4_ = measurement of the fourth variable (*ln* tib)

In multivariate allometry, the axis of PC 1 is comparable to a regression line through the log-transformed data points. As noted by Klingenberg (1996b), components orthogonal to PC 1 (e.g., in our study, PC 2-4) can be treated as residuals in multivariate allometric analyses. The perpendicular placement of PC 2-4 describes each specimen’s deviation from the allometric axis (i.e., PC 1). Each individual’s placement in the Euclidean space is defined as the square root of the sum of squared values (PC scores) for PC 2-4 (Equation 2).

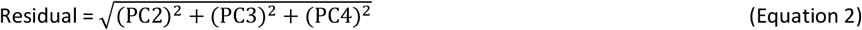

where PC2, PC3 and PC4 represent PC scores for a single specimen

Residuals were calculated for each extant species (*G. gorilla, P. troglodytes,* and *H. sapiens*) and fossil hominin (e.g., StW 573 and A.L. 288-1). For each extant group, the mean and standard deviation of the residuals were calculated, as well as the deviation of the two fossil hominins (StW 573 and A.L. 288-1) from each sample. Using Equation 3, a Z-score was computed for each residual and hominin/extant sample pair.

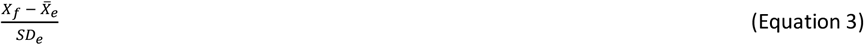

where *X_f_* = fossil hominin residual, 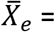 mean residual for the extant sample, and *SD_e_* = residual standard deviation of the extant sample

Evaluations of PC 1 served as a method to compare the ‘size’ of each fossil hominin to that of the extant populations. Remaining comparisons of the residuals (PC 2– 4) determined the likelihood of observing an individual of StW 573 and A.L. 288-1’s limb lengths along the allometric line (PC 1) of each extant group. The probability of observing the hominins within each population was calculated based upon their standardized value (e.g., Z-score) and Fourmilab Switzerland’s site (www.fourmilab.ch/rpkp/experiments/analysis/zCalc.html). Deviations along PC 1 (i.e., size) and from PC 1 (residuals) were considered significant if p < 0.05.

## Results

### Fossil Descriptions

Table 1 summarizes the general condition of the upper and lower limb bones of StW 573 with detailed descriptions of each element to follow.

**Table 1.** Catalog of StW 573 Long Limb Bones

| **Element (side)** | **Specimen Number** | **General condition <sup>a</sup>** |
| --- | --- | --- |
| Humerus (left) | **StW 573g** | Nearly complete, but both epiphyses damaged, proximal more so than distal; significant loss of periosteal lamellae |
| Humerus (right) | **StW 573h** | Complete, badly crushed, articulated to right, distorted scapula and partial right ulna; essentially unweathered |
| Radius (left) | **StW 573i** | Nearly complete but missing dorsal and anterolateral distal epiphysis and distal metaphysis and extreme distal diaphysis. |
| Radius (right) | **StW 573j** | Nearly complete, but missing distal epiphysis; proximal end badly crushed and distorted; heavily diagenetically altered cortex across most of specimen |
| Ulna (left) | **StW 573k** | Nearly complete, but missing distal epiphysis; mostly unweathered |
| Ulna (right) | **StW 573l** | Missing distal +1/3, crushed and distorted longitudinally; mostly unweathered |
| Femur (left) | **StW 573m** | Incomplete; missing neck and area near trochanters. Oblique fracture located in distal diaphysis. Preserved as two fragments: left femoral head (**StW 573m/1**) and left diaphysis and distal epiphysis (**StW 573m/2**) |
| Femur (right) | **StW 573n** | Complete, but femoral head crushed and slightly distorted. Fractured near distal diaphysis with significant cortical bone loss and weathering near fracture due to water action within cave. Preserved as right proximal femur (**StW 573n/1**) with head, neck and proximal diaphysis and right distal femur (**StW 573n/2**) with distal diaphysis and epiphysis. |
| Tibia (left) | **StW 573o** | Nearly complete, exhibits cortical exfoliation, especially near tuberosity. Fractured in distal 1/3 of diaphysis. Preserved as left proximal tibia and (majority of) diaphysis (**StW 573o/1**) and distal (1/3) of diaphysis and medial malleolus (**StW 573o/2**) |
| Tibia (right) | **StW 573p** | Complete, but heavily weathered. Fractured within talar surface and coursing obliquely through the medial malleolus. Exhibits heavy cortical exfoliation. |
| Fibula (left) | **StW 573q** | Fragments of left proximal fibular (**StW 573q/1**) and left distal fibula (**StW 573q/2**). Most of diaphysis missing, weathered |
| Fibula (right) | **StW 573r** | Fragment of right distal fibula, weathered |

<sup>a</sup> As of August 2018, physical and virtual reconstruction of the upper and lower limb elements of StW 573 were ongoing and may improve the condition of many specimens.

### The Humeri (Table 2, Figs. 1–3)

#### Preservation of StW 573g (left humerus)

The left humerus is complete. Most of its head is intact and in correct anatomical position, but the rest of that feature is obscured by the crushing and displacement of the anteromedial aspect of the proximal metaphysis. Specifically, the lesser tubercle of the specimen is crushed and deflected craniodorsally into the anterior portion of the head; the anteroproximal head was also fractured and dislocated superiorly as part of this mechanically induced shift in the proximal epiphysis of the bone. This displaced portion of the head now rests above the natural plane of the head, as indicated by the undistorted medial and posterior portions of that feature. Similarly, the proximal portion of the intertubercular groove is compressed dorsomedially into the medullary cavity; some of this damaged area has been re-enforced with plaster. Opposite this damage to the proximal intertubercular groove, the dorsolateral aspect of the proximal metaphysis is stove in ventromedially just below the distal rim of the head. The anterior aspect of the distal metaphysis is also damaged and repaired with plaster; this line of repaired damage continues onto the superior and anterior portions of the capitulum. The left humerus is more poorly preserved than are the left radius and ulna; most of its cortical bone surface shows significant diagenetic loss of lamellae. A SI ~45.0 mm unaltered stretch of the bicipital groove is unweathered, however. There is no indication of biotically induced damage on the specimen.

**Figure 1.**
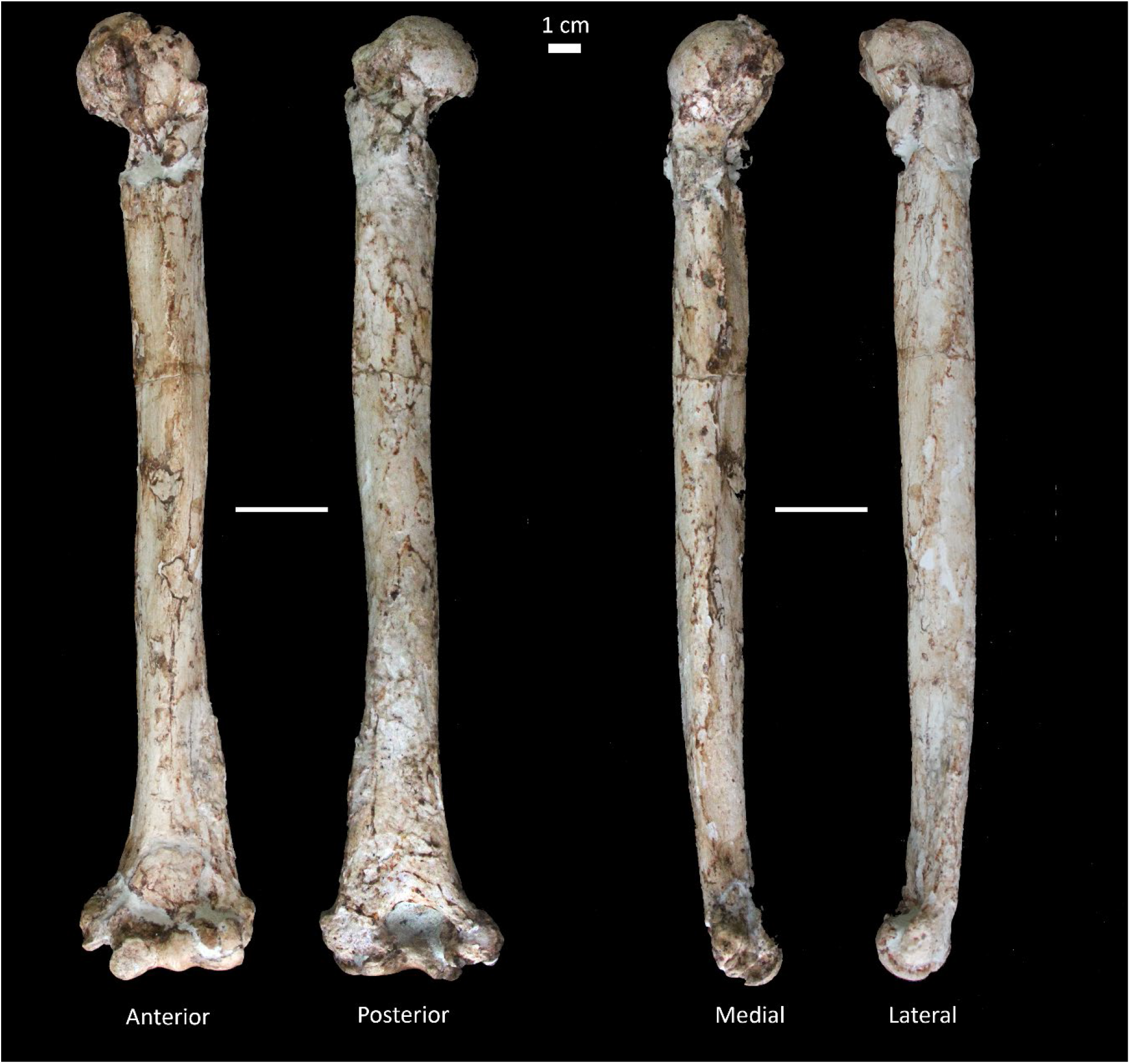
Left humerus of StW 573. Shown in anterior, posterior, medial and lateral views.

**Table 2.** Humeral osteometries and indices of StW 573. Estimated values are shown in square brackets.^a^

|  | StW573g<br>(left) |
| --- | --- |
| **Raw measures (mm)** |  |
| Maximum length | 290.0 |
| Physiological length | 290.0 |
| AP diaphysis diameter (midshaft) | 22.8 |
| ML diaphysis diameter (midshaft) | 22.3 |
| Diaphysis circumference (midshaft) | 69.0 |
| Trochlear AP width | 13.8 |
| Trochlear ML width | 20.5 |
| Lateral trochlear ridge AP diameter | 19.6 |
| Capitular ML width | 14.9 |
| Capitular SI height | 17.6 |
| Distal articular surface ML breadth | 40.0 |
| Biepicondylar ML breadth | 54.0 |
| Trochlea to medial epicondyle width | 36.7 |
| Capitulum to lateral epicondyle width | 25.7 |
| ML width of olecranon fossa | [20.0] |
| Olecranon fossa depth | [8.0] |
| ML width of medial wall of olecranon fossa | 10.0 |
| ML width of lateral wall of olecranon fossa | 15.5 |
| AP diaphysis diameter (superior to olecranon fossa) | 15.8 |
| ML width of medial epicondyle | 11.5 |
| **Angular measure (°)** |  |
| Torsion <sup>b</sup> | 120 |
| **Indices <sup>c</sup>** |  |
| Deltoid tuberosity position/maximum humeral length | [0.49] |
| Distal articular breadth/biepicondylar breadth | 0.74 |

<sup>a</sup> Due to its extremely distorted condition (see text), values for the right humerus (StW 573h) are not given.
<sup>b</sup> Humeral torsion was calculated via a method described in Larson (1996).
<sup>c</sup> Indices are described in Churchill et al. (2013).

#### Morphology of StW 573g (left humerus)

The left humerus is rugose. Although somewhat distorted, its head, which assumes a low torsional angle (Figure 2), is elliptical. The anatomical neck is partially obscured anteriorly and medially, but it presents as a shallow, unremarkable groove dorsolaterally. The greater tubercle is slightly altered but appears low and humanlike in position. The bone’s lateral ridge is rugose and is indented medially at about its midpoint. This inflection point probably represents the divide between the separate insertion areas for the supraspinatus and infraspinatus; if so, the supraspinatus facet appears more substantial. Both facets are elevations, and neither assumes an ovoid and/or concave form. The intertubercular groove is wide transversely (~10.0 mm at its widest point), and although shallowly excavated, is bounded by highly developed crests of the greater and lesser tubercles. The crest of the greater tubercle is well-elevated and elongated SI, imbuing the proximal half of the diaphysis with a strong triangular cross-section, with the anterolateral and anteromedial surfaces of the humerus sloping markedly away from one another. (Observations of the right humerus corroborate these characterizations of the anterior aspect of the proximal portion of the left element.) The crest for the origin of the lateral head of the triceps brachii appears to have been more subtly developed than are the previously mentioned linear elevations, but we note again that the posterior surface of the bone is poorly preserved and that this likely impacts the current expression of features on that surface. As it now appears, the gently expressed triceps brachii crest curves laterally about two-thirds of the way down the posterior diaphysis to connect with the well-developed and widely flaring brachioradialis crest (or lateral supracondylar ridge). The anterolaterally placed deltoid tuberosity is strongly developed and has quite a long SI, emerging ~80.0 mm distal to the most proximal point of the humeral head and coursing distally to near mid-diaphysis. Its exact distal extent was obliterated by the loss of bone cortex, but we estimate its deltoid tuberosity position (as indicated by the distance between the most proximal point on the humeral head to the most distal point on the deltoid tuberosity and measured parallel to the shaft axis) as quite low, at ~145.0 mm. The lateral epicondyle and especially the medial epicondyle of the specimen are strongly expressed and bound a well-defined distal articular surface. The lateral epicondyle is elongated SI, about half of it placed superior to the capitulum. Although damaged and partially reconstructed, the coronoid and radial fossae appear well-excavated to approximately the same depth. The olecranon fossa is wide ML. The specimen lacks a septal aperture. Any potential evidence of a nutrient foramen has been obscured or obliterated.

**Figure 2.**
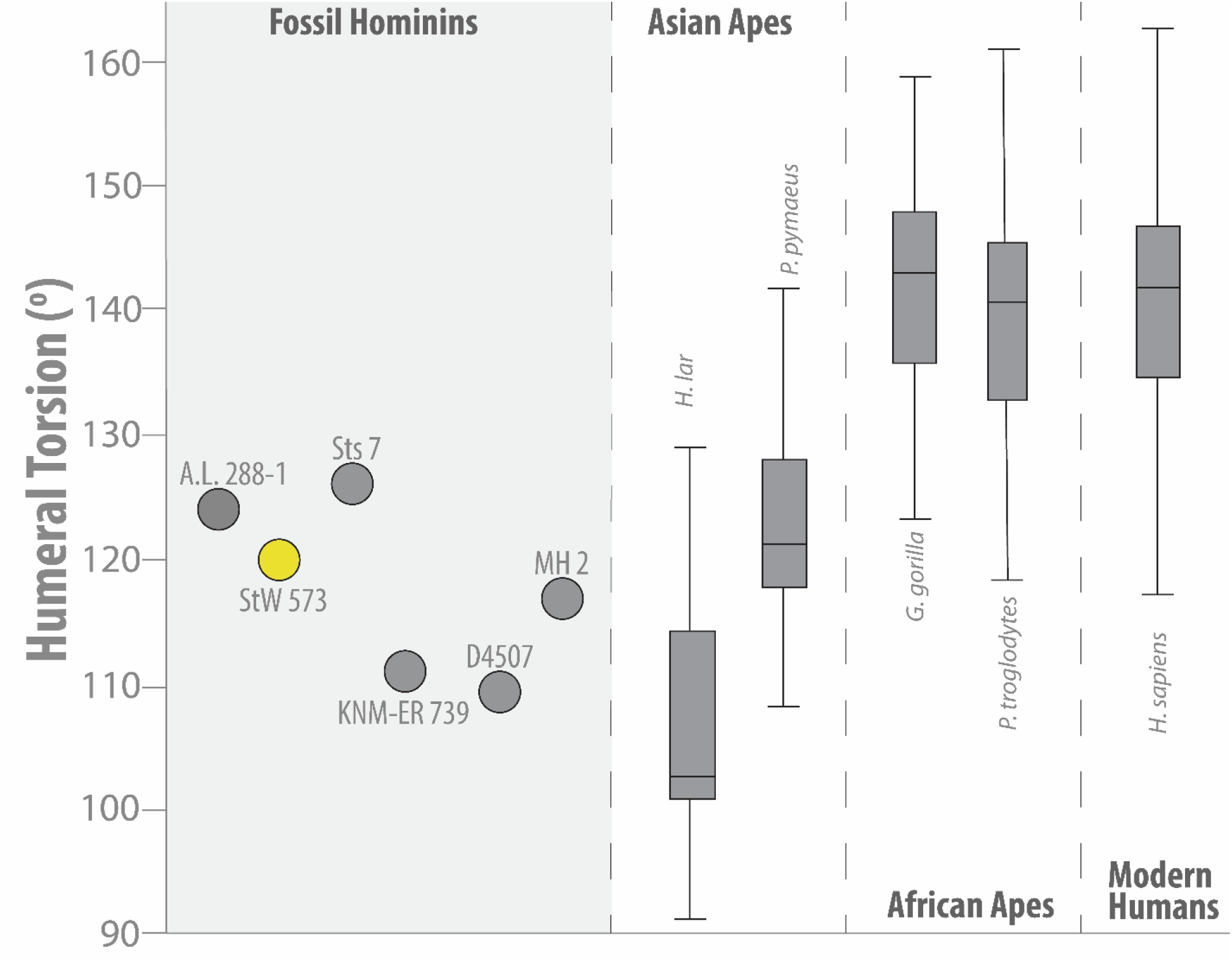
Humeral Torsion (in degrees). Shown are humeral torsion values for the apes, modern humans and fossil hominins including StW 573. Circles represent mean values (extant species) or individual values (fossil specimens) while whiskers represent the minimum and maximum values. Adapted from Larson (2009). Box-and-whisker plots denote the mean (dark horizontal line), upper and lower quartiles (boxes) and range (whiskers). Outliers (for *H. lar* at ~76° and *P. pygmaeus* at ~93° and ~154°) not shown.

**Figure 3.**
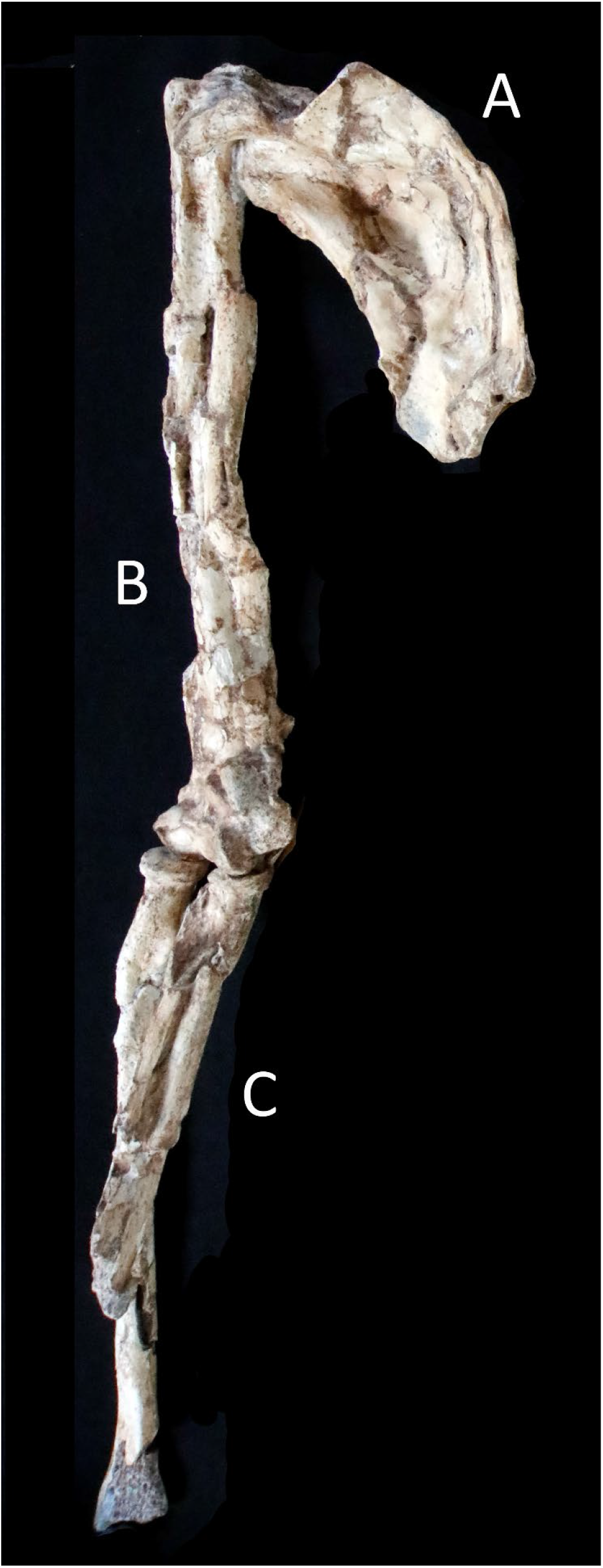
Articulated right upper limb and scapula of StW 573. Shown here are the right (A) scapula, (B) humerus and (C) radius and ulna (photo taken in January 2017).

#### Preservation of StW 573h (right humerus)

The right humerus of StW 573 is complete. From its proximal end to roughly the distal extent of the intertubercular groove (SI length = ~86 mm), the specimen is fairly undistorted, with an intact medullary cavity. It is, however, unfortunate that the humeral head is welded in rough articulation with the right scapula, obscuring that important feature. Distal to this undistorted proximal segment of the specimen, the diaphysis and distal metaphysis of the bone are twisted from original form and crushed AP, assuming an unnatural ML splay, under which the medullary cavity is completely collapsed. Distally, the trochlea is largely undistorted, but the rest of the distal epiphysis (like much of the diaphysis) is compressed AP. The right proximal ulna is cemented to the distal epiphysis of the humerus in approximate anatomical extension. The bone surface of the humerus is in good condition (weathering stage = 0, except for very occasional flaked patches of cortical lamellae), but is obscured by sporadic adhering breccia, calcite, and small manganese dioxide deposits. The specimen shows no indication of biotically induced modifications.

#### Morphology of StW 573h (right humerus)

We defer description of the highly distorted right humerus until virtual reconstruction of the element is completed. We refer readers to the description above of its less altered antimere.

### The Radii (Table 3, Fig. 4)

#### Preservation of StW 573i (left radius)

The left radius is nearly complete but is missing the dorsal and anterolateral distal epiphysis and distal metaphysis and extreme distal diaphysis. The specimen is composed of, starting proximally, three large pieces glued together, and another six smaller pieces glued together to form the anteromedial distal end of the bone; all contacts are tight and secure. There is a large, diagenetically induced hole, lined with calcite and breccia, in the dorsal portion of the rim of the radial head. The radius is stained lightly with sporadic circular deposits of manganese dioxide and shows very occasional tiny incidences of adhering breccia. The ventral surface of the entire specimen, as well as its whole neck, is unweathered. The rest of the dorsal surface of the radius has suffered the diagenetic loss of bone cortex, where multiple lamellae were removed in large, contiguous swaths. Similar but much smaller and localized circular incidences of lamellar loss occur sporadically across the rest of the specimen. The intact periosteal bone surface evinces no biotically induced damage, although there a few incidences of random striae distributed across (especially the distal end of) the specimen.

**Figure 4.**
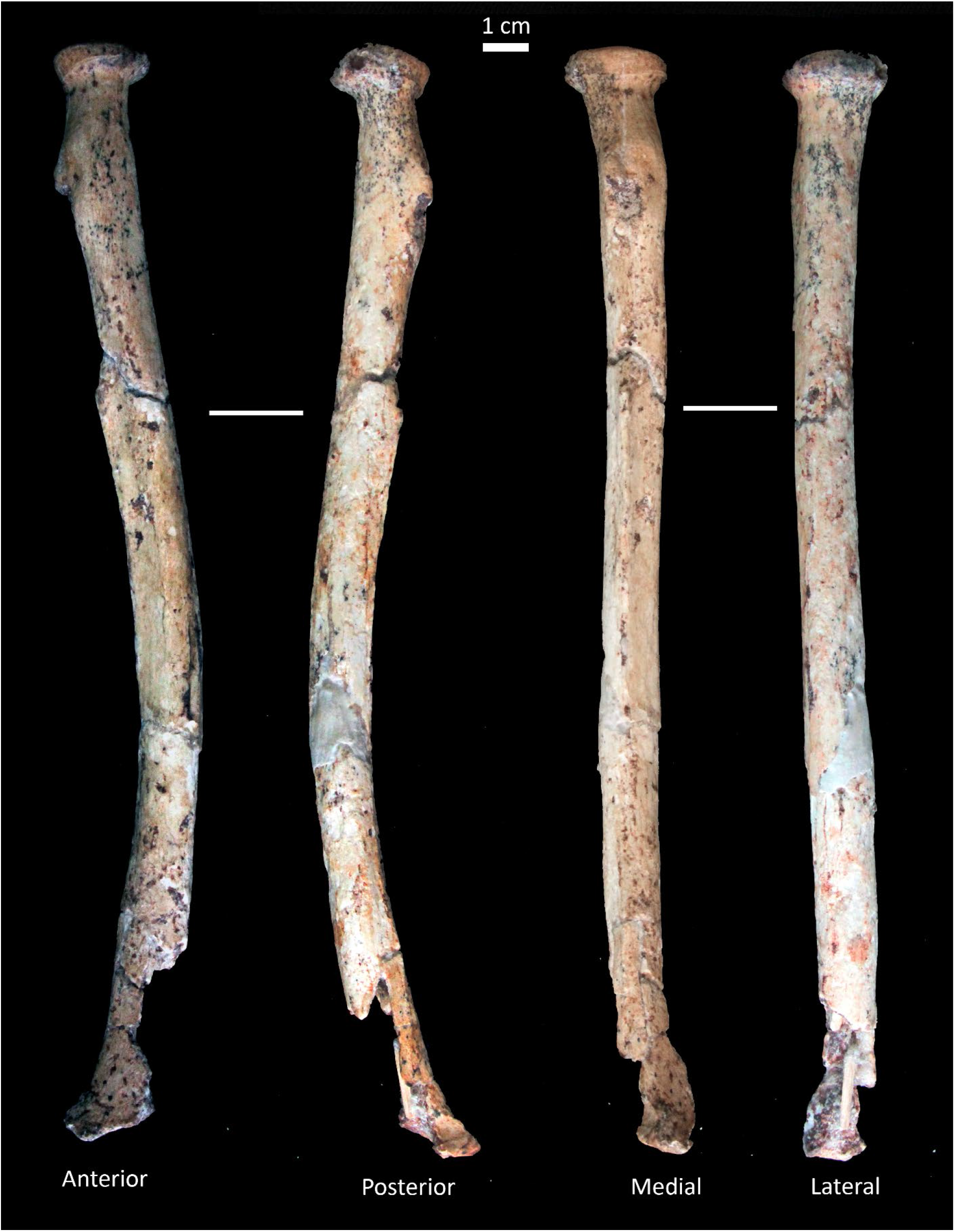
Left radius of StW 573. Shown in anterior, posterior, medial and lateral views.

**Table 3.**
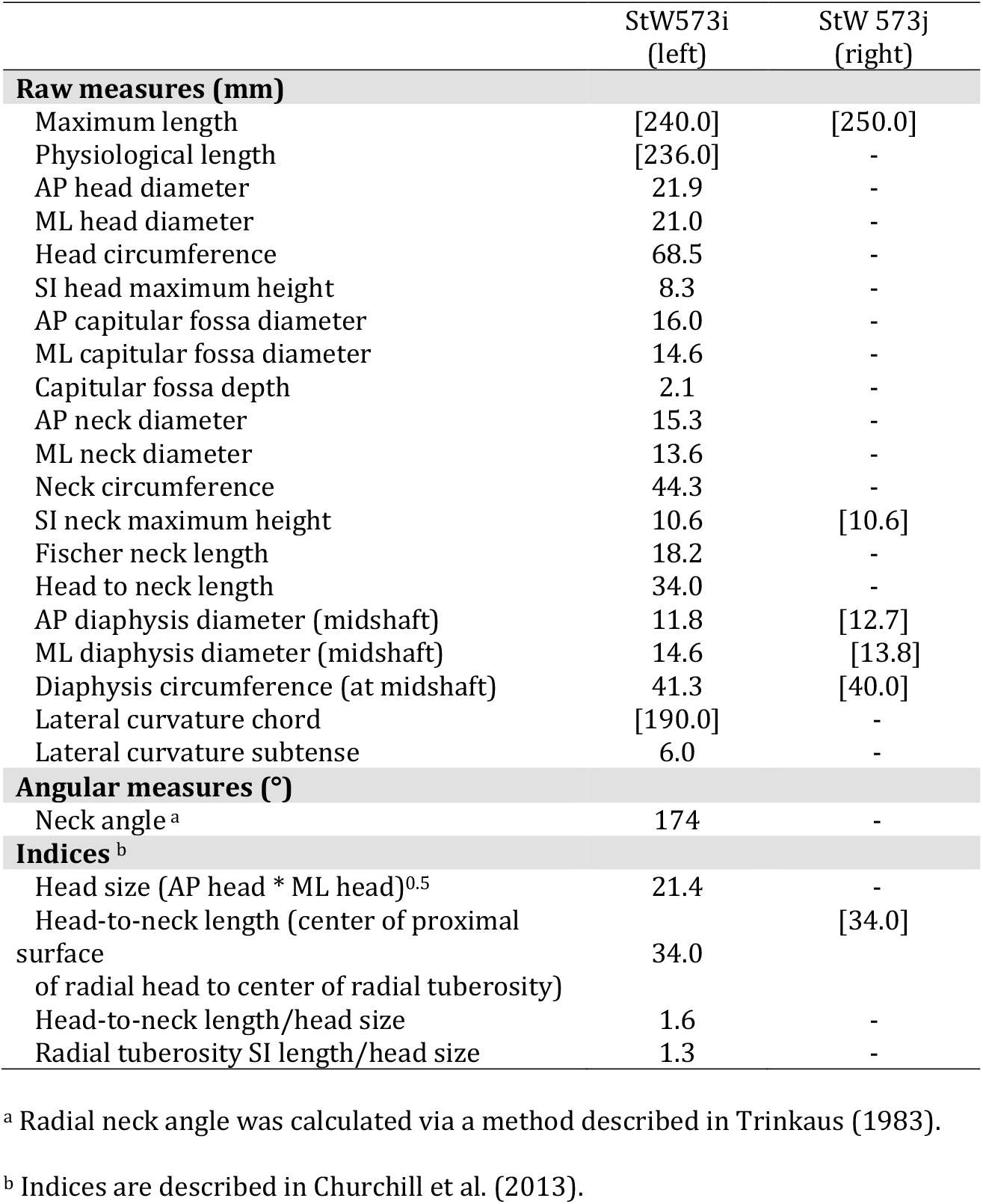
Radial osteometrics and indices of StW 573. Estimated values are shown in square brackets.

|  | StW573i<br>(left) | StW 573j<br>(right) |
| --- | --- | --- |
| **Raw measures (mm)** |  |  |
| Maximum length | [240.0] | [250.0] |
| Physiological length | [236.0] | - |
| AP head diameter | 21.9 | - |
| ML head diameter | 21.0 | - |
| Head circumference | 68.5 | - |
| SI head maximum height | 8.3 | - |
| AP capitular fossa diameter | 16.0 | - |
| ML capitular fossa diameter | 14.6 | - |
| Capitular fossa depth | 2.1 | - |
| AP neck diameter | 15.3 | - |
| ML neck diameter | 13.6 | - |
| Neck circumference | 44.3 | - |
| SI neck maximum height | 10.6 | [10.6] |
| Fischer neck length | 18.2 | - |
| Head to neck length | 34.0 | - |
| AP diaphysis diameter (midshaft) | 11.8 | [12.7] |
| ML diaphysis diameter (midshaft) | 14.6 | [13.8] |
| Diaphysis circumference (at midshaft) | 41.3 | [40.0] |
| Lateral curvature chord | [190.0] | - |
| Lateral curvature subtense | 6.0 | - |
| **Angular measures (°)** |  |  |
| Neck angle <sup>a</sup> | 174 | - |
| **Indices <sup>b</sup>** |  |  |
| Head size (AP head * ML head)<sup>0.5</sup> | 21.4 | - |
| Head-to-neck length (center of proximal surface of radial head to center of radial tuberosity) | 34.0 | [34.0] |
| Head-to-neck length/head size | 1.6 | - |
| Radial tuberosity SI length/head size | 1.3 | - |

<sup>a</sup> Radial neck angle was calculated via a method described in Trinkaus (1983).
<sup>b</sup> Indices are described in Churchill et al. (2013).

#### Morphology of StW 573i (left radius)

The specimen is strongly longitudinally curved. It possesses a prominent, circular head, with a distinct anteromedial bevel, which, in turn, yields eccentric placement of the capitular fossa. At its extreme, the bevel is elevated only ~4.0 mm above the inferior rim of the head. The maximum SI height of the radial head is just posterior to the position of the radial tuberosity. The capitular fossa is modestly deep. The radial neck is short SI, with its maximum length taken medially from distal edge of head rim to superior margin of radial tuberosity and is not especially constricted. A prominent radial tuberosity emerges just distal to the medial aspect of the neck and spans the width of the proximomedial diaphysis (maximum SI length = 27.0 mm; maximum width taken transverse to SI length = 14.0 mm). The anterior border of the tuberosity is low and blunt; the posterior border comprises a short (SI length = 7.6 mm), medially projecting enthesophytic flange proximally, which is truncated distally by a deep rectangular depression. A contiguous arc of low enthesophytic growth rims this laterally invasive depression anteriorly, posteriorly and distally. The anterior oblique line emerges from the anterodistal extremity of the radial tuberosity as a blunt, anterolaterally trending crest. Its lateral margin drops sharply, forming a steep anterolateral border of the proximal diaphysis. In sum, the attachment surfaces for the biceps brachii, the oblique cord, and the flexor digitorum superficialis, as well as for the inferoanterior fibers of the supinator are broad and well-developed. The proximal root of the interosseous border is placed 63.0 mm distal to the distal margin of the medial rim of the radial head. The border is only moderately developed, presenting as a sharp elevation for only ~31.0 mm from its proximal root distally, and then continues inferiorly as a low wall separating the anterior and posterior surfaces of the mid-to distal diaphysis at an obtuse angle. The interosseous border terminates distally at the distoanterior corner of the ulnar notch. Missing bone and diagenetic loss of cortex disallow observations of other surfaces where muscles and ligaments attached to the specimen and obscure the nutrient foramen.

#### Preservation of StW 573j (right radius)

Six separate pieces, reconnected with glue along good joins, comprise the nearly complete right radius. The specimen only lacks the anterior, medial and posterior portions of its distal metaphysis and distal epiphysis, as well as its styloid process. Trabeculae lining the endosteal wall of the lateral distal metaphysis and epiphysis are filled with breccia and calcite, attesting to the antiquity of the loss of the rest of the distal portion of the bone. Likewise, all fracture edges in this region are smoothed and coated in breccia and calcite; the edges also approximate right angles, indicating that the bone was leached of at least much of its organic fraction when the breakage occurred. Four connected pieces, starting just proximal to the single distal fragment described above span proximally for a combined (non-anatomical) SI length of ~170.0 mm and represent a fairly undistorted length of the diaphysis just distal to the distal margin of the radial tuberosity. There is a large SI elongated wedge (maximum length = ~30.0 mm) of missing bone along the distomedial edge of this diaphyseal span, but most of the unit is intact and encloses an unaltered medullary cavity. Proximally, the radial head, neck, and tuberosity contribute to the final segment of the specimen, which is crushed flat ML, with the distorted lateral rim of the head levered superomedially. This portion of the rim of head appears to be is missing its outer cortical layer(s). Although there are some areas of unweathered cortex on the radius, most of the fossil’s surface is severely altered diagenetically, with cracked, flaking surfaces and loss of lamellae in broad, longitudinally trending swaths or as small circular incidences. The specimen carries frequent deposits of breccia, calcite and manganese dioxide, but shows no sign of biotically induced modifications.

#### Morphology of StW 573i (right radius)

The right radius is fairly robust. Because the proximal end of the specimen is so severely damaged, we restrict our discussion of the morphology of that portion of the bone and refer readers to the description of the undistorted left radius for more detail about the proximal radial anatomy of StW 573. The superior surface of the head of the right radius appears moderately cupped, and the inferior margin of that feature’s rim is sharp, clearly demarcating the superior extent of what appears to be a short, only moderately constricted radial neck. The radial tuberosity is large and superoinferiorly elongated (maximum SI length = ~22.5 mm; maximum ML breadth = ~14.7 mm), with a particularly well-developed, medially bulging posterior border for the insertion of the biceps brachii. The anterior border of the tuberosity appears to be less projecting and smoother than is the posterior border. Distortion of the proximal portion of the radius disallows calculation of the bone’s lateral curvature chord and subtense. However, simple visual inspection of the specimen makes clear that the mid-diaphysis of the specimen, unlike that of its antimere, is relatively straight. The midshaft also carries a delicate but steeply developed interosseous border, which is consistently sharp for its preserved length and which imbues the radial shaft with a gentle triangular cross-sectional shape. Correcting for distortion, the proximal root of the interosseous border is placed ~60.0 – ~65.0 mm from the distal margin of the radial head’s medial rim. A minuscule nutrient foramen (<1.0 mm in maximum dimension) occurs ~8.0 mm lateral to this root, on a blunt crest that represents the distal extremity of the anterior oblique line. Crushing (proximally) and loss of bone surface (proximally and across much of the rest of the diaphysis) has obliterated or obscured other important anatomical details of the diaphysis. Distally, the dorsal tubercle and grooves for the extensor musculature do not appear particularly well-developed.

### The Ulnae (Table 4, Fig. 5)

#### Preservation of StW 573k (left ulna)

The left ulna is nearly complete but is missing the dorsal and anteromedial distal epiphysis, distal metaphysis and extreme distal diaphysis. The specimen is composed of five pieces glued together along tight, secure contacts. The distal fracture surfaces are ancient and now present as dry-bone, right-angle edges that have been subsequently smoothed via mechanical and diagenetic processes and are now coated with breccia and calcite. The specimen is stained with sporadic circular deposits of manganese dioxide. The ventral surface, as well as the proximal third of the dorsal surface, of the specimen, is unweathered (weathering stage 0). The rest of the dorsal surface has suffered a diagenetic loss of bone cortex, where multiple layers of lamellae were removed diagenetically in large, contiguous swaths. Similar but much smaller and localized circular incidences of lamellar loss occur sporadically across the rest of the specimen. The intact periosteal bone surface evinces no biotically induced damage.

**Figure 5.**
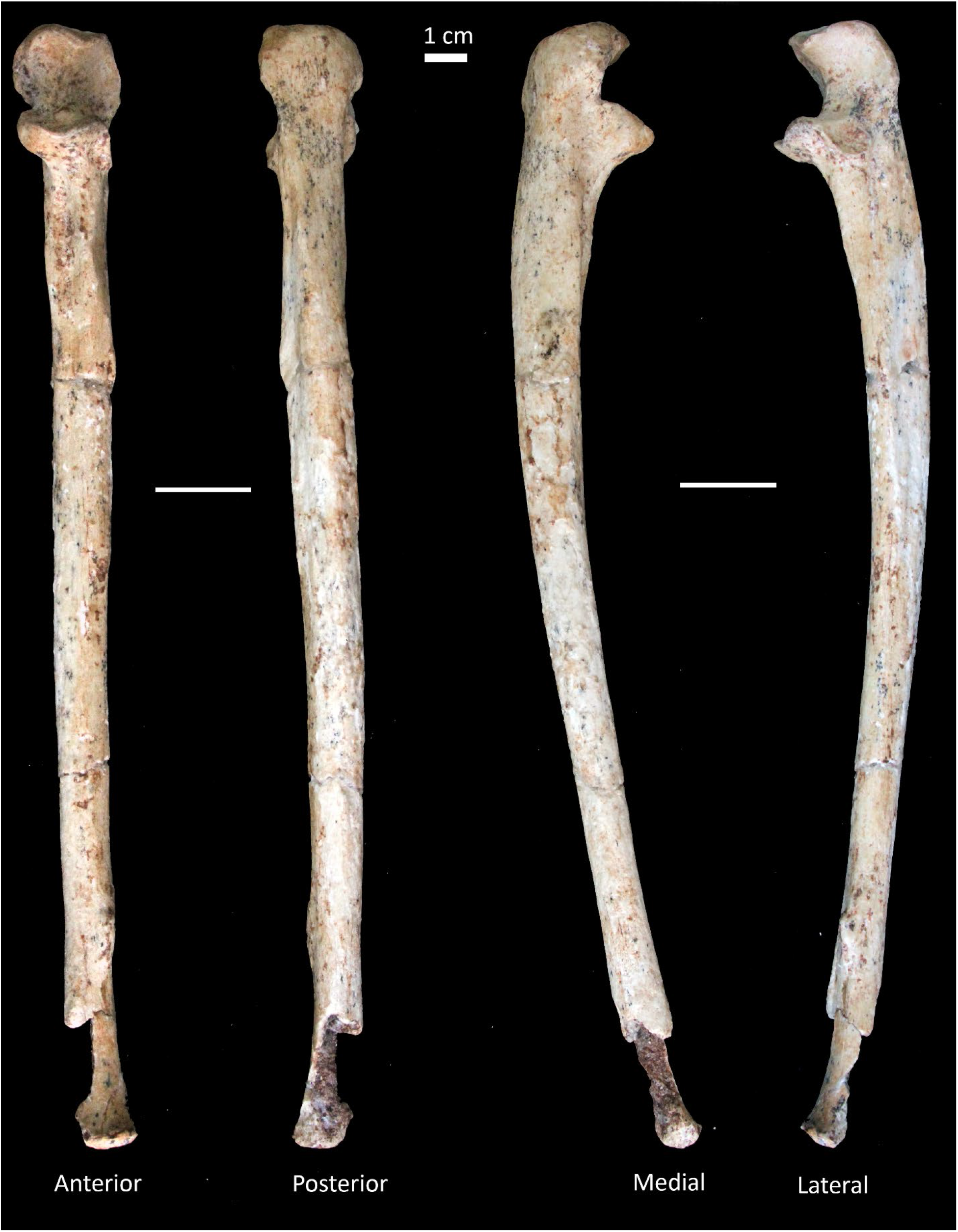
Left ulna of StW 573. Shown in anterior, posterior, medial and lateral views.

**Table 4.** Ulnar osteometrics and indices of StW 573. Estimated values are shown in square brackets.

|  | StW573k<br>(left) |
| --- | --- |
| **Raw measures (mm)** |  |
| Maximum length | 259.0 |
| Physiological length | [244.0] |
| Olecranon length | 25.5 |
| SI olecranon height | 8.4 |
| Trochlear notch height | 16.3 |
| Trochlear notch depth | 8.8 |
| SI radial notch maximum height | 9.8 |
| AP radial notch length | 13.4 |
| AP depth from coronoid tip to dorsal margin of bone | 28.0 |
| ML breadth of proximal portion of trochlear notch articular surface | 24.3 |
| ML breadth of middle portion of trochlear notch articular surface | 20.6 |
| ML breadth of trochlear notch dorsal to radial notch | 15.7 |
| SI length of articular trochlear notch medially | 22.1 |
| SI length of articular trochlear notch laterally | 22.6 |
| Distance from posterior margin of ulna to maximum depth of keel of trochlear notch | 15.7 |
| AP anconeal process | 23.0 |
| ML olecranon process maximum breadth | 26.8 |
| Minimum distance between posterior margin of ulna and articular surface of trochlear notch laterally | 9.5 |
| Mediolateral breadth of diaphysis immediately distal to radial notch | 15.5 |
| Ulnar tuberosity position | 33.6 |
| AP diaphysis diameter (proximal) | 16.2 |
| ML diaphysis diameter (proximal) | 14.3 |
| AP diaphysis diameter (midshaft) | 14.2 |
| ML diaphysis diameter (midshaft) | 12.0 |
| Diaphysis circumference (at midshaft) | 44.8 |
| Dorsal curvature chord | [224.0] |
| Dorsal curvature subtense | [12.0] |
| **Angular measures (°)** |  |
| Olecranon orientation | 118 |
| Trochlear notch proximal keel | 125 |
| Trochlear notch middle keel | 130 |
| Trochlear notch distal keel | 132 |
| Trochlear notch orientation | 8 |
| **Indices<sup>a</sup>** |  |
| Distance between middle of trochlea and middle of tuberosity taken parallel to shaft axis/physiological length | [0.14] |
| Projected distance between most proximal point on olecranon and center of trochlea taken parallel to shaft axis/physiological length | [0.7] |
| (AP anconeal process/ AP depth from coronoid tip to dorsal margin of bone) * 100 | 82.1 |
| 100 * (AP proximal diaphysis diameter * ML proximal diaphysis diameter)<sup>0.5</sup>/physiological length | [6.2] |

<sup>b</sup> Indices are described in Churchill et al. (2013).

#### Morphology of StW 573k (left ulna)

The specimen is strongly dorsally curved but is fairly straight in anterior and posterior views. Medially, the olecranon process projects only slightly superiorly above the cranial rim of the trochlear notch. A small pit near the dorsolateral corner of the proximal olecranon may be a foramen, but it is obscured by infilled breccia and calcite. The triceps brachii insertion area is broad both SI and ML but not particularly rugose, with no lipping of its borders. The trochlear notch of the ulna faces anteriorly and is keeled proximally at a sharper angle than it is in the middle or distally (Figure 6). All demifacets of the proximal joint surface complex, including the radial notch, are deeply cupped. The circumference of the trochlear notch is defined by a distinctly raised lip, which, laterally, is contiguous with the margin of the radial notch. The superomedial surface of the notch rim is particularly strongly built for the proximal reaches of the attachment of the proximal ulnar collateral ligament. Likewise, the inferomedial corner of the trochlear rim assumes the form of a large, well-developed boss that projects strongly distally and medially from the rim. There is a moderately developed gutter set dorsal to the medial rim of the trochlear notch, which provides a contiguous, SI elongated surface for the origin of the flexor digitorum superficialis proximally and that of the ulnar head of the pronator teres distally. The ulnar tuberosity emerges distally to the trochlear notch in the form of two markedly anteriorly projecting, SI elongated ridges that originate, respectively, at the bases of the inferomedial boss of the notch rim and of the coronoid process and that form a deep trough between them. The former ridge serves as the sharply elevated medial corner of the anterior aspect of the bone’s proximal metaphysis and diaphysis. The maximum SI span of the ulnar tuberosity from the base of the trochlear notch is ~32.0 mm. Like the medial and anterior portions of the proximal ulna, the proximolateral demifacet of the trochlear notch is well-developed, projecting laterally as a large, distinct flange for a SI length of ~15.5 mm, at which point it pinches medially into a short ridge, ~8.5 mm in length SI. This ridge bifurcates at its distal terminus into a weakly expressed supinator elevation anteriorly and a strongly expressed, rugose elevation posteriorly that served as the lateral boundary of the anconeus insertion. The supinator elevation sweeps anteriorly at its distal extent to connect with an underdeveloped interosseous border that originates as a short, laterally projecting enthesophytic flange ~40.0 mm inferior to the distal margin of the radial notch. The interosseous border flattens gradually from the distal base of this flange and its remainder presents as rather smooth and rounded until ~53.5 mm from the superolateral margin of the ulnar head. It is at this point that the interosseous margin rises laterally into a sharp, flange-like enthesophyte that is ~15.0 mm in SI length. There has been some loss of bone surface, but the contiguous span for the origins of the abductor pollicis longus, extensor pollicis longus and extensor indicis currently presents as a shallow, AP narrow groove along the lateral diaphysis between the interosseous border and longitudinal crest. There is significant cortical loss of the distal dorsal diaphysis, but it appears that the posterior border originally maintained its relatively steep peak for much of the length of the shaft, with perhaps slightly less distal rounding of the border than is typical of modern human ulnae. The superoanterior edge of the pronator ridge is a well-defined, SI elongated rugosity. Any potential evidence of a nutrient foramen has been obscured or obliterated.

**Figure 6.**
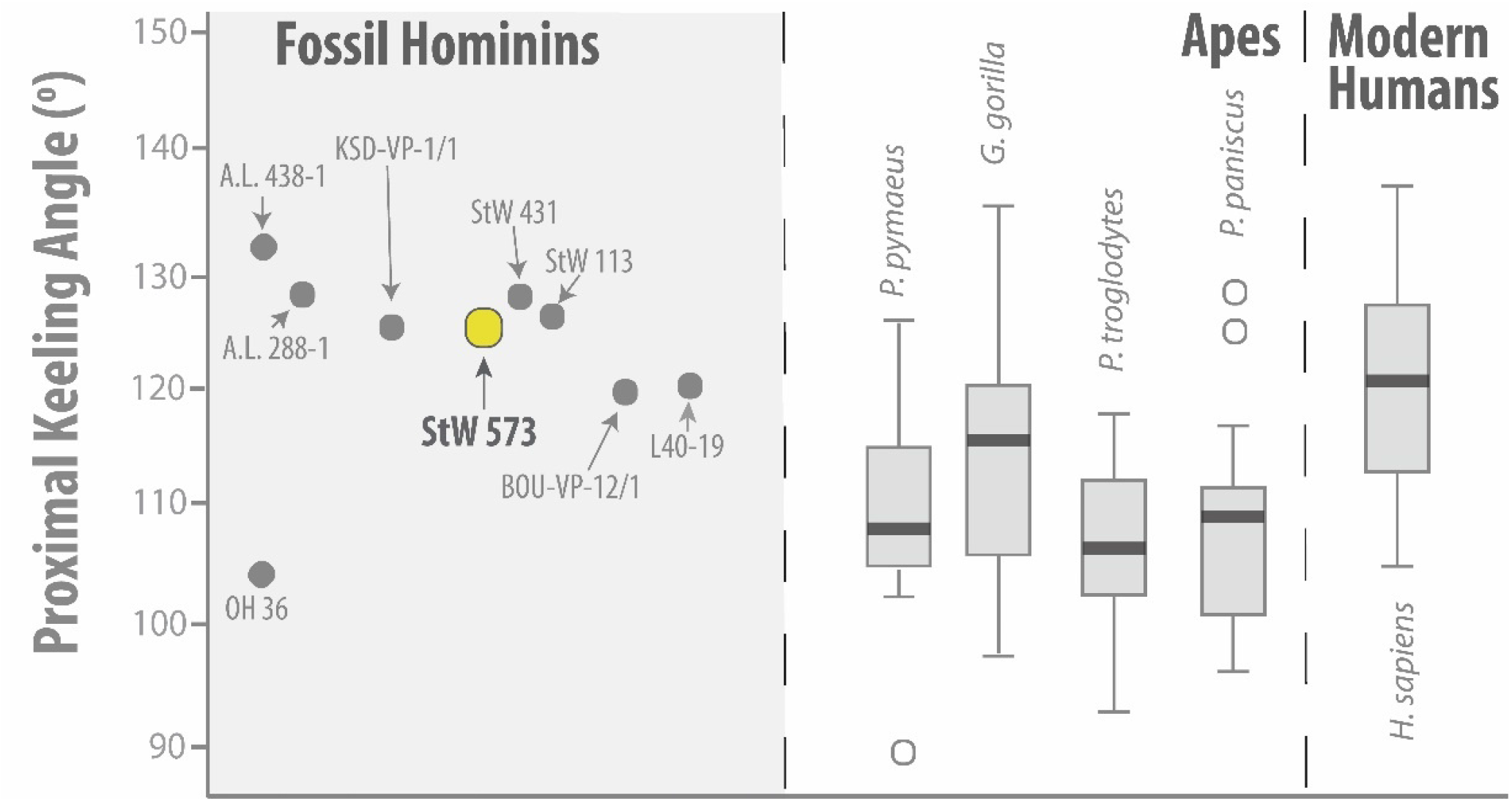
Proximal Ulnar Keeling (in degrees). Figure adapted from Drapeau (2008) where lower values indicate a proportionally more steeply keeled notch.

#### Preservation of StW 573l (right ulna)

The right ulna is incomplete, its distal +1/3 missing. The distal fracture surface is ancient, probably originally right-angled but now smooth via mechanical and diagenetic processes. The fracture surface is coated in breccia and calcite. The preserved portion of the diaphysis is crushed AP, across a collapsed medullary cavity, and bent and twisted along its SI length. The radial notch has been obliterated. Because the ulna is cemented to the right distal humerus in approximate anatomical extension at the elbow, it is difficult to ascertain, but the olecranon appears slightly splayed ML, rendering the trochlear notch and anterior olecranon invisible. Bone surface preservation across the specimen is generally good (weathering stage 0), but there are occasional patches of invasively removed cortex, especially on the lateral diaphysis and, to a lesser extent, on the dorsal and medial aspects of the shaft. The specimen shows light, sporadic deposits of breccia, calcite and manganese dioxide, but no biotically induced damage.

#### Morphology of StW 573l (right ulna)

Like the right radius, the right ulna appears to have a relatively straight diaphysis, but confirmation of that assessment awaits virtual reconstruction of the bone. Although not directly observable, the trochlear notch appears to be anteriorly directed. Dorsally, the olecranon is broad ML (~25.0 mm) (perhaps artificially; see above), with a correspondingly wide insertion surface for the triceps brachii. There is a small, proximally projecting enthesophyte set roughly in the middle of this surface. This bony projection is positioned superior to a small indention that serves as the midpoint of low-set but rugose elevation that runs nearly the transverse width of the olecranon. The coronoid process lacks appreciable anterior and superior projection, but is robust, as is the remainder of the distal rim of the trochlear notch. Posterior to the medial rim of the notch, the ulna shows a deeply cupped, SI elongated surface for the origin of the flexor digitorum superficialis superiorly and the origin of the ulnar head of the pronator teres inferiorly. This trough is situated between two especially strongly developed ridges, the anterior of which provided a surface for the attachment of the proximal ulnar collateral ligament superiorly and the insertion of the medial fibers of the brachialis inferiorly, the posterior of which provided a long surface for the proximal origin of the flexor digitorum profundus. Any potential evidence of a nutrient foramen has been obscured or obliterated.

### The Femora (Table 5, Figs. 7–10)

#### Preservation of StW 573m (left femur)

The left femur is incomplete, largely missing its neck, as well as its greater and lesser trochanters. It is preserved as two non-articulating segments: (1) a small anterior portion of the head (StW 573m/1) and attached ~31.5 mm long segment of the neck, and (2) much of the diaphysis, as well as the complete distal epiphysis (StW 573m/2). The anatomical placement of the head and neck of the left femur was achieved by comparison to its more complete antimere (StW 573n). The head fragment exhibits exposed trabeculae, and at the time of study, retained an adhering small fragment of rock near what is possibly the fovea capitis. Proximally, the medullary cavity of the complete portion of the left femur (i.e., StW 573m/2) is filled with breccia. The area near the gluteal ridge is exfoliated. Overall preservation of the left femoral diaphysis and distal epiphysis are much better than is that of its antimere. No biotically induced damage is observed.

**Figure 7.**
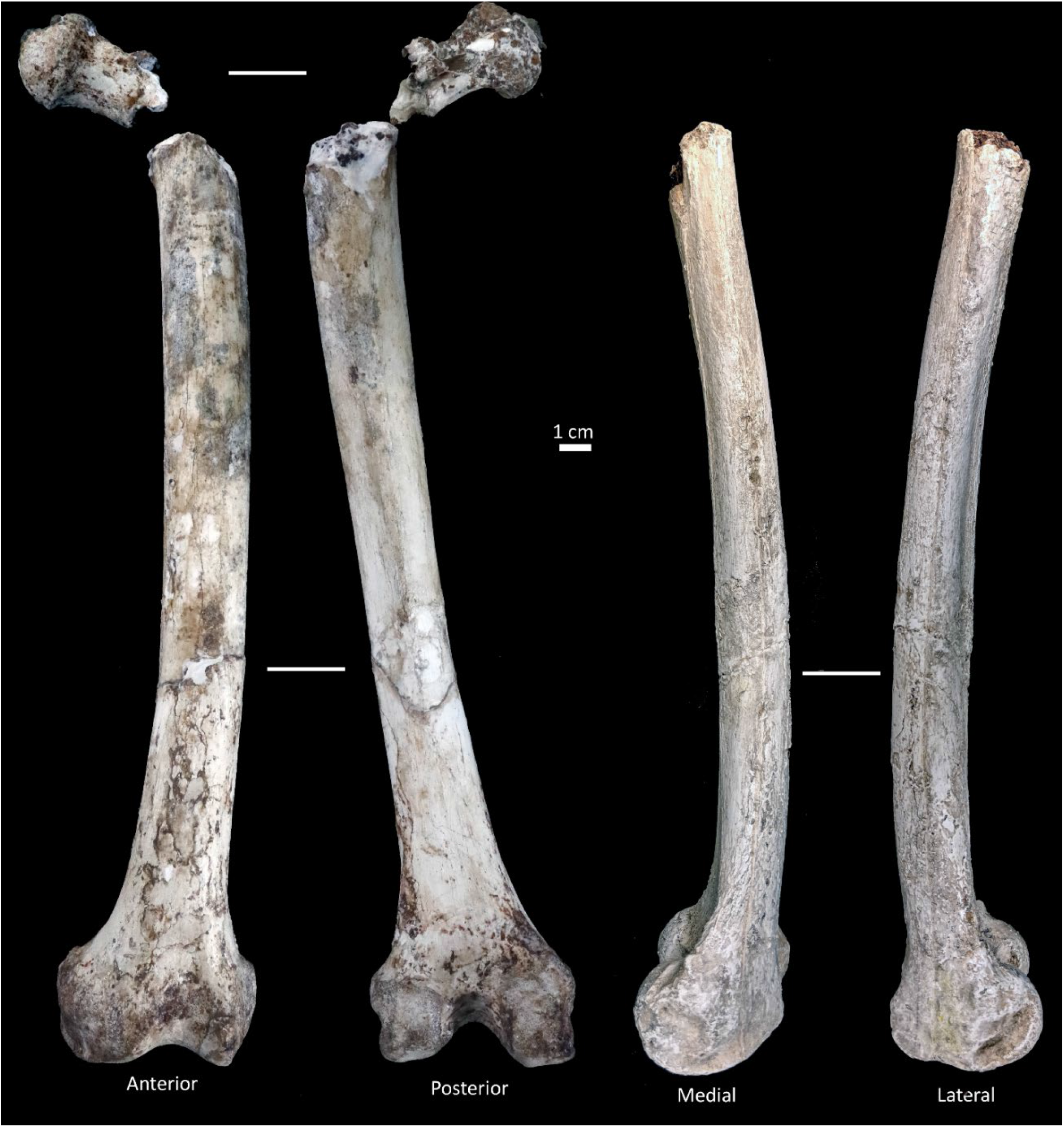
Left femur of StW 573. Shown in anterior, posterior, medial and lateral views. Medial and lateral views are shown on a cast.

**Table 5.**
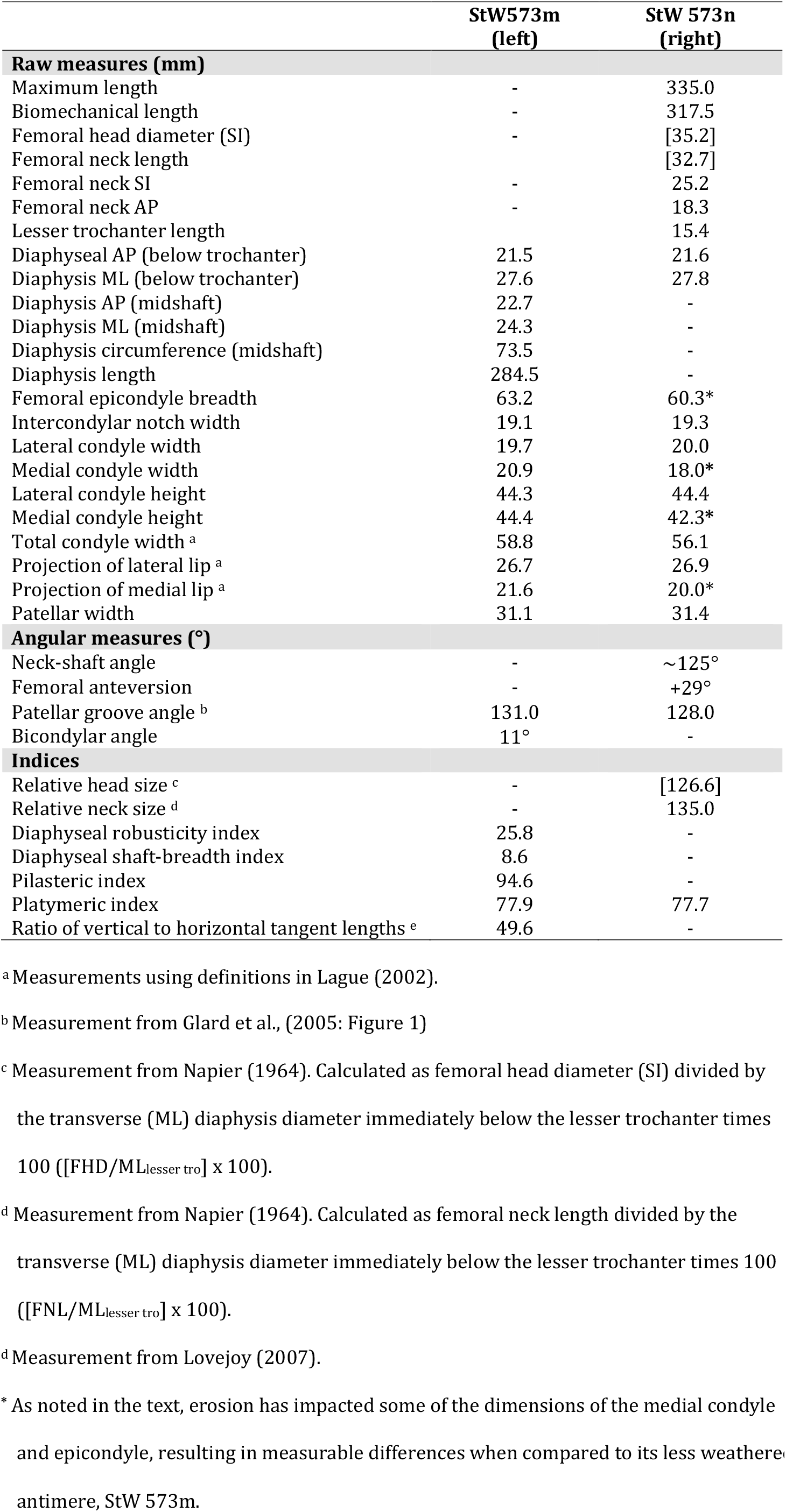
Femoral measurements and indices of StW 573. Estimated values are shown in square brackets.

#### Morphology of StW 573m (left femur)

The small head and neck fragment of the left femur has a maximum length of 51.5 mm. The shaft exhibits some proximal AP flattening, as indicated by the platymeric index (77.9) taken immediately inferior the lesser trochanter (AP=21.5 mm, ML=27.6 mm). The midshaft also exhibits AP flattening; however, it is less apparent due to the presence of a moderately developed pilaster. Due to its missing proximal segments, the anteroposterior curvature of the shaft (Martin and Saller, 1957) could not be computed, but we qualitatively estimate that the shaft (viewed laterally) possesses minimal anteroposterior bowing. However, the shaft does appear to bow laterally (anterior view) and anteriorly (medial view). The robusticity index of the diaphysis is 25.8 (midshaft circumference=73.5 mm, diaphysis length=284.5 mm) with a shaft-breadth index of 8.6 (ML diaphysis at midshaft=24.4 mm, diaphysis length=284.5 mm). Moving distally, a faint gluteal ridge is apparent on the posterolateral edge of the diaphysis. Between this ridge and the lateral border of the femur, a narrow and linear hypotrochanteric fossa is observed. The diaphysis also exhibits a moderately-developed pilaster (index = 93.0). The linea aspera is well-developed and most readily distinguished near the midshaft. The medial supracondylar line, insertion for the biceps femoris (not observable on the antimere of StW 573m) shows a distinct adductor tubercle but is itself fainter than is the lateral supracondylar line (origin for vastus lateralis). Assessed qualitatively, the muscle markings are more well-developed than observed in an available comparative *Pan* femur. A moderately elevated area for the attachment of the medial collateral ligament is preserved on the medial epicondyle. Posteriorly, the popliteal surface is a wide, flat, triangular area. The lateral epicondyle is convex preserving the facet for the gastrocnemius, as well as discernible impressions for the popliteus and its tendon, posterolaterally. Compared to the lateral condyle, the medial is larger in width but similar in height. In overall shape, the differences between the condyles appear larger due to the anterior shift of the lateral condyle, as well as its well-developed lip. However, both condyles possess elliptical profiles when viewed from the side. In distal view, the condyles of the left femur appear to approximate the square outline observed in modern humans, which are roughly symmetrical around the patellar groove. The superior border of the patellar surface is relatively flat but possesses a well-defined sustrochlear hollow (Tardieu, 2010), more clearly visible here than on the right femur. Additionally, the specimen possesses a deep patellar groove with well-marked medial and lateral margins. The lateral patellar margin forms an elevated lip and projects more anteriorly than the medial. The intercondylar notch and line are well-preserved and intact. Measurable in this femur, StW 573 possesses a human-like bicondylar angle (11°). Cruciate ligament sites are not readily visible on this femur.

#### Preservation of StW 573n (right femur)

As the proximal and distal portions of the femora were separated due to the faulting (Clarke, 2018), a large gap opened within the breccia in which the StW 573 was encased. Subsequently, a flowstone filled in the region between the two right femoral fragments: the right head, neck and proximal diaphysis (StW 573n/1) and the distal diaphysis and epiphysis (StW 573n/2). Although largely cleaned away, a calcite infilling still adheres along the fracture edges of both fragments. In these areas, the cortical bone is now thin and delicate, with a significant lamellar loss. Thus, any attempt to further clean and articulate the two fragments was deemed too perilous. Consequently, the two pieces of StW 573n do not cleanly join. Fortunately, the left femur of StW 573 preserves the missing region of the distal diaphysis of the right. Using features, such as portions of the linea aspera and pectineal and gluteal lines, the femora can be aligned to provide a relatively complete picture of its femoral morphology. Proximally, the right femur has suffered crushing damage. The anterior and posterior regions of the head have been pinched together, which has forced an inferior portion of the head to be deflected inferolaterally, while the superoposterior rim appears less damaged and in its original location. Crushing and oblique fractures along the neck, as well as similar damage to the area extending inferior to the lesser trochanter, seem to only minimally impact the proximal femur’s overall shape and dimensions. The greater trochanter of the specimen is heavily damaged and preserves only a small segment of its anterior and lateral regions. Largely present, the lesser trochanter shows only minimal cortical damage. The diaphysis has been heavily damaged diagenetically. The area surrounding the femoral condyles and epicondyles are preserved; however, the area is in poorer condition than that of its antimere. Due to diagenetic changes of its medial epicondyle, the femoral epicondyle breadth is smaller for the right femur than the better-preserved left. Measures of the right distal femur should be treated with caution, as it has suffered more cortical damage and erosion. However, values are reported here, when measurable. The medial patellar margin is slightly, but not significantly, impacted by the weathering and damage that has impacted much of the medial distal end. A small stone and damage obstruct the lateral groove while the medial is minimally visible. The medial half of the epiphysis has suffered considerable cortical damage. No evidence of biotic damage is observed.

#### Morphology of StW 573n (right femur)

Because proximal damage, a femoral head SI measurement is not directly measurable but can be estimated in accounting for the inferior damage. A measurement (35.0 mm) along the well-preserved edges, superoposterior and inferoanterior, of the femoral head (~35° from SI vertical) is similar to the SI estimated measure (35.2 mm) using Image J. This latter measurement was obtained using ImageJ and accounting for the 1-2 mm inferomedial deflection of bone. CT reconstructions are currently in process and will refute or confirm our estimation. However, we do not expect it to be more than 1 mm different from the estimated value provided here. The maximum femoral length of the right femur was estimated by aligning overlapping areas of both femora. Viewed superiorly, the angulation of head is positioned so that the head is placed slightly anteriorly onto the neck. Based upon a small preserved portion of the intertrochanteric crest, the femoral neck is comparatively long (Figure 9) and flattened anterio-posteriorly (Figure 10). The neck-shaft angle (~125°) is within modern human ranges and slightly outside of African ape values (see Napier, 1964). While fragmentary, the most inferior portion of the greater trochanter flares laterally. However, this observation should be treated with caution, as the area is filled with bone-reinforcing plaster. And, the impact of damage, or flowstone intrusion, of this flaring cannot confidently be determined. The lesser trochanter, serving as the insertion of the iliopsoas (major flexor of the thigh) is small (SI=15.4mm), but medially directed. Connecting the remnants of the intertrochanteric line with the linea aspera, the spiral line – the origin of the vastus medialis – is moderately developed. The pectineal line cannot be assessed, as the area is damaged on the right femur. Neither the obturator externus groove nor the trochanteric fossa is clearly visible; only a small, uninformative portion of the latter is preserved. The platymeric index of StW 573m is 77.9 (immediately inferior to lesser trochanter, AP=21.5 mm, ML=27.6 mm). The robusticity index of the diaphysis is 25.8 (midshaft circumference=73.5 mm, diaphysis length=284.5 mm) with a shaft-breadth index of 8.6 (ML diaphysis at midshaft=24.4 mm, diaphysis length=284.5 mm). The remaining features of the shaft are better observed in the left femur. The medial condyle is smaller in width and height than is the lateral condyle.

**Figure 8.**
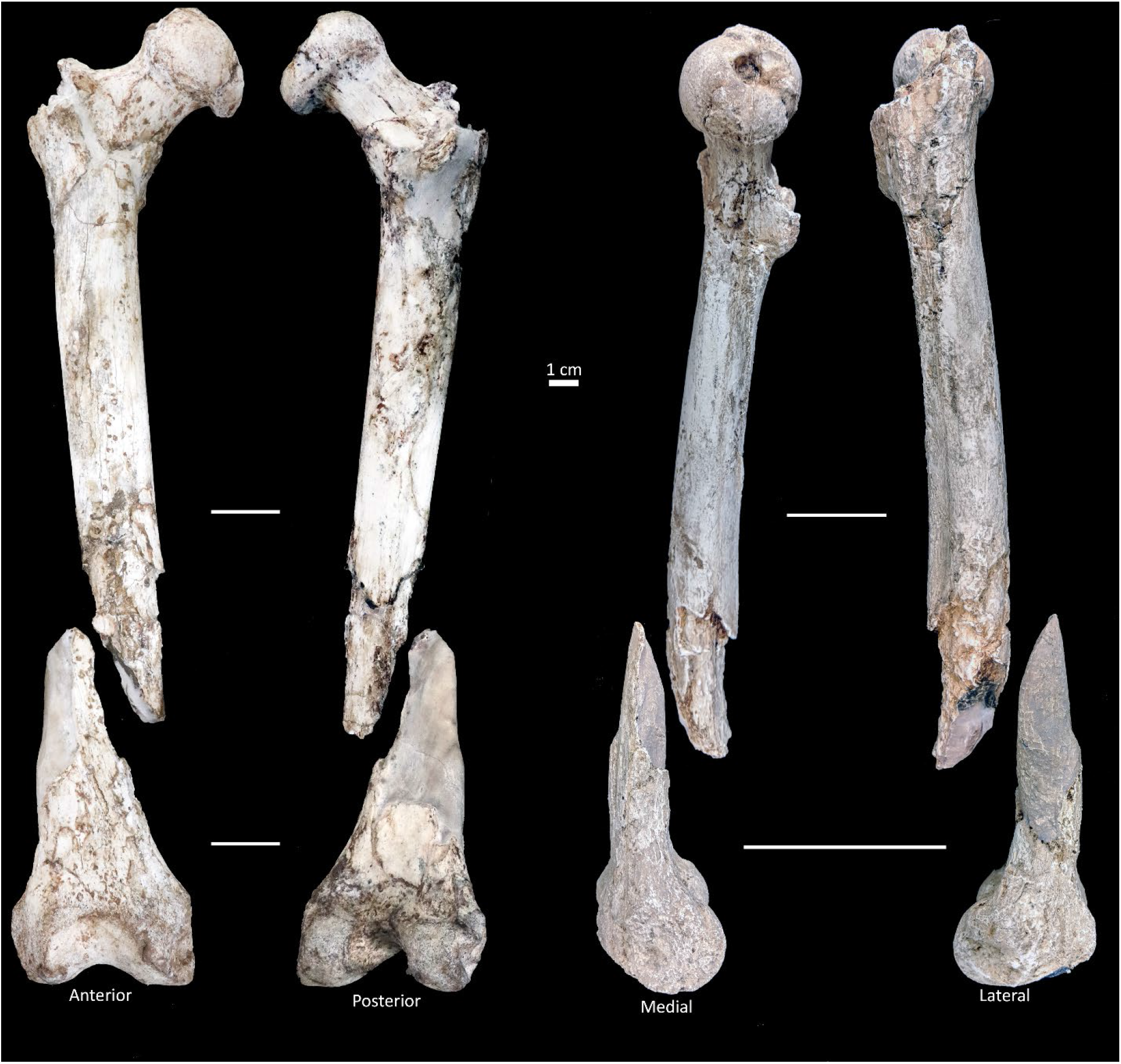
Right femur of StW 573. Shown in anterior, posterior, medial and lateral views. Medial and lateral views are shown on a cast.

**Figure 9.**
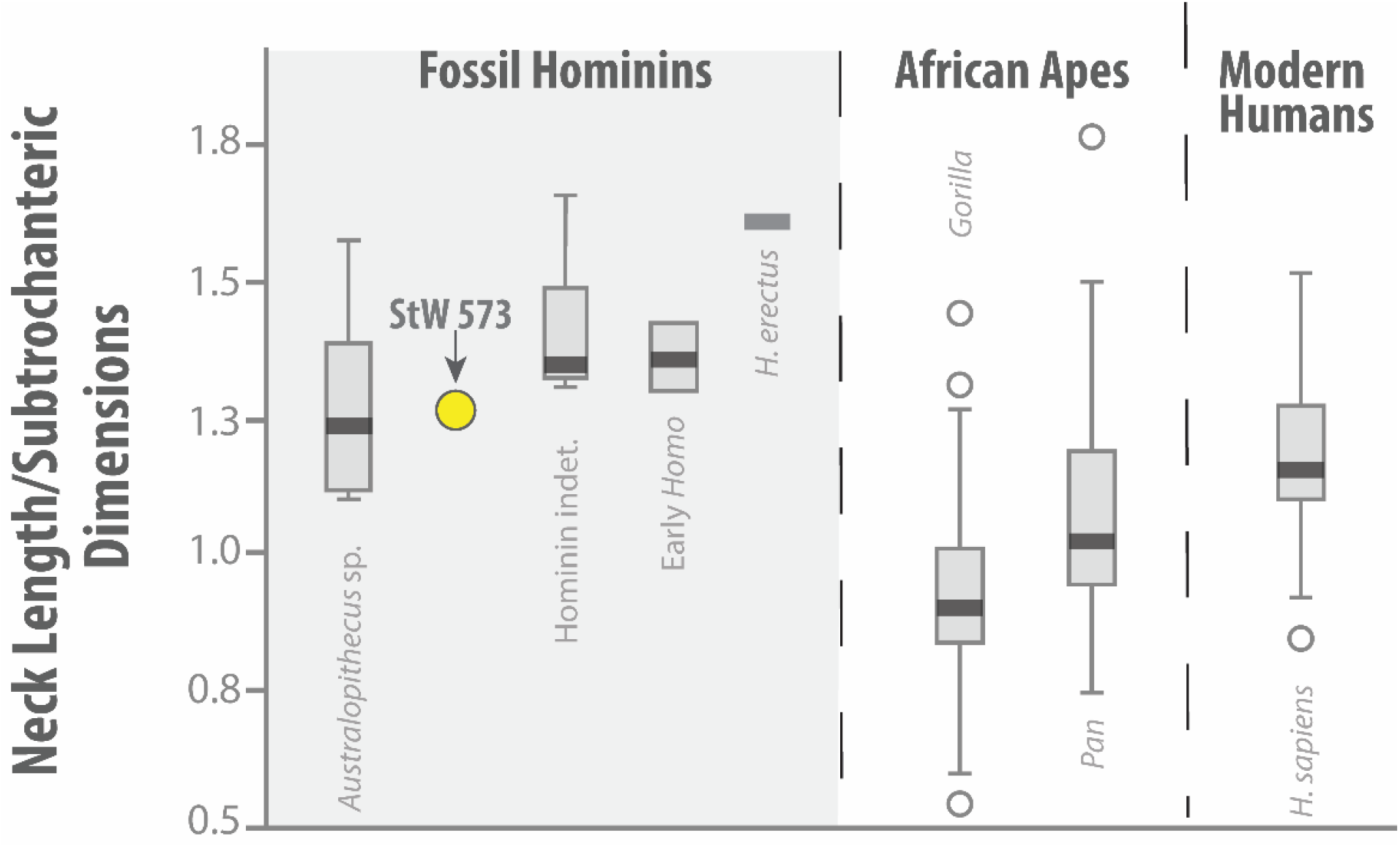
Relative neck length. Shown are the femoral neck lengths (relative) for African apes and fossil hominins. StW 573’s value was calculated following and figure adapted from Marchi et al.’s (2017). Box-and-whisker plots denote the mean (dark horizontal line), upper and lower quartiles (boxes), range (whiskers) and outliers (open circles).

**Figure 10.**
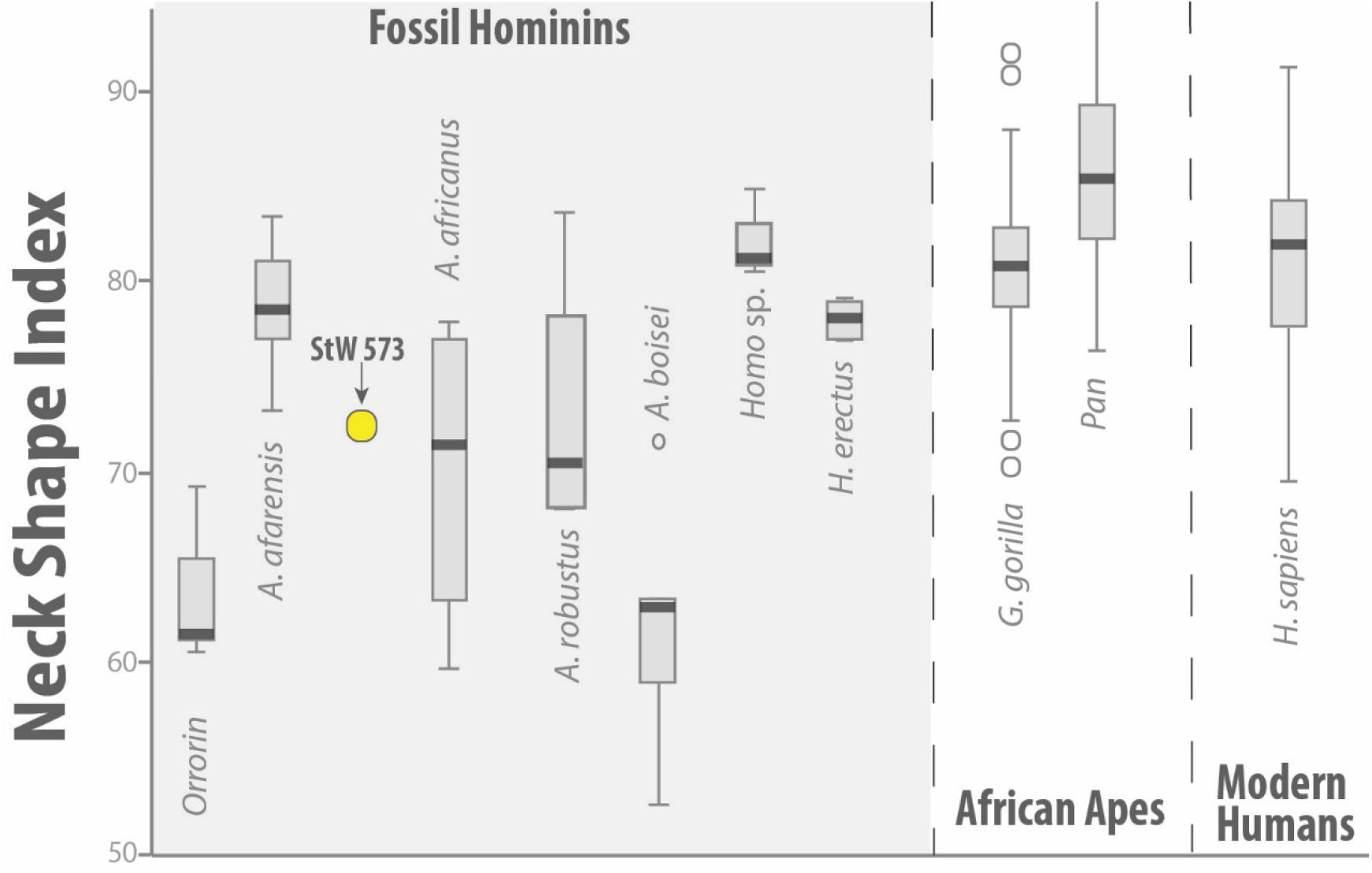
Neck Shape Index. Relative shape of the femoral neck (AP*100/SI) shown in fossil hominins, African apes, and modern humans. Box-and-whisker plots denote the mean (dark horizontal line), upper and lower quartiles (boxes), range (whiskers) and outliers (open circles). Measurements for fossil hominins derive from Grabowski et al. (2015: Supplemental Online Material and sources cited, therein). In this analysis, taxonomic assignments in the original sources were maintained. However, as there is some debate on the number of species at Sterkfontein *A. africanus* may include specimens attributed to *A. prometheus* (for discussion, see Clarke, 2013; Clarke and Kuman, 2018) and whether some specimens should be included within *H. habilis* (Clarke, 2017).

The specimen shows a deep patellar groove, with a well-developed lateral lip. The patellar surface exhibits a relatively flat superior (proximal) surface. The intercondylar line and notch are reasonably intact and undamaged. Of the two femora, the attachment sites for the cruciate ligaments are best viewed, here. The area of attachment for the posterior cruciate ligament is relatively flat, while the anterior cruciate ligament site exhibits a slightly elevated ridge.

### The Tibiae (Table 6, Figs. 11–12)

#### Preservation of StW 573o (left tibia)

The left tibia is preserved as two separate fragments: the proximal epiphysis along with approximately 2/3 of the diaphysis (StW 573o/1) and the distal 1/3 of the diaphysis, as well as the medial malleolus and talocrural joint (StW 573o/2). Of the two tibiae, the left is in an overall poorer condition due to fragmentation of its shaft. The diaphysis is relatively lightly covered with breccia. Immediately distal to the tuberosity, small cracks (<1 mm) frequently occur along the proximal and mid-diaphysis. In some cases, the cracks surround large islands of cortical bone with one of the largest (~38 mm x16 mm fragment) being placed proximal to mid-diaphysis. In many cases, breccia has filled in these small cracks. Larger (> 1 mm) cracks more frequently occur in the distal half of the diaphysis, as they approach the fracture between StW 573o/1 and StW 573o/2. The latter was blasted from the remainder of the skeleton (along with the medial malleolus of StW 573p) during lime-mining activity within the Silberberg Grotto (Members 2 and 3). The two left tibial fragments of StW 573 were joined along a perfect contact in the core of the shaft, and only the cortical surfaces of the lower tibial fragment had been blasted away during lime mining. This distal fragment (StW 573o/2) is lighter in color due to blasting damage, which has also removed any breccia that most of the excavated bones exhibit. Much of the cortical surface has been blasted away superiorly. A transverse fracture is observed near the talar surface and travels through the medial malleolus.

**Figure 11.**
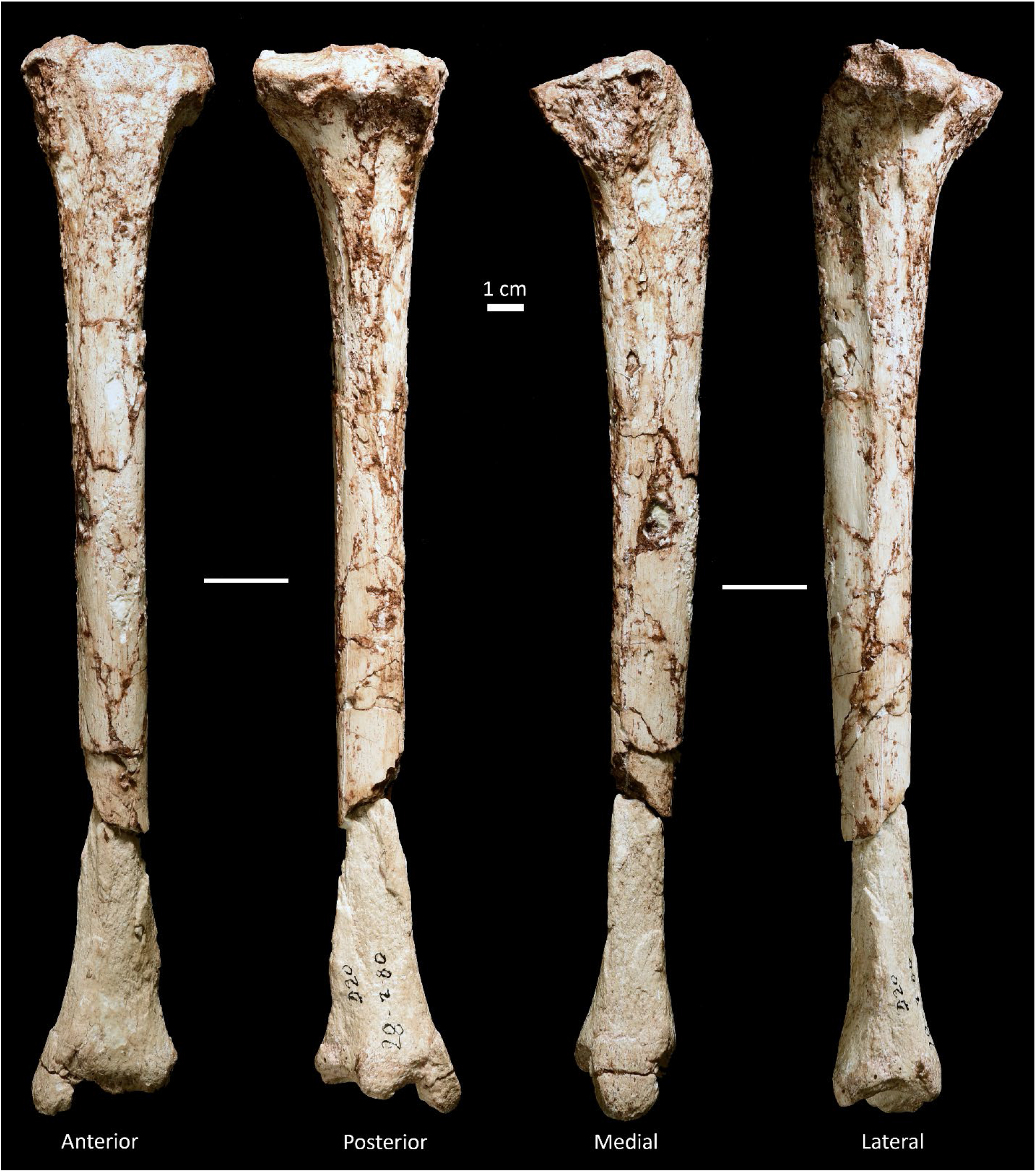
Left tibia of StW 573. Shown in anterior, posterior, medial and lateral views.

**Figure 12.**
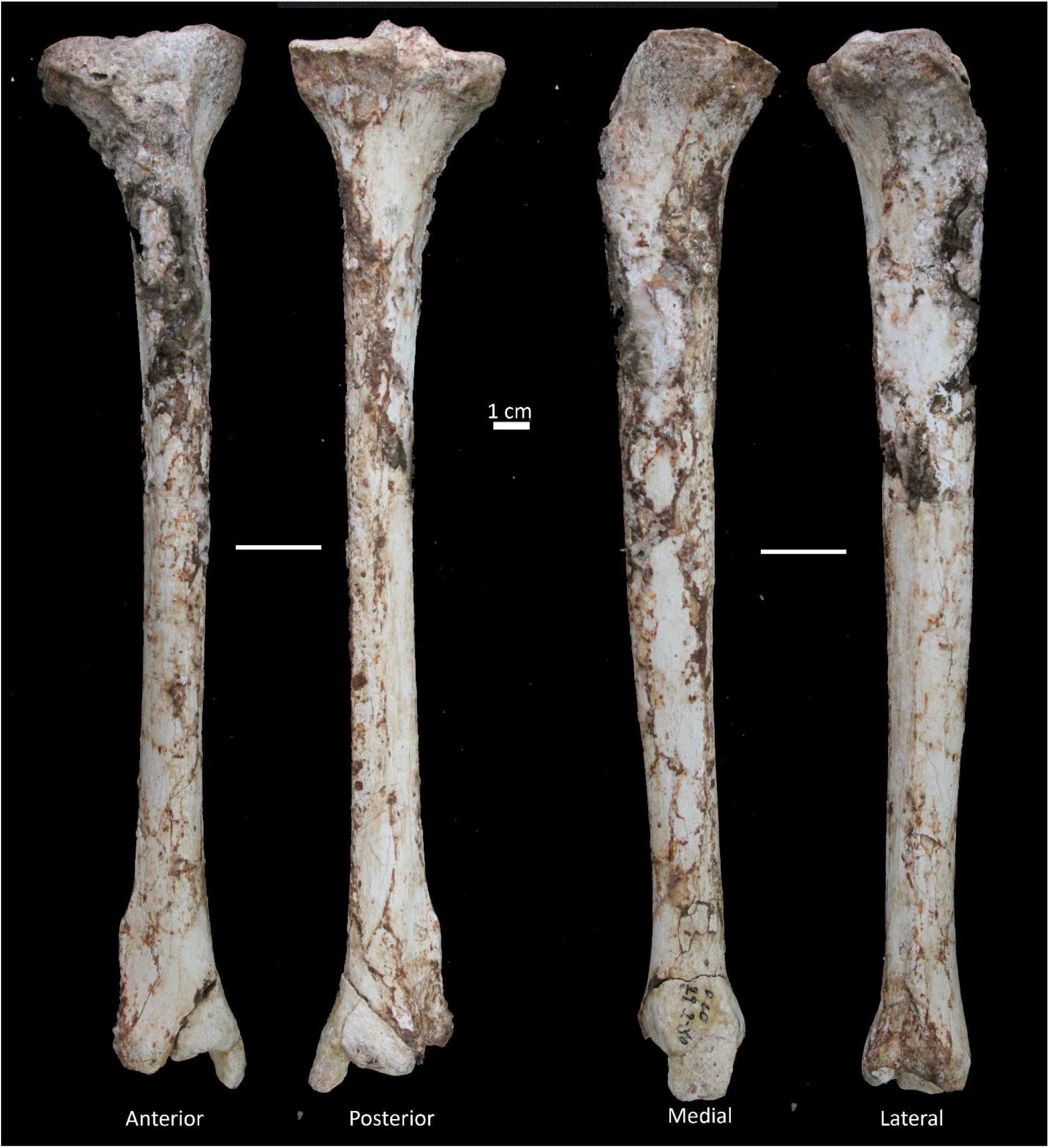
Right tibia of StW 573. Shown in anterior, posterior, medial and lateral views.

**Table 6.** Tibial osteometrics and indices of StW 573. Estimated values are shown in square brackets.

|  | StW573o<br>(left) | StW 573p<br>(right) |
| --- | --- | --- |
| **Raw measures (mm)** |  |  |
| Maximum length | - | 285.0 |
| Biomechanical length | - | 273.0 |
| Interspinal distance | 8.7 | 8.9 |
| AP plateau depth | [38.0] | 38.3 |
| ML plateau breadth | [58.6] | 57.1 |
| AP medial condyle depth | 36.8 | 36.7 |
| ML medial condyle width | 24.0 | 23.5 |
| AP lateral condyle depth | 31.2 | 31.5 |
| ML lateral condyle width | 24.3 | 22.4 |
| AP diaphysis diameter (at nutrient foramen) | 28.9 | 28.7 |
| ML diaphysis diameter (at nutrient foramen) | [20.3] | 18.5 |
| AP diaphysis diameter (midshaft) | 23.3 | 23.4 |
| ML diaphysis diameter (midshaft) | [19.3] | 18.3 |
| Diaphysis circumference (at nutrient foramen) |  | 73.0 |
| Diaphysis circumference (at midshaft) | [66.0] | 67.0 |
| AP depth of distal end (maximum) | - | 30.7 |
| ML breadth of distal end (maximum) | - | 33.2 |
| ML anterior talar surface | 26.1 | 26.6 |
| ML midpoint talar surface | 21.6 | 19.6 |
| ML posterior talar surface | 18.0 | [17.8] |
| AP lateral talar surface | 24.0 | 25.2 |
| AP midpoint talar surface | 21.7 | 23.1 |
| AP medial talar surface | 19.5 | [18.9] |
| Distal projection of medial malleolus <sup>a</sup> | 11.8 | 11.1 |
| **Angular measures (°)** |  |  |
| Retroversion angle | [110°] | 110° |
| Inclination angle | [109°] | 109° |
| Tibial arch angle | +3.8° | [+3.9°] |
| Angle between tibial diaphysis and tibiotalar joint | [90.2°] | - |
| **Indices** |  |  |
| Intercondylar space ratio <sup>b</sup> | 44.2 | 46.1 |
| Platycnemia index | [70.2] | 64.5 |
| Robusticity Index <sup>c</sup> | - | 14.6 |

<sup>a</sup> Measured after Tallman et al. (2013).
<sup>b</sup> Calculated by $[100 \times \text{Interspinal distance (of tibia)}] / \text{Intercondylar notch width (of femur)}$
<sup>c</sup> Robusticity Index = (Maximum + Minimum diaphyseal breadth/bone length) \* 100; (Stock and Shaw, 2007)

#### Morphology of StW 573o (left tibia)

The oval tibial plateau is wider ML than deep AP. The medial condyle is roughly triangular while the lateral is sub-circular; both are concave. The medial condyle is more extensive in all dimensions when compared to the lateral. The latter appears to possess a single attachment point for the lateral meniscus. The lateral condyle is also un-notched and not markedly convex along its posterior border. Of the two tubercles, the medial intercondylar eminence is taller and more steeply inclined. The tibial diaphysis is larger near the nutrient foramen than at the midshaft. Because the diaphysis of the left tibia is more poorly preserved than its antimere, we have refrained from further detailed descriptions, here. The tibiotalar surface is approximately trapezoidal in shape (when viewed inferiorly) with its anterior border longer ML than its posterior. A slight keel is observed along the tibiotalar surface with a posterior swelling that is much larger than the smaller and beak-like anterior margin of the facet. The maximum inferior projection of the medial malleolus is slightly posterior to the diaphyseal long axis. The medial malleolus (ML = 12.5 mm, AP = 19.0 mm) is comparatively thinner^4^ than that of apes (ML/AP*100) and at the higher range of modern humans.

We refer the reader to the section below on the morphology of right tibia.

#### Preservation of StW 573p (right tibia)

The diaphysis is largely intact but is impacted by distal exfoliation and fine longitudinal cracking of the cortical bone. The proximal half of the right tibia is covered with remnants of breccia and calcite. The tibial condyles and plateau are largely unaffected by adhering matrix. However, the breccia significantly obscures much of the anatomy near the tibial tuberosity, as disconnected patches around the proximal circumference of the diaphysis. The lateral and posterior (proximal) margins are less obscured by breccia. Because of the adhering matrix, the diaphyseal measurements near the nutrient foramen may be slightly larger than that reflected by the true anatomy. A very small fragment of cortical bone is missing distomedially (~0.5 mm). The distal end of the tibia had been blasted off during lime mining and was recovered in 1980 during processing of breccia from the rubble on the floor of the Silberberg Grotto. It has been re-joined perfectly to the distal diaphysis. Proximal to the medial malleolus, blasting damage has removed an irregular patch (2cm^2^) of cortical bone near the fracture caused by blasting. This small (~1 mm wide) fracture originates proximal to the medial malleolus and courses obliquely through the middle of the talocrural joint.

Originally placed among the non-hominin materials curated at Sterkfontein, this medial malleolus fragment had been recovered earlier (on February 2, 1980) during sorting of the Silberberg Grotto miner’s dump (Dump 20). The fragment then sat in a box for nearly 20 years before being recognized, as hominin (Clarke, 1998). The break between the medial malleolus and the remainder of the tibia was fresh, not prehistoric, and the result of blasting in the cave decades before by lime-miners. Perhaps due to further exposure to the elements (e.g., being placed in a lime dump) or through differential preservation treatment, the malleolus fragment exhibits a lighter color than the remainder of the right tibia. And while the fracture travels through the talocrural joint, it does not appear to have impacted the dimensions or morphology of the joint significantly. However, the form of the anterior groove cannot be determined, as this region of the anterior tibial surface is largely missing. The distal fibular articular surface is partially observed as a small crescent facet.

#### Morphology of StW 573p (right tibia)

As reconstructed, the maximum tibial length can be confidently assessed by measurement. The tibial plateau of the right tibia is roughly oval and possesses an un-notched lateral condyle. Both condyles are distinctly concave across their articular surfaces; the lateral less so than the medial. The medial condyle is roughly triangular, with its apex directed medially and its base adjacent to the medial intercondylar tubercle. For the medial condyles, the greatest depth of concavity is situated slightly anteriorly. In contrast, the lateral condyle is sub-circular with a more centrally located concavity. In both ML and AP dimensions, the medial condyle is much larger than is the lateral condyle. The posterior border of the lateral condyle shows a slight convex edge. Additionally, the lateral condyle appears to possess a single attachment point for the lateral meniscus, and a posterior border that is short and continuous. Relative to the plateau, the tibial diaphysis appears fairly straight. The intercondylar eminence is preserved in its near entirety. The medial tubercle of the eminence is taller and more steeply inclined than the lateral, and as a result, the eminence appears to distally slope as it travels laterally. The line connecting the intercondylar eminences is approximately 20° from the ML plane. The sites of attachment for the anterior and posterior cruciate ligaments were not confidently observed. Inferior to the medial condyle, the origin for the semimembranosus is preserved as an elongated hollow on the posteromedial side. The resulting groove is wider AP (18.7 mm) than SI (11.4 mm) and exhibits a modern humanlike taper. Proximally, the tibia possesses a narrow, and shelf-like, lateral epicondyle while medially, the epicondyle is more gradual in merging with the diaphysis – a more human-like condition. The medial edge of the tibial diaphysis is rather straight. Viewed anteriorly, the diaphysis appears laterally concave (proximally) and laterally convex (distally). As a result, proximal (knee) and distal (talocrural) articulations are not exactly parallel to one another. The tibial diaphysis of StW 573 is straight in all remaining dimensions. The proximal fibular facet is directly posteroinferior to the lateral condyle and appears to merge with the round edge of the lateral condyle. The tibial tuberosity is smooth and wide at its most superior position. The inferior-most portion of the tuberosity is large and rugose; it is like modern humans in its degree of rugosity. But, distally, the tuberosity tapers much more gently. The tibial circumference is larger at the nutrient foramen than it is at the midshaft. This is also true of the AP and ML dimensions of the diaphysis. The tibial diaphysis appears to be markedly flattened but is mesocnemic (between 63.0-69.9, as defined by Bass, 2004). Proximally, the anterior crest is covered with breccia and therefore, poorly defined. However, the crest appears round in its preserved portions. The lateral surface preserves portions of the interosseous border; however, over half is partially obscured by adhering matrix. The most proximal observable portion (~15.0 mm inferior to the tuberosity) of the border would have ended near the anterior intersection of the tuberosity and the tibial plateau. Its distal course travels postero-obliquely. There is no discernible soleal line, suggesting that the soleus may attach to the fibula.

Together, these features, interosseous border, and soleal line, may indicate ape-like positions of the tibialis anterior/posterior and soleus, respectively (Aiello and Dean, 1990). The groove for gracilis is well-developed. In cross-section, the diaphysis varies from a triangle with rounded apices (proximally) to an oval with a faint interosseous border (midshaft) to a square with a laterally extremely well-developed tibiofibular ligament ridge in its distal part. Medially, the area of attachment of the flexor digitorum longus is also well-developed. The medial border possesses a well-marked ridge along its distal half for the deep transverse fascia. Just inferior to the tibial tuberosity, the medial and lateral surfaces appear to be similar in size. However, it is difficult to estimate because so much of the surface is covered by calcite and breccia. The posterior surface is ML narrow (proximally) and broad and flat (distally). Partially obscured, but in its visible portions, the posterior surface does not appear to possess a distinct soleal line, proximally. The diaphyseal nutrient foramen is not clearly visible. However, an estimation was made on its placement, and most measures (e.g., AP, ML, and circumference) varied little (<1.5 mm) based upon placement near the hypothesized foramen. The anterior colliculus of the medial malleolus is largely intact. However, the posterior colliculus is damaged by a fracture that travels obliquely (SI) which has removed most of it. There is no evidence of a posterior groove. The fibular notch is well-preserved and is bounded, anteriorly, by a large crest for the tibiofibular ligament. Because of the crest, the fibular notch looks large. Cautiously, the tibial arch angle has been estimated and indicates an anteriorly directed set of the ankle. This measure may require revision when virtual reconstructions are completed. The tibiotalar surface is roughly orthogonal to the long axis of the tibial diaphysis and appears approximately trapezoidal in shape (when viewed inferiorly). The tibiotalar surface also appears to possess a slight keel dividing its surface into medial and lateral portions; however, damage in the area has reduced our ability to comment further on its true form confidently. There is evidence of a slight posterior swelling associated with the talar facet keel. The distal end of the tibia exhibits a moderate degree of lateral torsion relative to the tibial plateau. Even so, more than the left, the medial malleolus (ML = 10.6 mm, AP = 18.8 mm) is thin; however, this measure should be treated with caution, as it possibly reflects an underestimation of the ML dimension due to bone fragmentation.

#### The Fibulae (Table 7, Figs. 13–14)

##### Preservation of StW 573q (left fibula)

The left fibula is preserved as two fragments: the left head and diaphysis (StW 573q/1) and the left lateral malleolus (StW 573q/2). The latter was one of the original fossil elements that led to the recovery of the complete skeleton (Clarke, 1998) and has been described elsewhere (Deloison, 2003). The proximal fragment (StW 573q/1) displays a poorer cortical surface along its anterior and medial margins while the distal end (of the proximal fragment) is covered with breccia with a small embedded stone. The distal fragment (StW 573q/2) is a comparatively well-preserved lateral malleolus different in coloration from the proximal fragment which was recovered in situ. There is no observed biotic damage.

**Figure 13.**
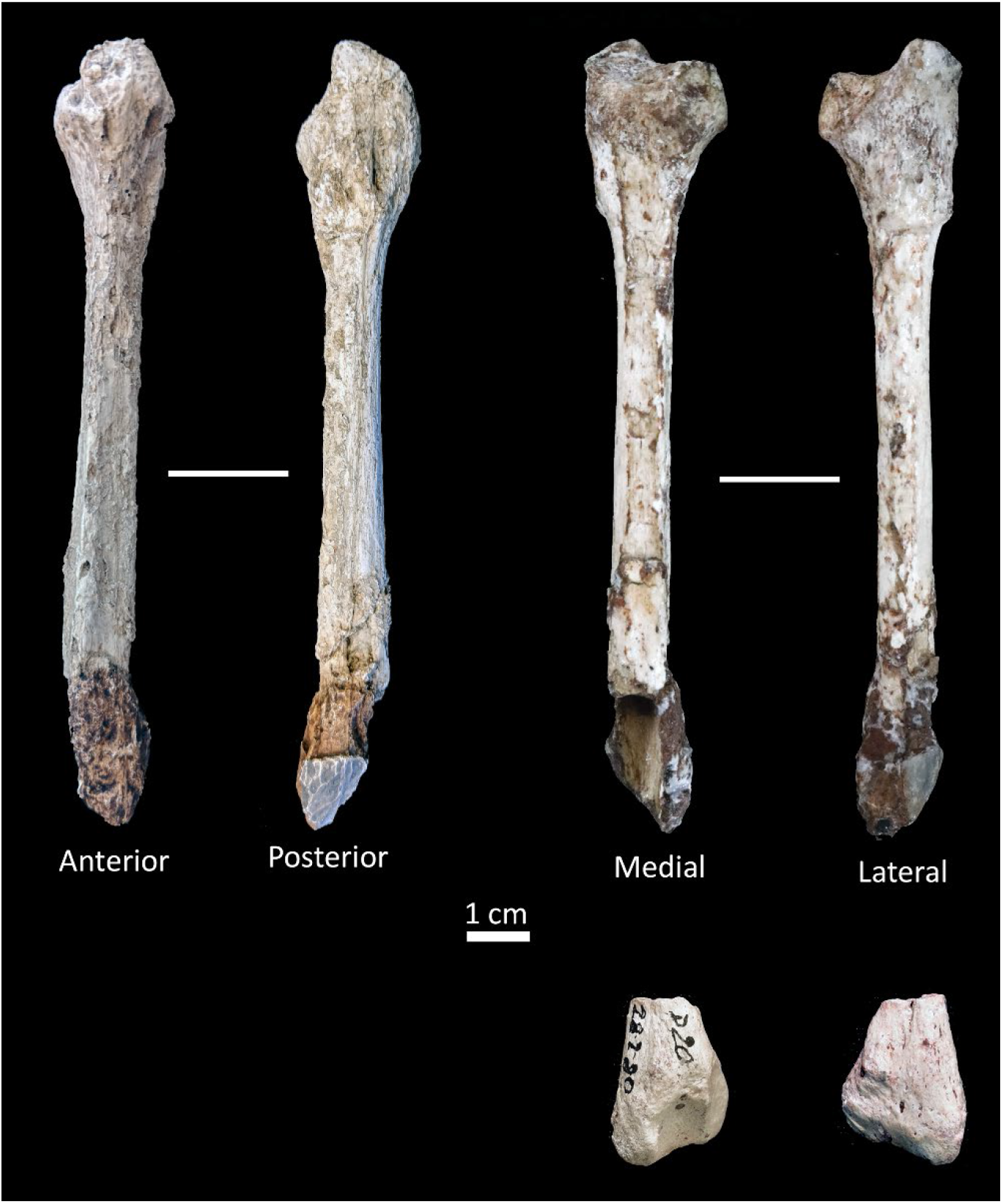
Left Fibula of StW 573. Shown in anterior, posterior, medial and lateral views. Note, anterior and posterior views are not shown.

**Figure 14.**
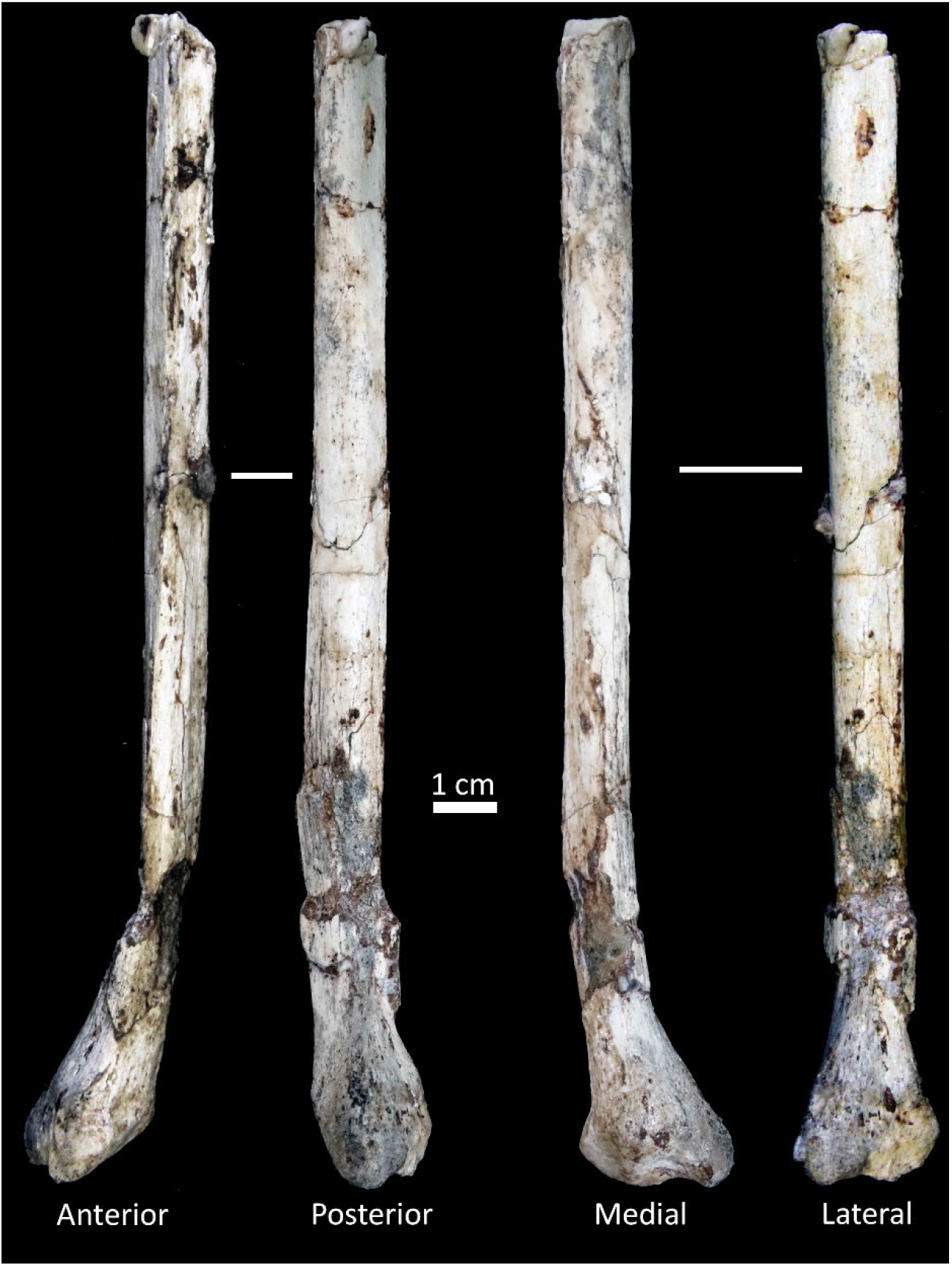
Right fibula of StW 573. Shown in anterior, posterior, medial and lateral views.

**Table 7.** Fibular measurements. Estimated values are shown in square brackets

|  | StW573q<br>(left) | StW 573r<br>(right) |
| --- | --- | --- |
| **Raw Measures (mm)** |  |  |
| Neck ML | 10.7 | - |
| Neck AP | 9.9 | - |
| Midshaft ML | - | 11.8 |
| Midshaft AP | - | 10.7 |
| **Angular Measures (°) <sup>a</sup>** |  |  |
| Angle between subtriangular space and fibulotalar surface | - | 56.0° |
| Angle of fibulotalar articulation (proximal) | [12.5] | [14.0] |
| Angle of fibulotalar articulation (distal) | [67.4] | [64.4] |
| **Indices** |  |  |
| Neck robusticity (ML/AP) x 100 | 1.08 | - |
| Midshaft robusticity (ML/AP) x 100 | - | 1.10 |

<sup>a</sup> Measured as outlined in Marchi (2015).

##### Morphology of StW 573q (left fibula)

StW 573q/1 preserves a small portion of the diaphysis. The head possesses a relatively large, and laterally splayed, styloid process with a slightly concave proximal fibular articular facet. Moving distally, the specimen narrows at the neck, then expands again into the slender diaphysis. At this point, the diaphysis is approximately circular in shape but quickly changes into a more triangular cross-section with the development of its anteriorly directed interosseous crest. This crest is well-developed, especially at the distal 1/3 of StW 573q/1. The division of the crest into medial and lateral lips cannot be seen. On the medial shaft, the area of origin for the extensor digitorum longus muscle is not well delineated. In contrast, laterally, the origin for the fibularis longus better is observed as a hollow adjacent to the interosseous crest. Of its preserved portion, the neck is the narrowest part of the diaphysis. Additionally, the shaft is also comparatively slender (see diaphyseal measures of antimere). Representing roughly 40% of the proximal diaphysis, the dimensions are like those observed mid-diaphysis in the right fibula. On the distal fragment (StW 573q/2), the distal fibulotalar facet angle of StW 573 is more ape-like, while the proximal angle is considerably more like the mean value (17.2; Marchi, 2015) reported for *A. afarensis.* The malleolar fossa is well-developed with slight elevations for the posterior talofibular and posterior tibiofibular ligaments.

##### Preservation of StW 573r (right fibula)

The right fibula is the most complete of the preserved fibular elements. Additionally, its cortical surface is comparatively well-preserved. StW 573r represents approximately 65% of the diaphysis, as well as the right lateral malleolus. Midway (~70mm) down the preserved diaphysis, the specimen had been broken in antiquity but has been subsequently cleaned and rejoined. The largest of the fractures is the one which displaces the lateral malleolus. Breccia holds the two pieces (the diaphysis and lateral malleolus) together. The lateral malleolus is shifted laterally and rotated medially (counterclockwise) around the diaphyseal main axis. The malleolar articular surface, as well as the peroneal groove, are well-preserved and minimally damaged.

##### Morphology of StW 573r (right fibula)

Proximally, the shaft possesses a well-developed interosseous crest which adds to the mediolateral dimensions of the diaphysis. The diaphyseal cross-section is triangular, proximally, like that observed at the distal end of StW 573q/1 and progressively becomes ML flattened (traveling distally). The midshaft robusticity is nearly identical to that of the neck robusticity observed in the left fibula. The shaft exhibits a linear hollow near the position of the extensor hallucis longus muscle but flattens near the origin of the fibularis tertius muscle. Overall, the fibulotalar articulation is inferiorly oriented relative to the long axis of the diaphysis. The distal end of the right fibula is considerably more damaged but appears similar in morphology to the fibulotalar angles reported for its antimere.

#### Relative Lengths of Limb Segments

Tables 8 and 9 report long bone lengths for StW 573 and other hominin fossils, as well as mean values for extant comparative species (also illustrated in Figure 15).

**Figure 15.**
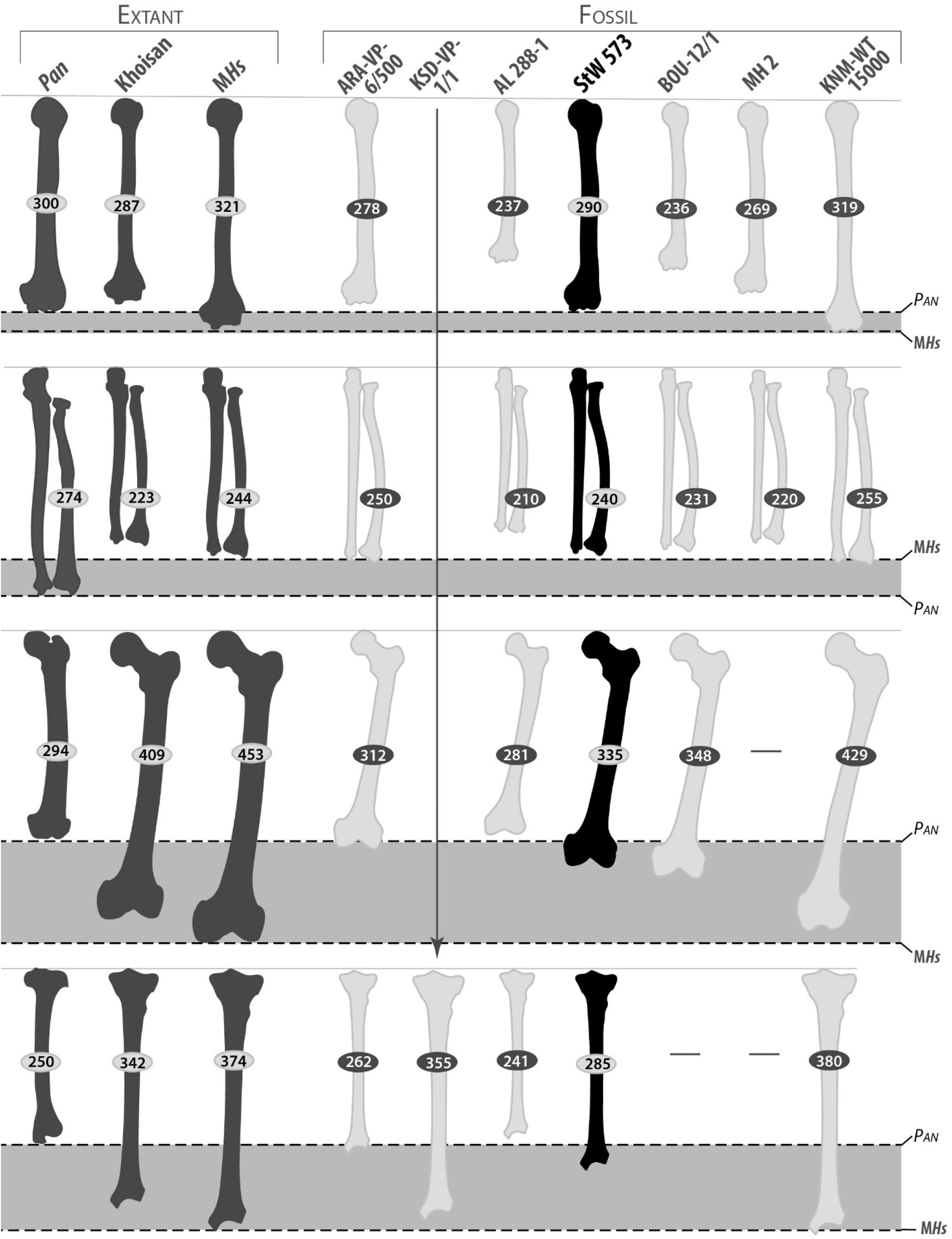
Comparative long bone lengths among extant and extinct species/specimens. Figure modeled after Asfaw et al. (1999). While all limb elements are shown as whole, the fossil bones, in most cases, vary in their completeness. Readers are encouraged to refer to the original publications for an accurate reflection of an element’s form and the degree to which lengths have been estimated. The lengths of modern groups (medium gray) are reported as averages while fossil hominins (black=StW 573, light gray = all remaining fossil hominins) are the estimated or actual values for each element. Modern groups averages were computed here, except for the Khoisan. See Table 9 for fossil data sources. For each element group (e.g., humerus), the horizontal shaded area represents the range between the modern humans (MHs) and chimpanzees (*Pan*) averages.

**Table 8.**
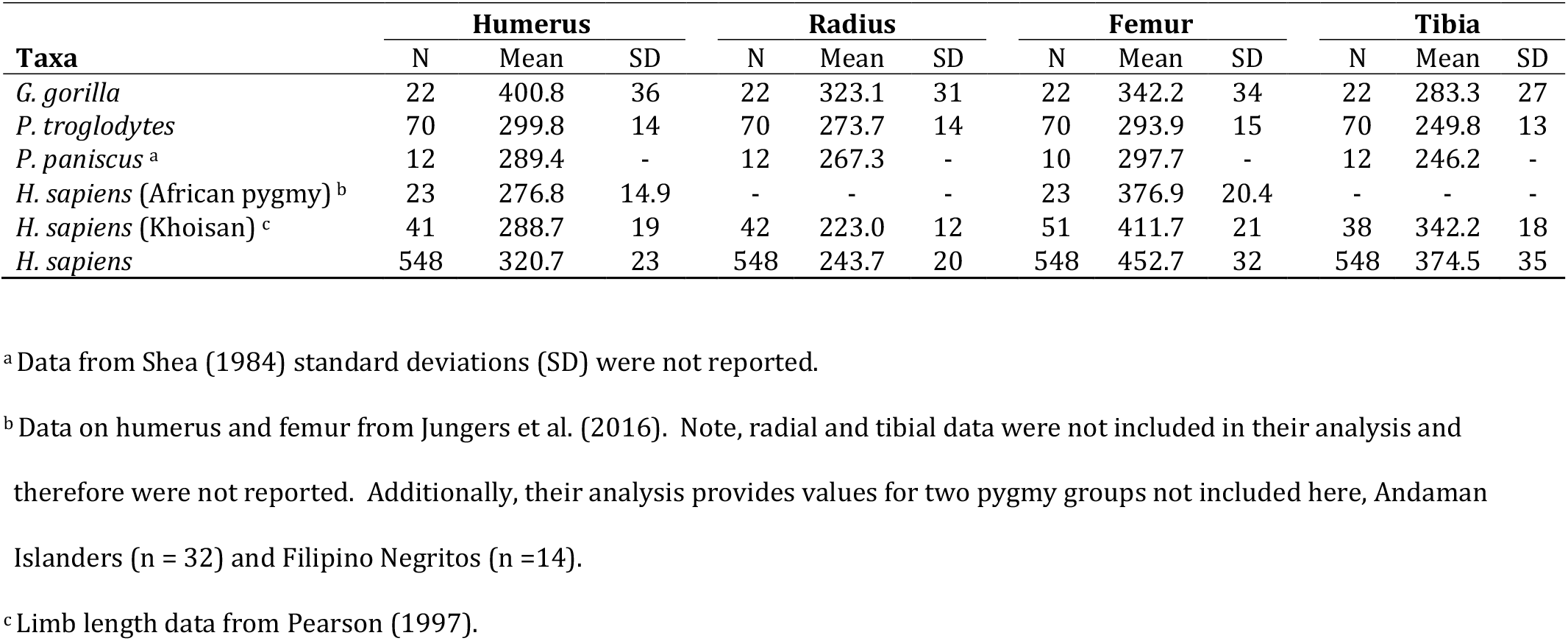
Average Long Limb Lengths Among Extant African Apes and Humans.

Figure 16 illustrates the percentage that each element contributes to the total limb length (humerus + radius and femur + tibia) for extant groups and fossil specimens. As a proportion of the total limb length, the humerus of ARA-VP-6/500 (*Ardipithecus,* 4.4 Ma) and StW 573 (~3.67 Ma) are identical, while the lower limb elements increase in length minimally through this time interval. During this same interval, the most considerable change in limb element contribution to total limb length is a decrease (1.8%) in radius length. Between the time of StW 573 and that of modern humans, the hominin lineage experienced a decrease (3.4%) and increase (3.2%) in radial and femoral lengths, respectively. Moreover, after StW 573, hominins underwent a more moderate change in humeral (2% decrease) and tibial (2.1% increase) lengths. Overall, the changes were greatest in the radius (5.2% decrease from ARA-VP-6/500 to modern humans), followed by the femur (4.2% increase), then, the tibia (3.1% increase) and last, the humerus (2% decrease).

**Figure 16.**
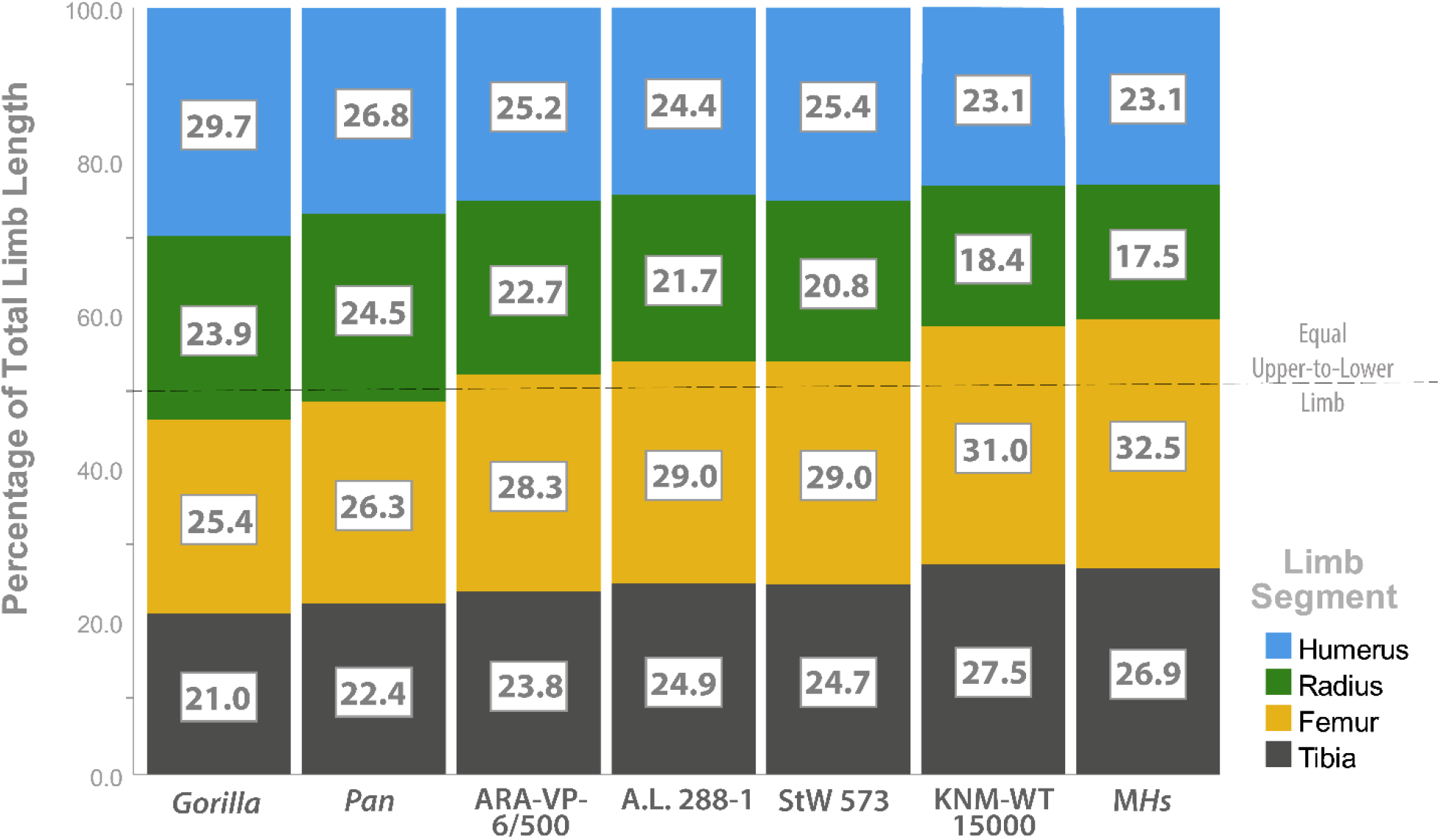
Comparative limb length. Relative limb lengths of extant groups, *Gorilla, Pan,* and modern *H. sapiens* are shown relative to a wide temporal variation of fossil hominins (e.g., ARA-VP-6/500, A.L. 288-1, StW 573 and KNM-WT 15000). For each stacked bar, elements were calculated based upon the percentage (value in white boxes) that each element contributed to the total limb length of that species or specimen. For modern species, percentages represent the average for each element. Species or groups (i.e., ARA-VP-6/500) without complete elements (e.g., humerus and femur) are illustrated based upon limb length estimates reported in the literature and cited within the text.

A regression of the radii and femora suggests possible allometric patterns among the extant groups (Figure 17). Notably, the fossil hominins (A.L. 288-1 and StW 573) are shown to occupy the space between the African apes and modern humans. Similar results hold for the other limb elements, as well. However, with a multivariate approach, differences are more difficult to visualize. Thus, the trends, and the fossil deviations from them are quantified here in the results of our multivariate allometric analysis. For the extant samples, eigenvalues and the percent variance accounted for in each of the principal component analyses are reported in Table 10. Note, the first principal component (PC 1) accounted for between 85.2% – 95.9% of the three samples’ total variances.

**Figure 17.**
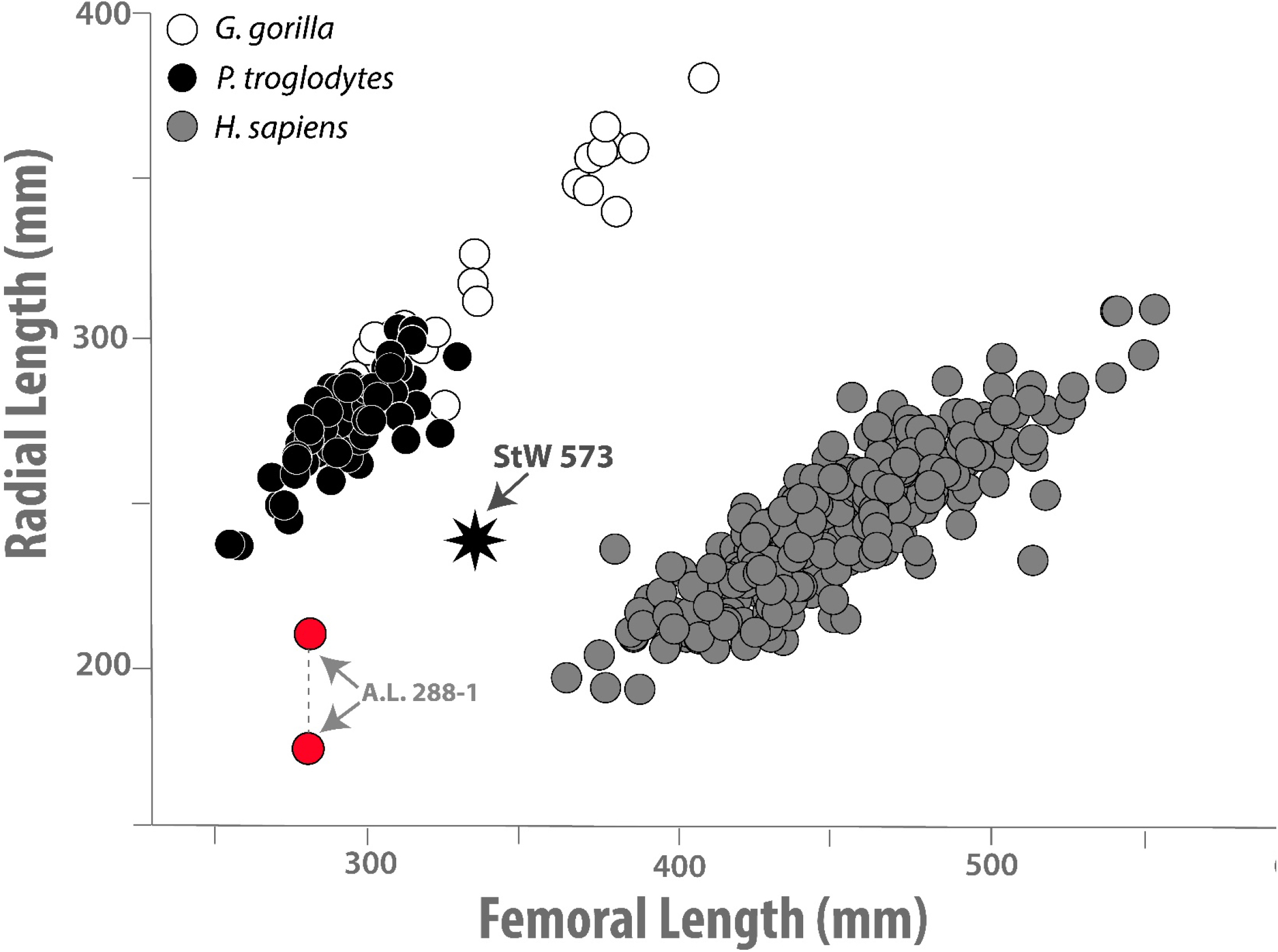
Regression (unlogged) of ape, human and fossil hominin radii and femora. The African apes appear to exhibit a similar allometric relationship when compared to humans. The slopes for each extant group are: (*H. sapiens*) y = 0.51 + 0.54x (R^2^ = 0.71), (*Pan*) y = 54.8 + 0.75x (R2 = 0.65) and (*Gorilla*) y = 31.7 + 0.85x (R^2^ = 0.90). Selected for further allometric analysis, the fossil specimens, StW 573 and A.L. 288-1 are shown to fall between the two modern trends. Because of considerable differences in the estimated radial length of A.L. 288-1 (174 mm – 215 mm), low (Schmid, 1983) and high (Asfaw, et al., 1999) estimates are both shown. There is a small difference (~10mm) in the lengths of the StW 573 radial antimeres, and therefore, the average (235 mm) is plotted. See text for discussion of the bilateral asymmetry.

**Table 9.** Reported Long Limb Lengths of Fossil Specimens Used in this Analysis. Note, square brackets indicate limb lengths which are estimated. For each, see measurement source for more information on element completeness, or estimation methodology.

| **Specimen** | **Humerus** | **Radius** | **Femur** | **Tibia** |
| --- | --- | --- | --- | --- |
| ARA-VP-6/500 <sup>a</sup> | [278] | 250 | [312] | 262 |
| KSD-VP-1/1 <sup>b</sup> | - | - | [418-438] <sup>h</sup> | 355 |
| A.L. 288-1 <sup>c</sup> | [237] | [210] <sup>d</sup> | [281] | [241] |
| **StW 573 <sup>e</sup>** | **290** | **240** | **335** | **285** |
| MH1 <sup>f</sup> | - | - | - | 266 |
| BOU-VP-12/1 <sup>d</sup> | [226-236] | [231] | [335-348] | - |
| KNM-ER 1472 <sup>d</sup> | - | - | 401 | [340] |
| KNM-ER 1481 <sup>d</sup> | - | - | 396 | [333] |
| OH 62 <sup>g</sup> | [264] | [210-246] | [280] | - |
| Dmanisi-1 <sup>c</sup> | [295] | - | 382 | 306 |
| KNM-ER 1808 <sup>g</sup> | - | - | [485] | - |
| KNM-WT 15000 <sup>c</sup> | [319] | 255 | 429 | 380 |

Sources of comparative fossil data:
<sup>a</sup> Lovejoy et al. (2009).
<sup>b</sup> Haile-Selassie et al. (2010).
<sup>c</sup> Pontzer et al. (2010).
<sup>d</sup> Steudel-Numbers & Tilkens (2004).
<sup>e</sup> This study.
<sup>f</sup> Berger et al (2010).
<sup>g</sup> McHenry (1991, 1992).
<sup>h</sup> Ryan and Sukhdeo (2016).

**Table 10.** Eigenvalues and percent variance accounted for in principal component analysis by group.

|  | *H. sapiens* |  | *P. troglodytes* |  | *G. gorilla* |  |
| --- | --- | --- | --- | --- | --- | --- |
|  | Eigenvalue | % variance | Eigenvalue | % variance | Eigenvalue | % variance |
| PC 1 | 0.02026 | 89.76 | 0.03458 | 95.93 | 0.00838 | 85.22 |
| PC 2 | 0.00105 | 4.66 | 0.00071 | 1.96 | 0.00061 | 6.22 |
| PC 3 | 0.00087 | 3.87 | 0.00057 | 1.59 | 0.00047 | 4.77 |
| PC 4 | 0.00038 | 1.71 | 0.00019 | 0.53 | 0.00037 | 3.78 |

The PC 1 loadings of the four elements (i.e., *ln* hum, *ln* rad, *ln* fem and *ln* tib) among the extant samples are reported in Table 11. Measures with the highest PC 1 loadings varied by comparative sample, but all groups possessed at least two elements with high loadings (e.g., > 0.5). Among the three, *Pan* and *Gorilla* were the most similar in their loading pattern. The exception to this, however, is that the tibiae of *Gorilla* increase at a slower rate with increasing size than do those of *Pan.* Humeri consistently possessed the lowest PC 1 loadings across all groups. A unique pattern of higher loading distal (radius and tibia) and lower proximal (humerus and femur) limb elements was observed in *H. sapiens.* These similarities and differences are additionally reflected in the allometric coefficients. Ranging across all groups from 0.9 (negative allometry) to 1.09 (positive allometry), the allometric coefficients are shown in Table 12. All three groups possessed at least one potentially non-isometric limb with computed coefficients differing by at least 5% from isometry (e.g., 1.0). Yet, based upon 95% confidence intervals (Figure 18), humans are the only group for which isometry could be rejected (p < 0.05).

**Figure 18.**
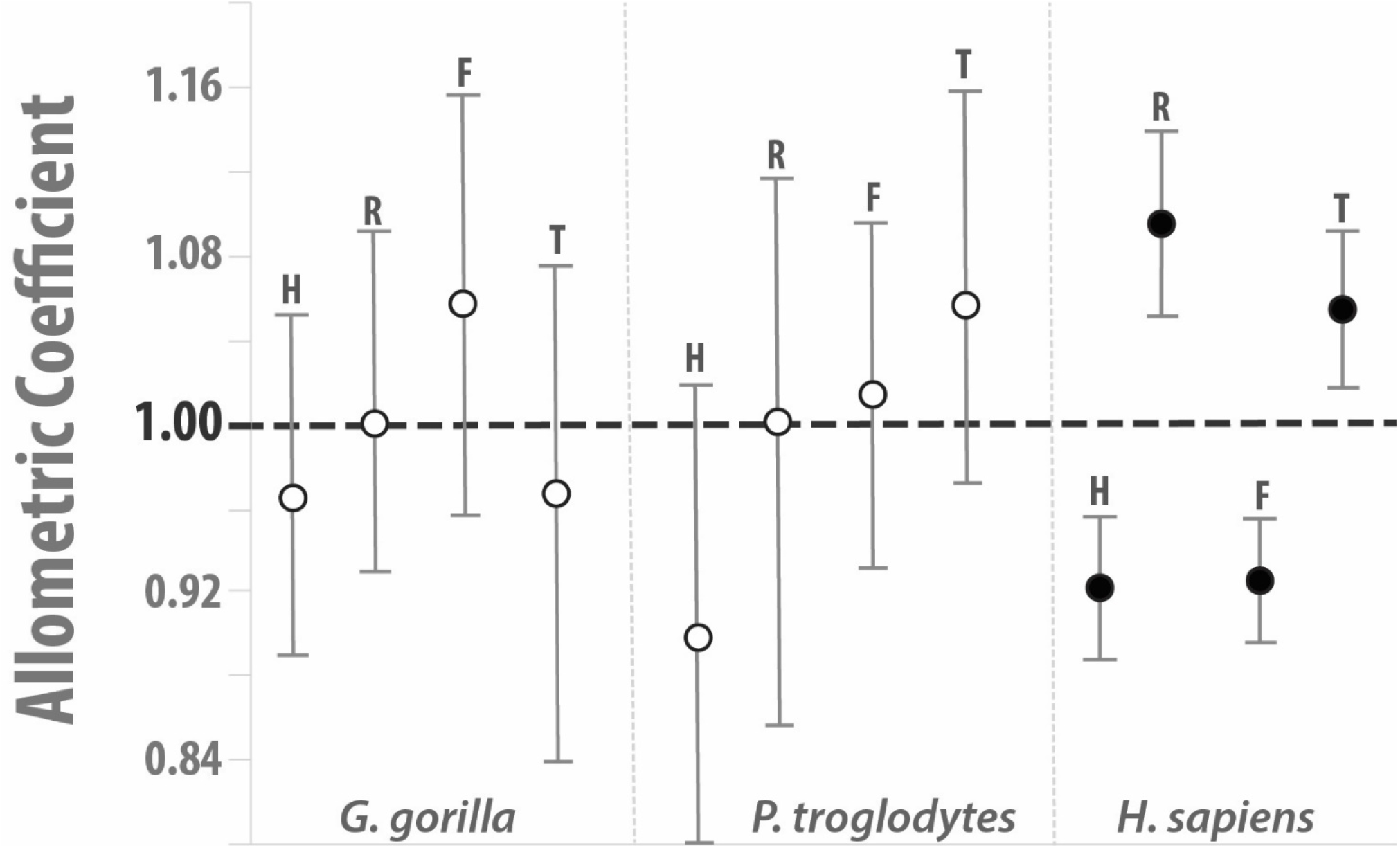
Allometric coefficient confidence intervals. For each species, all four limb elements are shown: humerus (H), radius (R), femur (F) and tibia (T). Circles represent the computed allometric coefficients (given in Table 12) while whiskers represent the 95% confidence interval of each element. Confidence intervals were calculated through bootstrapping (with 2000 replicates) for *Gorilla, Pan* and *H. sapiens.* As outlined in the text, isometry can only be rejected (p < 0.05) if the confidence interval does not include the value one (1.0). Solid circles represent significant deviations from isometry.

**Table 11.** Principal component one (PC 1) loadings by element and species. Variable names indicate the natural log (*ln*) transformed element notation used in the analysis: humerus (*ln* hum), radius (*ln* rad), femur (*ln* fem) and tibia (*ln* tib).

| **Variable** | ***H. sapiens*** | ***P. troglodytes*** | ***G. gorilla*** |
| --- | --- | --- | --- |
| *ln hum* | 0.46083 | 0.44982 | 0.48417 |
| *ln rad* | 0.54637 | 0.50207 | 0.50151 |
| *ln fem* | 0.4613 | 0.50973 | 0.52836 |
| *ln tib* | 0.52565 | 0.53457 | 0.48467 |

**Table 12.** Long limb bone allometric coefficients for African apes and modern humans. Values deviating from isometry (*k* = 1.0) by more than 5% (*k* < 0.95 or > 1.05) are shown in bold. Figure 17 illustrates the 95% confidence intervals of these allometric coefficients and their statistical significance. Variable names indicate the natural log (*ln*) transformed element notation used in the analysis: humerus (*ln* hum), radius (*ln* rad), femur (*ln* fem) and tibia (*ln* tib).

| Variable | Allometric Coefficients (k) |  |  |
| --- | --- | --- | --- |
|  | *G. gorilla* | *P. troglodytes* | *H. sapiens* |
| $\ln$ hum | 0.97 | **0.9** | **0.92** |
| $\ln$ rad | 1.00 | 1.00 | **1.09** |
| $\ln$ fem | **1.06** | 1.02 | **0.92** |
| $\ln$ tib | 0.97 | **1.06** | **1.05** |

In terms of size (e.g., Z-score on PC 1 distribution), StW 573 is not significantly (p = 0.278) different from *Pan.* However, all other comparisons between StW 573 or A.L. 288-1 and the comparative samples were significant for PC 1 (Table 13). Additionally, comparisons of residuals found that in all cases the fossil hominins significantly (p < 0.001) deviated from the allometric axis of all three groups.

**Table 13.** Probabilities of observing fossil hominins (i.e., StW 573 and A.L. 288-1) within each comparative sample’s allometric variation. Significant values (*p* < 0.05) are shown in bold.

|  | *H. sapiens* |  | *P. troglodytes* |  | *G. gorilla* |  |
| --- | --- | --- | --- | --- | --- | --- |
|  | StW 573 | A.L. 288-1 | StW 573 | A.L. 288-1 | StW 573 | A.L. 288-1 |
| PC1 | **<0.010** | **<0.001** | 0.278 | **<0.010** | **<0.050** | **<0.001** |
| Residuals | **<0.001** | **<0.001** | **<0.001** | **<0.001** | **<0.001** | **<0.001** |

### Limb Proportions

Data for limb indices of extant and extinct groups are provided in Tables 14 and 15.

**Table 14.** Limb Indices Among Extant African Apes and Humans.

| Taxa | Brachial |  |  | Crural |  |  | Humerofemoral |  |  | Intermembral |  |  |
| --- | --- | --- | --- | --- | --- | --- | --- | --- | --- | --- | --- | --- |
|  | N | Mean | SD | N | Mean | SD | N | Mean | SD | N | Mean | SD |
| *G. gorilla* | 22 | 80.6 | 1.8 | 22 | 82.9 | 2.6 | 22 | 117.3 | 3.8 | 22 | 115.8 | 2.9 |
| *P. troglodytes* | 70 | 91.3 | 3.1 | 70 | 85.0 | 2.3 | 70 | 102.1 | 2.9 | 70 | 105.6 | 2.4 |
| *P. paniscus*<sup>a</sup> | 12 | 92.3 | - | 10 | 82.7 | - | 10 | 97.4 | - | 12 | 103.4 | - |
| *H. sapiens* (Khoisan)<sup>b</sup> | - | 77.2 | - | - | 83.1 | - | - | 70.1 | - | - | 67.9 | - |
| *H. sapiens* | 548 | 75.7 | 3.3 | 548 | 82.9 | 2.9 | 548 | 71.1 | 2.2 | 548 | 68.4 | 2.0 |

<sup>a</sup> Data from Shea (1984) standard deviations (SD) were not reported.
<sup>b</sup> Indices for Khoisan calculated based upon limb length data reported in Pearson (1997).

**Table 15.** Long limb indices among hominins. Note, square brackets indicate limb lengths which are estimated. For each, see measurement source for more information on element completeness, or estimation methodology.

| **Specimen** | **Brachial** | **Crural** | **Humero-femoral** | **Inter-membral** |
| --- | --- | --- | --- | --- |
| ARA-VP-6/500 | [90.0] | [84.0] | [89.1] | [89-91] |
| A.L. 288-1 | [88.6] <sup>‡</sup> | [85.8] | [84.3] | [85.6] |
| **StW 573** | **82.8** | **85.1** | **86.6** | **85.5** |
| MH 2 | [84.0] | - | - | - |
| BOU-VP-12/1 | [97.9-102.2] | - | [67.5-67.8] | - |
| KNM-ER 1472 | - | [84.8] | - | - |
| KNM-ER 1481 | - | [84.1] | - | - |
| OH 62 | [79.5-93.2] | - | [94.3] | - |
| Dmanisi-1 | - | 80.1 | [77.2] | - |
| KNM-WT 15000 | [79.9] | 88.6 | [74.4] | - |

#### Brachial Index (100 × radius/humerus)

Compared to extant groups, the brachial index of StW 573 (left radius = 82.8, right radius = 86.2) is most like *Gorilla* 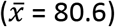 but 2.8 SDs below the mean of chimpanzees 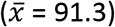 and 2.2 SDs above the mean of modern humans 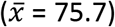. Known specimens of *Ardipithecus* (ARA-VP-6-500, 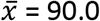) and *Australopithecus* (A.L. 288-1^5^, 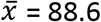 and MH 2, 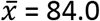) possess shorter forearm bones than the antebrachium mean for *Pan.* The exception is the BOU-VP-12/1 ulna (Bouri, Ethiopia), possibly belonging to *Australopithecus garhi* (Asfaw et al., 1999) exhibiting a chimpanzee-like brachial index, with an estimated range of 97.9-102.2. Relative estimated lengths of the antebrachium of the ~1.8 Ma OH 62 skeleton, from Olduvai (Tanzania), varies widely 79.5 to 93.2 (see Richmond et al., 2002). In contrast, the brachial index of West Turkana (Kenya) *H. ergaster* (KNM-WT 15000 = 71.0) is within the modern human range.

#### Crural Index (tibia length × 100/femur length)

In terms of mean (x = 82.7-85.0) and variation (SD = 2.3-2.9), this index appears relatively stable across all taxa and specimens. Among great apes, *P. troglodytes* possess the relatively longest tibia (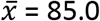; *P. paniscus*, 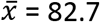; *G. gorilla*, 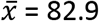). Although they exhibit a greater range of variation, modern humans possess a crural index (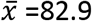; Khoisan, 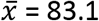) that is practically indistinguishable from those of bonobos and gorillas. StW 573’s crural index (85.1) is similar to those of ARA-VP-6/500 (84.0) and A.L. 288-1 (A. *afarensis)(85.8),* as well as to most individuals of early *Homo,* (estimated values for KNM-ER 1472 = 84.8, KNM-ER 1481 = 84.1, see Table 9). Exceptions include Dmanisi-1^6^ and KNM-WT 15000, which exhibit relatively smaller (80.1) and larger (88.6) indices, respectively, values that are within the range of modern humans.

#### Humerofemoral Index (100 × humerus/femur)

Among apes, *P. troglodytes* (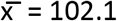) and *G. gorilla* (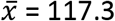) possess humeri that are relatively longer than their femur. At the other extreme, modern humans (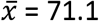; Khoisan, 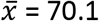) are characterized by relatively longer femora than humeri. Interestingly, StW 573 exhibits a humerofemoral index (86.6) significantly, yet equally, distant from the means for *Pan* and modern humans (5.3 SDs from 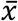 of each). This value is consistent with those derived for *Ar. ramidus* (ARA-VP-6/500 = 89.1) and *A. afarensis* (A.L. 288-1 = 84.3), but quite different from the estimated value for BOU-VP-12/1 (67.5-67.8)^7^. Most fossils of the genus *Homo* show modern human-like ratios (e.g., Dmanisi-1 = 77.2; KNM-WT 15000 = 74.4). We note that, OH 62, sometimes attributed to early *Homo* (e.g., Johanson et al., 1987), but disputed by others (e.g., Clarke, 2017), possesses a humerofemoral index (94.3) approaching the lower bounds of the *Pan* range.

#### Intermembral Index, or IMI, (100 × [humerus + radius] [femur + tibial])

In a pattern like that observed for humerofemoral indices, modern humans possess upper limbs (i.e., humerus + radius) that are significantly shorter when compared to their lower limbs (i.e., femur + tibia). On average, the intermembral indices of modern humans (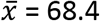; Khoisan, 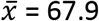) are approximately 2/3 of that of extant African apes (*P. paniscus*, 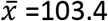; *P. troglodytes*, 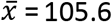; *G. gorilla*, 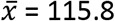). While significantly outside a normal range of variation different, all early hominins considered here have intermembral indices that lie between the ranges of the African apes and modern humans. The reported index of ARA-VP-6/500 (89.0-91.0) is more similar to the chimpanzee mean (6.5 SDs from 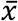), than it is to either modern humans (10.8 SDs from 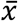) or gorillas (8.6 SDs from 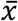). And while somewhat like one another, the *Australopithecus* sample, StW 573 (85.5) and A.L. 288-1 (85.6), exhibits ratios that are considerably different (~8 SDs from 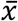) from the chimpanzee and modern human averages.

## Discussion

Previous conclusions that the evolution of hominin limb proportions might have occurred in two stages (Jungers, 1982; Walker and Leakey, 1993; Asfaw et al., 1999; Richmond, et al., 2002; Reno et al., 2005; Young et al., 2010; Holliday, 2012), posit the following: the first stage is characterized by a modest increase in the relative thigh and leg lengths, along with a relative decrease in the length of the antebrachium of *Australopithecus,* relative to a purported basal hominin, *Ardipithecus;* the second stage saw considerable increases in the relative length of the thigh and leg with decreases in the relative brachium and antebrachium lengths, occurring over the span from *Australopithecus* to *H. sapiens.* Unfortunately, sample sizes for these types of comparisons are small, with only a few partial skeletons possessing overlapping skeletal regions of interest. Values used in our analysis for ARA-VP-6/500 (Lovejoy et al., 2009) include estimations for the femur based upon tibial length (and the hypothesized crural index) while the humerus is derived from another individual. Therefore, the slight degree of proportional differences between *Ardipithecus* and *Australopithecus* (Table 15) might simply reflect individual variation within these taxa and, thus, should be subjected to further testing.

Caution is also advised in using extant great apes to model a postulated chimpanzee-human last common ancestor (CHLCA), as research suggests that these groups possess highly derived morphologies (Lovejoy et al., 2009).

Compared to basal hominoids, *Pongo, Pan* and *Gorilla* seem to have independently evolved (i.e., homoplasy) elongated antebrachia (Ward, 2002). Recently, morphological analyses of *A. afarensis* antebrachial fossils (including those of the A.L. 288-1 skeleton) concluded that it did not possess a *Pan*-like antebrachium (Drapeau and Ward, 2007; Haeusler and McHenry, 2004). In agreement, our results show that StW 573 possessed a brachial index of 82.8 – 86.2, midway between the mean chimpanzee and modern human ratios, and also similar to that of *A. sediba* (~84) (Berger et al., 2010; Churchill et al., 2013). Overall, the pattern is one of reduction in relative and absolute radial length (Holliday, 2012) with the purported basal hominin, *Ardipithecus,* possessing ratios within the range of the Miocene ape genus *Proconsul* (Lovejoy et al., 2009). Thus, brachial indices among *Australopithecus* may be reflecting the primitive large-hominoid condition from which extant apes diverged (Ward, 2002).

When regressed against femoral length (see Figure 17), early hominin radial lengths fall directly between the clusters exhibited by modern humans and apes. Unfortunately, most hominins sampled here (e.g., A.L. 288-1, OH 62 and BOU-VP-12/1) have limb length estimates that are widely disparate, which can add uncertainty to their relative placement (Holliday, 2012: Fig. 4). Based on currently available data, it is difficult to determine if the slope of the early hominin radius-to-femur length is more like that of one group (e.g., non-human ape), or the other (e.g., human), but what is clear is that neither StW 573 nor A.L. 288-1 comes close to inclusion within 95% confidence intervals for any extant population (ape or human).

In our sample, crural indices were relatively stable across taxa and time, varying by as little as 3% on average. This suggests that diachronic increases in the hominin lower limb were the result of relatively equal elongation of the thigh and leg segments.

### Interpretation of Allometry

An early analysis of A.L. 288-1’s humerofemoral index suggested that *Australopithecus* possessed ratios that were vastly different from those of chimpanzees and modern humans (Johanson et al., 1982). The primary source of this difference was attributed to relative humeral elongation, femoral shortening, or a combination of both (Jungers, 1982, 1991; Jungers and Stern, 1983; Wolpoff, 1983). Some argue that the proportions of A.L. 288-1 reflect its small body size (Franciscus and Holliday, 1992, Holliday and Franciscus, 2012), as body size and lower limb length have been shown to be positively allometric (Auerbach and Sylvester, 2011). Therefore, small-bodied hominins would be expected to have shorter lower limbs. Yet, compared to A.L. 288-1, StW 573 possesses individual limb lengths that are absolutely longer (by an average of 18%), but nevertheless, exhibits a near equivalent index (Figure 19). Thus, our results suggest that the humerofemoral index of A.L. 288-1 may not be entirely attributable to its small body size. Moreover, the indices of StW 573 and A.L. 288-1 are far above the range for the smallest-bodied modern *H. sapiens* (see Jungers, 2009), and therefore, allometry alone most probably does not explain the humerofemoral proportions observed in *Australopithecus* (contra Franciscus and Holliday, 1992; Holliday and Franciscus, 2012).

**Figure 19.**
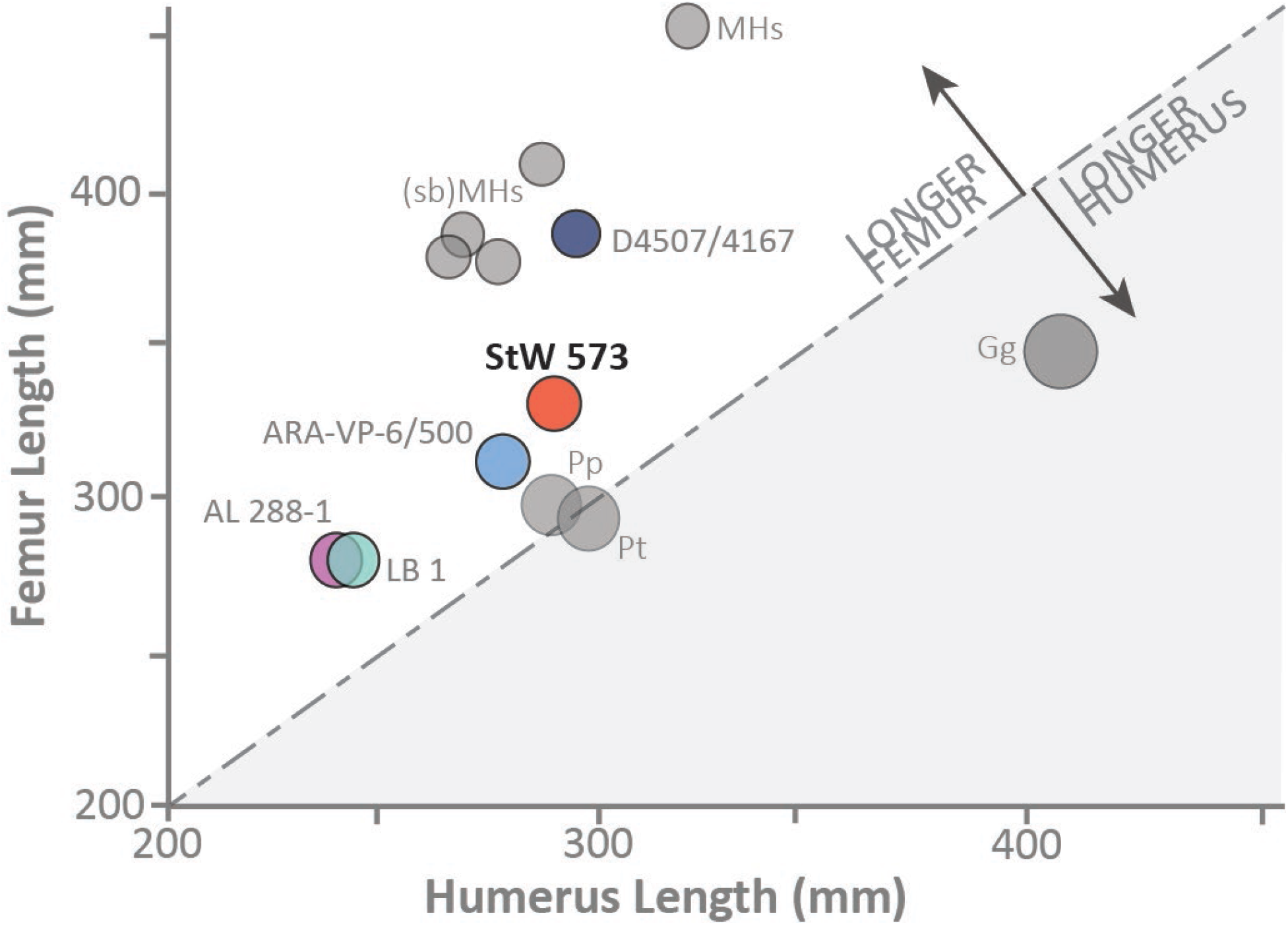
Humerofemoral indices for extant and extinct groups or specimens. Populations, such as modern *Homo sapiens* (MHs), small-bodied modern humans, more commonly referred to as pygmies (sbMHs), and the African apes (*Gorilla gorilla, Gg; Pan troglodytes, Pt; P. paniscus, Pp*) are represented by their group’s mean value. Fossil hominins preserving both long limb bones were also included. Note, circles representing each specimen (or species) are scaled to their humerofemoral index. Therefore, larger circles represent higher humerofemoral indices (e.g., gorillas).

The presence of allometry in human limb lengths and their evolutionary changes has been well-studied (Trinkaus, 1981, Holliday, 1999, Jantz and Jantz, 1999; Haeusler and McHenry, 2007). Limb longitudinal growth among humans is non-isometric and complex, as an individual’s sex may even play a role in limb length variation (Holliday and Ruff, 2001; Sylvester et al., 2008). However, a near-universal trend towards positively allometric distal (e.g., radius and tibia) and negatively allometric proximal (e.g., humerus and femur) limb elements – has been observed in humans (Sylvester et al., 2008). Because of this unique pattern, human distal elements should compose a higher proportion of the total limb length in larger individuals. In fact, males have been shown to possess higher brachial indices than females (Trinkaus, 1981, Aiello and Dean, 1990, Ruff, 1994) possibly highlighting this disjunction between proximal limb (e.g., humerus and femur) and distal limb (e.g., radius and tibia) element growth. As limb proportions are often used as a salient gauge of locomotor behavior (Jungers, 1985), it is essential to assess the allometric patterns observed among fossil hominins. If upheld by subsequent studies, our analysis suggests that *Australopithecus* (e.g., StW 573) possessed an allometric scaling pattern (in limb lengths) that differs from that of any living African ape or modern human.

As observed here, the humerus was the only element that deviates consistently from isometry in the same (negative allometry) direction across all species. Yet, African apes exhibit patterns more like one another than either does to modern humans. And while all species exhibited some isometric deviations, as evidenced by the allometric coefficients, humans were the only group for which isometry could be rejected. However, we must note that sample size may influence the observation (or not) of whether isometry is observed. As Sylvester et al. (2008) have remarked, small sample sizes (<50-60), at least among humans, typically failed to reject isometry. In our analysis, *Gorilla* was the only taxon that did not exceed this sample size threshold.

By conducting a multivariate allometric analysis, we have rejected the hypothesis that *Australopithecus* (represented here by StW 573 and A.L. 288-1) possessed a human-like pattern of allometry. Our data, therefore, suggest that their limb lengths (and proportions) do not result from their shared status as small-bodied individuals using a modern human allometric scale. And while not considerably different in ‘size’ from our *Pan* sample (see PC 1), StW 573 exhibited significant deviations from the allometric pattern of both African apes. The case for A.L. 288-1 was significantly different across all comparisons with apes and humans. However, this may also reflect its considerably smaller body size (i.e., outside of the ‘size’ range exhibited by all extant groups).

Currently, there are too few data to determine if a single allometric pattern typified all of *Australopithecus,* but we can conclude that the patterns exhibited by the two representatives in this study are different from those observed in our extant samples. It should be noted that the relative completeness of each skeleton, A.L. 288-1 and StW 573, that we used to represent *Australopithecus* means that these are robust results. However, we do not assert any definitive ancestral-descendant relationships between these species and later *Homo.* We understand that each may have its own evolutionary trajectory, as cladistic analyses have shown South African *Australopithecus* to be one of the most phylogenetically unstable hominin groups (Skelton et al. 1986; Chamberlain and Wood, 1987; Skelton and McHenry, 1992; Strait et al., 1997; Strait and Grine, 2004; Kimbel et al., 2004). Among other factors, this instability, accentuated by the high degree of morphological variation at Makapansgat and Sterkfontein (which has a much larger sample), has led some authors to argue that two *Australopithecus* species are represented within these deposits (Clarke 1989, 1994, 2013; Clarke and Kuman, 2018; but see also Crompton et al., 2018).

### Functional Anatomy

Regardless of taxonomy, it is often implied that South African *Australopithecus* is more arboreal than its eastern African congeners (Johanson et al., 1987; McHenry and Berger, 1998a, b; Gordon et al., 2007). Much of this interpretation is based on the analyses of lower limb fossils. Combined with a relatively large head, the femoral neck of StW 573 is small relative to some Sterkfontein specimens, StW 598 and StW 99 (Partridge, et al., 2003). In many of its features, the thigh and leg of StW 573 exhibit clear evidence of habitual terrestrial bipedality. Some lower limb traits, such as the bicondylar angle and flattened femoral condyles, are developmentally plastic and only develop during repeated use of bipedal gaits (Frost, 1990, Duren, 1999, Hamrick, 1999, Tardieu, 2010). Therefore, the presence of these features in StW 573 adds considerable weight that it was an obligate terrestrial biped. In addition, the ridge for the anterior tibiofibular ligament of StW 573 is large and extends distally to lie anterior to the distal articular fibular surface – a feature which Carlson et al. (in press) have suggested creates stability within the joint, especially during bipedal walking. Furthermore, substantial contractile forces generated by the quadriceps femoris during pre-swing phase of bipedal gait is indicated on the StW 573 femora by the elevated lateral lip of the patellar surface, and its association with the presence of a high bicondylar angle (Ward, 2002). Contrasted with values for *A. sediba,* the medial malleolus of StW 573 is relatively thin, but comparable to values observed for *A. anamensis* (KP 29285) and *A. afarensis* (A.L. 288-1) (see Zipfel et al., 2001: Fig. S6).

These features, however, also appear alongside primitive morphological retentions on the tibiae of StW 573, such as the absence of a clear soleal line, an ape-like interosseous border, and a trapezoidal tibiotalar surface (see Aiello and Dean, 1990; Zipfel et al., 2011). Taken together, these features may indicate that while clearly bipedal, the leg musculature of StW 573 was yet to be reorganized, as it is I modern humans. Presently, it is not clear if these retentions would have significantly impacted the range of motion or energetic efficiency during climbing or walking movements, or whether they merely represent normal variation. These features are noteworthy, however, as they reflect the position of some of the major plantar-flexors (i.e., soleus and tibialis posterior) and dorsi-flexors (i.e., tibialis anterior) of the foot. Yet, within our observations of StW 573’s anatomy, mainly the upper limb, climbing and/or suspension appear to have continued to play a role in its adaptation (Crompton et al., 2018).

In fact, the humerus of StW 573 possessed a widely flaring and high (i.e., proximal) lateral supracondylar ridge, as well as a large epicondyle, which in apes reflects a well-developed brachioradialis and the long (e.g. longus vs. brevis) wrist and digital extensor muscles, respectively (Miller, 1932; Knussman, 1967; Senut and Tardieu, 1985). Crompton et al. (2018) have suggested that these muscles, especially the brachioradialis, would have increased power in elbow postures (pronation and flexion) that were most likely employed during climbing. During contraction, these muscles generate forces across the elbow that are mainly resisted by ulnar keeling (Drapeau, 2008). In StW 573, the proximal ulnar keel is intermediate in its development between the more arboreal (*Pongo* and *Pan*) and the more terrestrial groups (*Gorilla* and *H. sapiens*) and is similar to other fossil hominins of *A. afarensis* and *A. africanus.*

Among fossil hominins, flatter keeling is a possible indicator of a relative reduction in the muscule mass (or power generation) of the wrist and superficial digital flexors (Drapeau, 2012). Based upon the work of Drapeau (2008, 2012; Drapeau et al., 2005), this suggests less upper limb use by StW 573, relative to apes, with extended elbow postures, such as suspensory behaviors. While they did not report the proximal angle, Churchill, et al. (2013) argued that the keeled ulna of MH 2 indicated a significant adaptation to forelimb climbing and suspension. And recently, Rein et al. (2017) have noted that the ulna of *A. sediba* is more strongly indicative of suspensory behavior than *A. afarensis* (e.g., A.L. 438-1), but they add that forelimb suspension was performed at low frequencies, more similar to patterns observed in extant *P. troglodytes* (and *Presbytis melalophos*) compared to hylobatids.

Limb torsion, another possible epigenetic trait (Martin and Saller, 1959; King et al., 1969; Sarmiento, 1985; Pieper, 1998), may provide further clues to limb function and trait polarity (i.e., primitive/derived). Low humeral torsional values among early hominins possibly represent primitive retentions since basal hominoids, such as *Proconsul* and *Sivapithecus,* as well as extant *Hylobates* (Begun and Kordos, 1997; Begun et al., 1997; Ward, 1997; Larson, 2009), all exhibit this feature. Humeral torsion may also reflect shoulder orientation and torso organization (Ward, 2002: 198 and references, therein). Larson (2009) has indicated that low humeral torsion values typically occur in connection with high-positioned scapulae (with a cranially directed glenoid fossa) and a funnel-shaped thorax. Therefore, the low humeral torsional values observed in StW 573 appear to be a reflection of its apically narrow thorax (Heaton et al., in prep.) and relatively (to humans) cranially oriented glenoid fossa (Carlson et al., in prep.).

Therefore, we conclude that bipedalism was actively selected for in StW 573, but that many features of the upper limb may represent primitive retentions of arboreal adaptations.

### Potential Issues

Although critically important to understanding the nature of early bipedalism, there are relatively few fossil hominin skeletons complete enough to provide complete data. Indeed, as most of the hominin fossil record consists of fragmentary remains, bone lengths are, out of necessity, often estimated from partial elements. Here, we have relied upon published values of limb lengths and/or estimates and acknowledge that in certain cases estimates may have considerable error ranges. And as a result, conclusions about the long limb proportions of some specimens (e.g., OH 62) may be problematic (Korey, 1990; Hartwig-Scherer and Martin, 1001; Richmond et al., 2002; Haeusler and McHenry, 2004; Reno et al., 2005). Estimates can vary dramatically by the method chosen. The estimates for radial length of A.L. 288-1, for example, vary by nearly 20-25%, i.e., from 174-215 mm (Schmid, 1983; Asfaw et al., 1999). Furthermore, Holliday (2012) and Churchill et al. (2013) have expressed caution regarding the low humerofemoral index of the taxonomically unassigned BOU-VP-12/1 (see Reno et al., 2005).

Beyond methodological differences, Ohman et al. (2002) have also suggested that potential pathologies and/or ontogeny may be problematic for determining limb length estimates. Relevant to our analysis of StW 573, Heile et al. (in prep.) have concluded that antemortem trauma may be responsible for differences in bilateral asymmetry observed in the antebrachium. As noted in our descriptions, there are significant differences in the curvature of the left and right antebrachial elements of StW 573, as well as potential differences in length. In this context, it is unfortunate that until virtual reconstructions are completed, many standard measurements of the right radius and ulna could not be confidently be measured, at the time of study. However, the difference between the left and right radius (in maximum length) are approximately 4% (10 mm). Fortunately, these differences in radial length appear to only minimally impact the overall pattern of proportions observed in StW 573. If using the straighter (right) radius, to compute total limb length, the relative proportion (to total limb length) of the remaining elements (i.e., humerus, femur, and tibia) would each be 0.2% smaller than what is reported, above. Because it is ~10mm longer than the left, the brachial index (if using the right radius) would increase to 86.2 (from 82.8). Regardless of which radius is used, these values are still consistent with those of other members of *Australopithecus.* This finding illustrates the difficulty of drawing conclusions from an incomplete fossil record, even in a remarkably complete skeleton, such as StW 573. Had both sides not been preserved, we would not have been aware of this bilateral asymmetry.

As it is often difficult to agree on the most appropriate reference sample to use (e.g., ape or human), the prediction of indices, or limb lengths for that matter, from other complete dimensions (e.g., articular dimensions) may serve as another potential source of error in quantifying features of early hominins. The completeness of StW 573 offers a rare opportunity to test the power of these predictive models. Using data presented here for StW 573, and formulae from the noted sources, we were able to test the effectiveness of two such models: (1) predicting the intermembral index from humerofemoral dimensions (Jungers, 2009), and (2) predicting femoral and humeral lengths from femoral head diameter (Jungers et al., 2016). Similar to previous findings (Jungers, 2009), we observed that regardless of the reference population, the modern human model was the poorest predictor of the StW 573 intermembral index. All formulae tended to underpredict (−7.1%) the intermembral index of StW 573 (combined sample predicted StW 573 value = 79.4, actual StW 573 value = 85.5). Even while using only small-bodied reference samples, predictions were even more unsatisfactory (Andaman Islanders predicted StW 573 value = 71.8 (−16.0%); Khoisan predicted StW 573 = 69.8 (−18.4%)). This further supports the notion that StW 573 does not scale allometrically in the same fashion as do African apes or small-bodied modern humans.

Yet, we did find some combinations (e.g., femoral head diameter and femoral length) that were reasonably accurate in their predicted values for individual limb elements. While using a modern human reference sample (which exhibits relatively longer lower limbs and shorter upper limbs), the observed results were somewhat be expected. Using formulae from Jungers et al. (2016) the femoral model overpredicted (+11.7%) femoral length (374.3 mm vs. 335 mm); while it underpredicted (−9.1%) humeral length (263.6 mm vs. 290 mm). Plotting its measured values, StW 573’s femoral length falls within the 95% confidence interval of the predicted values for small-bodied modern *H. sapiens* samples while its humerus falls above it (Figure 20). The disparity in predicted versus actual values in StW 573 derives from its unique combination (among extant taxa) of limb lengths.

**Figure 20.**
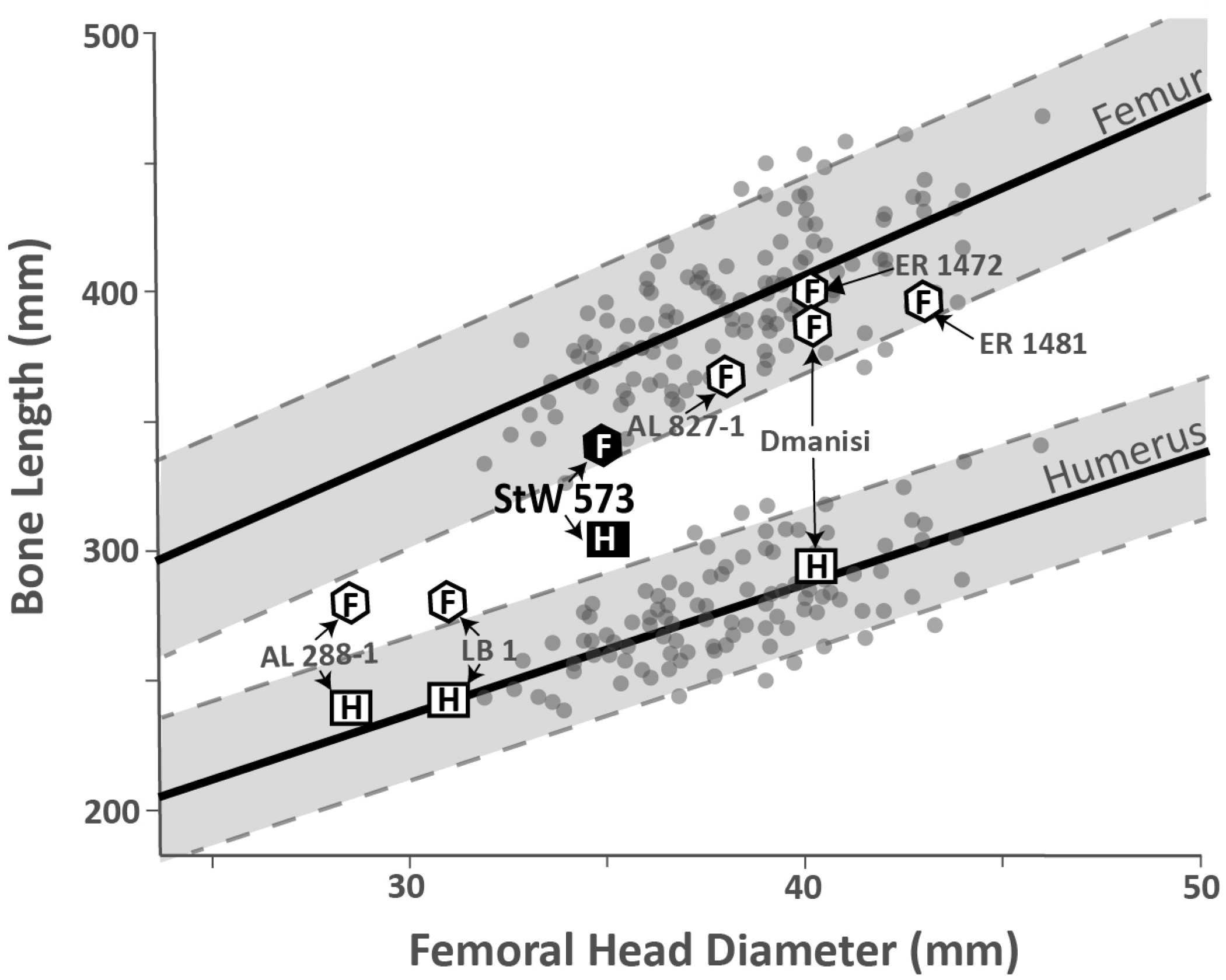
Regression of femoral head diameter to humerus and femoral length. Shown here, the femoral head diameter regressed on a modern small-bodied sample of modern human humeral and femoral lengths (figure adapted, and data obtained, from Jungers et al., 2016). Solid dots represent individual measurements for modern humans, while the dashed lines represent the 95% confidence intervals for each element. Fossil specimens are shown with their humeral or femoral lengths designated as boxes, or hexagons, respectively. Data for StW 573 are based upon those values reported in this study.

As discussed here, limb allometry in StW 573 (and A.L. 288-1) is significantly different from that observed in apes and humans. Therefore, estimations based upon models from these extant groups will undoubtedly result in some level of error.

## Conclusions

Our analyses indicate that limb lengths, and therefore proportions of StW 573 do not follow a modern human allometric pattern. The functional significance of isometry, while important, is mainly unresolved (Sylvester et al., 2008). Current computer modeling, however, has suggested that the lack of isometry in modern humans may account for at least 1/3 of the variation in energy expenditure.

Excluding the crural index, the pattern of relative limb proportions of StW 573 was not observed in any living population of modern humans, even among the smallest-bodied (i.e., pygmy) groups, and the pattern was often unlike those of suspensory-adapted apes. Typically, with increases in hominoid body size, the relative length of the upper limb increases at a faster rate than the lower limb (Jungers, 1985). Therefore, among primates, large-bodied taxa would be expected to possess higher intermembral indices (or IMI’s) than smaller groups. Yet in the similarly-sized *Pan* and StW 573, the IMI’s were markedly different (respectively, >103 and 86.1) suggesting considerably less below-branch suspension characterized the behavioral repertoire of the latter. The IMI of StW 573 appears to be a primitive retention, similar in value to Old World monkeys, *Proconsul* and *Ar.* ramidus, while its brachial index indicates shortening of the antebrachium relative, to those same groups (Lovejoy et al., 2009: Table S3 and Fig. S3).

Due to the discordant combinations of traits commonly found among primates, early limb comparative studies indicated complications in associating locomotor types with taxa (Erickson, 1963). Current evidence suggests that StW 573 possessed arboreal traits, in the upper limb, while exhibiting clear evidence for habitual bipedality in the lower limb. As an adaptive complex, Crompton et al. (2018) suggested that this post-cranial suite of features would have reduced StW 573’s energetic efficiency – relative to apes – in arboreal climbing while also enhancing its ability to walk medium-to-long distances. However, there is no discipline-wide consensus on the selective nature of arborealism among early hominins. As Ward (2002) notes, the resolution of the fossil record may prevent precise locomotive interpretations, as retained arboreal traits may have been adaptive, neutral or even actively selected against. Current differences among eastern and southern African *Australopithecus* may reflect differences in selective pressures related to arborealism (Green et al., 2007), and as the most complete *Australopithecus* skeleton, StW 573 may help understand the polarity of these traits. While the limb proportions of A.L. 288-1 and StW 573 are similar, in many ways, and perhaps even on the same allometric scale, this may not equate to functional equivalence. An expansion of post-cranial studies is necessary to understand these similarities and differences fully.

To that end, an ecomorphological reconstruction of StW 573’s post-cranial anatomy has been qualitatively considered in Crompton et al. (2018). In StW 573’s bony labyrinth, Beaudet et al. (In Press) found a mixture of ape-like and human-like features within the semicircular canals and cochlea, respectively, and suggested that its obligatory bipedalism was paired with some degree of arborealism. The first four footbones that led eventually to the discovery of this complete skeleton were published by Clarke and Tobias (1995), who observed that there was some degree of mobility at the joint of the medial cuneiform with the first metatarsal. They concluded that this mobility of the hallux (i.e. big toe) would have assisted in arboreality. Beyond that, further post-cranial studies are nearing completion on the pectoral girdle (Carlson et al., in prep.) and hand (Jashashvili et al., in prep.) providing further clarification on StW 573’s locomotive habitus and manual abilities. As South African hominins are often the most diverse (i.e. greatest degree of variation) and the least stable in phylogenetic studies, the extraordinary completeness of StW 573 will allow us to test hypotheses in previously impossible ways and to understand further the evolutionary forces that shaped the *bauplan* of *Australopithecus.*

## Footnotes

1 Data were accessed via the file “Humans, Apes and Fossil Hominins” on Pontzer’s Australopithecus.org site (Data Mine section).

2 Some extant human sample comparisons include small-bodied, or pygmy, samples defined as a population with male mean statures of <150cm (Migliano et al., 2007). Individual group names (e.g., Khoisan and African Pygmy) follow the nomenclature of Jungers et al. (2016).

3 The most recent version of PAST (ver. 3.21) does not contain multivariate allometry. PAST is preprogrammed to produce bootstrapping with 2000 replicates in its computation of multivariate allometry (ver. 2.71).

4 The medial malleolar thickness was calculated following Zipfel et al., (2011: Fig. S6).

5 We note that Lovejoy et al. (2009, SOM p. 4) report a computed brachial index for A.L. 288-1 of 92, without an actual radial length reported. Thus, we relied upon published limb length values for our analyses.

6 Dmanisi-1 is used by Pontzer et al (2010: online database) to denote bones belonging to a single large adult individual (Lordkipanidze et al., 2007) which include elements D4507 (left humerus), D4167 (right femur) and D3901 (right tibia).

7 See Churchill et al (2013) for a critique of, and explanation for, the humerofemoral value observed in BOU-VP-12/1.

## References

Aiello, L., Dean, C., 1990. An Introduction to Human Evolutionary Anatomy. Academic Press.

Alemseged, Z., Spoor, F., Kimbel, W.H., Bobe, R., Geraads, D., Reed, D., Wynn, J.G., 2006. A juvenile early hominin skeleton from Dikika, Ethiopia. Nature. 443, 296–301.

Almécija, S., Tallman, M., Alba, D.M., Pina, M., Moyà-Solà, S., Jungers, W.L., 2013. The femur of *Orrorin tugenensis* exhibits morphometric affinities with both Miocene apes and later hominins. Nature Communications. 4, 2888.

Asfaw, B., White, T., Lovejoy, O., Latimer, B., Simpson, S., Suwa, G., 1999. *Australopithecus garhi*: A new species of early hominid from Ethiopia. Science. 284, 629–635.

Auerbach, B.M., Sylvester, A.D., 2011. Allometry and apparent paradoxes in human limb proportions: Implications for scaling factors. American Journal of Physical Anthropology. 144, 382–391.

Bass, W.M., 2004. Human Osteology. A laboratory and field manual.

Beaudet, A., Clarke, R., Bruxelles, L., Carlson, K.J., Crompton, R., de Beer, F., Dhaene, J., Heaton, J.L., Jakata, K., Jashashvilli, T., Kuman, K., McCloymont, J., Pickering, T.R., Stratford, D. (In Press). The bony labyrinth of StW 573 (“Little Foot”): Implications for early hominin evolution and paleobiology. Journal of Human Evolution.

Begun D.R., Kordos L., 1997. Phyletic affinities and functional convergence in Dryopithecus and other Miocene and living hominids. In: Begun D.R., Ward C.V., Rose M.D., (Eds.), Function, phylogeny, and fossils: Miocene Hominoid evolution and adaptations. New York: Plenum Press. 291–316.

Begun D.R., Ward C.V., Rose M.D., 1997. Events in hominoid evolution. In: Begun DR, Ward CV, Rose MD, editors. Function, phylogeny, and fossils: Miocene Hominoid evolution and adaptations. New York: Plenum Press. 389–415.

Behrensmeyer, A.K., 1978. Taphonomic and Ecologic Information from Bone Weathering Taphonomic and ecologic information from bone weathering. Paleobiology. 4, 150–162.

Berger, L.R., De Ruiter, D.J., Churchill, S.E., Schmid, P., Carlson, K.J., Dirks, P.H.G.M., Kibii, J.M., 2010. *Australopithecus sediba:* A New Species of Homo-Like Australopith from South Africa. Science. 328, 195–204.

Blumenschine, R.J., Marean, C.W., Capaldo, S.D., 1996. Blind tests of inter-analyst correspondence and accuracy in the identification of cut marks, percussion marks, and carnivore tooth marks on bone surfaces. Journal of Archaeological Science. 23, 493–507.

Buikstra, J.E., Ubelaker, D.H., 1994. Standards for data collection from human skeletal remains. Arkansas Archaeological Survey Research Series. 44.

Carlson, K., et al., (In Preparation) The pectoral girdle of StW 573

Chamberlain, A.T. and Wood, B.A., 1987. Early hominid phylogeny. Journal of Human Evolution. 16, 119–133.

Churchill, S.E., Holliday, T.W., Carlson, K.J., Jashashvili, T., Macias, M.E., Mathews, S., Sparling, T.L., Schmid, P., De Ruiter, D.J., Berger, L.R., 2013. The upper limb of *Australopithecus sediba*. Science. 340, 1233477.

Clarke, R.J., 1989. A new *Australopithecus* cranium from Sterkfontein and its bearing on the ancestry of *Paranthropus*. In: Grine, F. (Ed.), Evolutionary History of the “Robust” Australopithecines. New York: Aldine de Gruyter. 285–292.

Clarke, R.J., 1994. Advances in understanding the craniofacial anatomy of South African early hominids. In: Corruccini, R.S., Ciochon, R.L., (Eds.), Integrative Paths to the Past: Essays in Honor of F. Clark Howell. New Jersey: Prentice-Hall, pp. 205–222.

Clarke, R.J., 1998. First ever discovery of a well-preserved skull and associated skeleton of *Australopithecus*. South African Journal of Science. 94, 460–463.

Clarke, R.J., 2017. *Homo habilis:* the inside story. In: Sahnouni, M., Semaw, S., Garaizar, J.R. (Eds.), Proceedings of the II Meeting of African Prehistory. Centro Nacional de Investigación sobre la Evolución Humana, Burgos, Spain, 25–51.

Clarke, R.J., Tobias, P. V, 1995. Sterkfontein member 2 foot bones of the oldest South African hominid. Science. 269, 521–524.

Clarke, R.J., Kuman, K. 2018. The skull of StW 573, a 3.67 Ma *Australopithecus* skeleton from Sterkfontein Caves, South Africa bioRxiv 483495 [Preprint]. December 4, 2018; doi: https://doi.org/10.1101/483495

Conroy, G.C., Pontzer, H., 2012. Reconstructing Human Origins: A Modern Synthesis, 3rd ed. W.W. Norton, New York.

Corruccini, R.S., 1978. Comparative osteometrics of the hominoid wrist joint, with special reference to knuckle-walking. Journal of Human Evolution. 7, 307–321.

Corruccini, R.S., McHenry, H.M., 2001. Knuckle-walking hominid ancestors. Journal of Human Evolution. 40, 507–511; discussion 513–520.

Creighton, G.K. and Strauss, R.E., 1986. Comparative patterns of growth and development in cricetine rodents and the evolution of ontogeny. Evolution. 40, 94–106.

Crompton, R.H., Pataky, T.C., Savage, R., D’Août, K., Bennett, M.R., Day, M.H., Bates, K., Morse, S., Sellers, W.I., 2012. Human-like external function of the foot, and fully upright gait, confirmed in the 3.66 million year old Laetoli hominin footprints by topographic statistics, experimental footprint-formation and computer simulation. Journal of the Royal Society, Interface. 9, 707–19.

Crompton, R.H., Sellers, W.I., Thorpe, S.K.S., 2010. Arboreality, terrestriality and bipedalism. Philosophical transactions of the Royal Society of London. Series B, Biological sciences. 365, 3301–14.

Crompton, R.H., Thorpe, S., Weijie, W., Yu, L., Payne, R., Savage, R., Carey, T., Aerts, P., Van Elsacker, L., Hofstetter, A., Günther, M., Richardson, J., 2003. The biomechanical evolution of erect bipedality. CFS Courier Forschungsinstitut Senckenberg. 135–146.

Crompton, R.H., Vereecke, E.E., Thorpe, S.K.S., 2008. Locomotion and posture from the common hominoid ancestor to fully modern hominins, with special reference to the last common panin/hominin ancestor. J. Anat. 212, 501–543.

Crompton, R.H., Yu, L., Wang, W., Gunther, M., Savage, R., 1998. The mechanical effectiveness of erect and “bent-hip, bent-knee” bipedal walking in *Australopithecus afarensis*. Journal of Human Evolution. 35, 55–74.

Crompton, R.H., McClymont, J., Thorpe, S., Sellers, W., Heaton, J.L., Pickering, T.R., Pataky, T. Straftord, D., Carlson, K.J., Jashashvili, T., Beaudet, A., Elton, S., Bruxelles, L., Goh, C., Kuman, K., Clarke, R.J., 2018. Functional Anatomy, Biomechanical Performance Capabilities and Potential Niche of StW 573: an Australopithecus Skeleton (circa 3.67 Ma) From Sterkfontein Member 2, and its significance for The Last Common Ancestor of the African Apes and for Hominin Origins. bioRxiv 481556 [Preprint]. November 29, 2018; doi: https://doi.org/10.1101/481556

Dainton, M., 2001. reply: Did our ancestors knuckle-walk? Nature. 410, 324–326.

Deloison, Y., 2003. Anatomie des os fossiles de pieds des hominidés d’Afrique du Sud datés entre 2, 4 et 3, 5 millions d’années. Interprétation quant à leur mode de locomotion. Biométrie humaine et anthropologie. 21, 189–230.

DeSilva, J.M., Throckmorton, Z.J., 2010. Lucy’s flat feet: The relationship between the ankle and rearfoot arching in early hominins. PLoS ONE. 5, e14432.

DeSilva, J.M., 2009. Functional morphology of the ankle and the likelihood of climbing in early hominins. Proceedings of the National Academy of Sciences. 106, 6567–6572.

Drapeau, M.S., 2008. Articular morphology of the proximal ulna in extant and fossil hominoids and hominins. Journal of Human Evolution. 55, 86–102.

Drapeau, M.S., 2012. Forelimb adaptations in Australopithecus afarensis. In: Reynolds, S.C., Gallagher, A. (Eds.), African Genesis: Perspectives on Hominin Evolution. 223–247.

Drapeau, M.S., Ward, C.V., Kimbel, W.H., Johanson, D.C. and Rak, Y., 2005. Associated cranial and forelimb remains attributed to Australopithecus afarensis from Hadar, Ethiopia. Journal of Human Evolution, 48: 593–642.

Drapeau, M.S.M., Ward, C. V., 2007. Forelimb segment length proportions in extant hominoids and *Australopithecus afarensis*. American Journal of Physical Anthropology. 132: 327–343.

Duren, D.L., 1999. Developmental determinants of femoral morphology (Doctoral dissertation, Kent State University).

Franciscus, R.G., Holliday, T.W., 1992. Hindlimb skeletal allometry in plio-pleistocene hominids with special reference to A.L.-288–1 (“Lucy”). Bulletins et Mémoires de la Société d’anthropologie de Paris. 4, 5–20.

Frost, H.M., 1990. Skeletal structural adaptations to mechanical usage (SATMU): 2. Redefining Wolff’s law: the remodeling problem. The anatomical record. 226, 414–422.

Galik, K., Senut, B., Pickford, M., Gommery, D., Treil, J., Kuperavage, A.J., Eckhardt, R.B., 2004. External and internal morphology of the BAR 1002’00 *Orrorin tugenensis* femur. Science. 305, 1450–1453.

Gordon, A.D., Green, D.J., Richmond, B.G., 2008. Strong postcranial size dimorphism in *Australopithecus afarensis:* results from two new resampling methods for multivariate data sets with missing data. American Journal of Physical Anthropology. 135, 311–328.

Grabowski, M., Hatala, K.G., Jungers, W.L. and Richmond, B.G., 2015. Body mass estimates of hominin fossils and the evolution of human body size. Journal of Human Evolution. 85, 75–93.

Green, D.J., Gordon, A.D. and Richmond, B.G., 2007. Limb-size proportions in *Australopithecus afarensis* and *Australopithecus africanus*. Journal of Human Evolution. 52, 187–200.

Grine, F.E., 2013. The alpha taxonomy of *Australopithecus africanus*. In: Reed, K.E., Fleagle, J.G., Leakey, R.E. (Eds.), The Paleobiology of Australopithecus. Springer, Dordrecht. 73–104.

Haeusler, M., McHenry, H.M., 2004. Body proportions of *Homo habilis* reviewed. Journal of Human Evolution. 46, 433–465.

Haeusler, M., McHenry, H.M., 2007. Evolutionary reversals of limb proportions in early hominids? evidence from KNM-ER 3735 (*Homo habilis*). Journal of Human Evolution. 53, 383–405.

Haile-Selassie, Y., Latimer, B.M., Alene, M., Deino, A.L., Gibert, L., Melillo, S.M., Saylor, B.Z., Scott, G.R., Lovejoy, C.O., 2010b. An early *Australopithecus afarensis* postcranium from Woranso-Mille, Ethiopia. Proceedings of the National Academy of Sciences. 107, 12121–12126.

Haile-Selassie, Y., Saylor, B.Z., Deino, A., Levin, N.E., Alene, M., Latimer, B.M., 2012. A new hominin foot from Ethiopia shows multiple Pliocene bipedal adaptations. Nature. 483, 565–569.

Halsey, L.G., Coward, S.R.L., Crompton, R.H., Thorpe, S.K.S., 2017. Practice makes perfect: Performance optimisation in ‘arboreal’ parkour athletes illuminates the evolutionary ecology of great ape anatomy. Journal of Human Evolution. 103, 45–52.

Hammer, Ø. & Harper, D.A.T. 2006. Paleontological Data Analysis. Blackwell.

Hammer, Ø., Harper, D.A.T., Ryan, P.D. 2001. PAST: Paleontological statistics software package for education and data analysis. Palaeontologia Electronica. 4, 1–9.

Hamrick MW. 1999. A chondral modeling theory revisited. Journal of Theoretical Biology. 201, 201–208.

Hartwig-Scherer, S., Martin, R.D., 1991. Was “Lucy” more human than her “child”? Observations on early hominid postcranial skeletons. Journal of Human Evolution. 21, 439–449.

Hatala, K.G., Demes, B., Richmond, B.G., 2016. Laetoli footprints reveal bipedal gait biomechanics different from those of modern humans and chimpanzees. Proceedings of the Royal Society B: Biological Sciences. 283, 20160235.

Heaton, J.L., Pickering, T.R. and R.J. Clarke (In Preparation) Body shape in StW 573 (Little Foot) and its potential phylogenetic implications.

Heile, A.J., Pickering, T.R., Heaton, J.L., R.J. Clarke., 2018. Bilateral Asymmetry of the Forearm Bones as Possible Evidence of Antemortem Trauma in the StW 573 *Australopithecus* Skeleton from Sterkfontein Member 2 (South Africa) bioRxiv 486076 [Preprint]. December 5, 2018; doi: https://doi.org/10.1101/486076

Heinrich, R.E., Rose, M.D., Leakey, R.E., Walker, A.C., 1993. Hominid radius from the middle Pliocene of Lake Turkana, Kenya. American Journal of Physical Anthropology. 92, 139–148.

Holliday, T.W., 1999. Brachial and crural indices of European Late Upper Paleolithic and Mesolithic humans. Journal of Human Evolution. 36, 549–566.

Holliday, T.W., 2012. Body Size, Body Shape, and the Circumscription of the Genus *Homo*. Current Anthropology. 53, S330–S345.

Holliday, T.W., Franciscus, R.G., 2012. Humeral Length Allometry in African Hominids (sensu lato) with Special Reference to A.L. 288-1 and Liang Bua 1. PaleoAnthropology. 2012, 1–12.

Holliday, T.W. and Ruff, C.B., 2001. Relative variation in human proximal and distal limb segment lengths. American Journal of Physical Anthropology 116, 26–33.

Howell, F.C., Wood, B.A., 1974. Early hominid ulna from the Omo basin, Ethiopia. Nature. 249, 174–176.

Jantz, L.M. and Jantz, R.L., 1999. Secular change in long bone length and proportion in the United States, 1800–1970. American Journal of Physical Anthropology. 110, 57–67.

Jashashvili, T. et al. (In Preparation) The hand of StW 573—description and virtual reconstruction.

Johanson, D.C., Lovejoy, C.O., Kimbel, W.H., White, T.D., Ward, S.C., Bush, M.E., Latimer, B.M., Coppens, Y., 1982. Morphology of the Pliocene partial hominid skeleton (A.L. 288-1) from the Hadar formation, Ethiopia. American Journal of Physical Anthropology. 57, 403–451.

Johanson, D.C., Masao, F.T., Eck, G.G., White, T.D., Walter, R.C., Kimbel, W.H., Asfaw, B., Manega, P., Ndessokia, P. and Suwa, G., 1987. New partial skeleton of *Homo habilis* from Olduvai Gorge, Tanzania. Nature, 327(6119), p.205.

Jolicoeur, P. 1963. The multivariate generalization of the allometry equation. Biometrics 19:497–499.

Jungers, W.L., 1982. Lucy’s limbs: Skeletal allometry and locomotion in *Australopithecus afarensis*. Nature. 297, 676–678.

Jungers, W.L., 1985. Body size and scaling of limb proportions in primates. In: Jungers, W.L., (Ed.), Size and scaling in primate biology. Springer, Boston, MA. 345–381.

Jungers, W.L., 1991. A pygmy perspective on body size and shape in *Australopithecus afarensis* (A.L. 288-1,”Lucy”). Origine (s) de la bipédie chez les hominidés. 215–224.

Jungers, W.L., 2009. Interlimb proportions in humans and fossil hominins: variability and scaling. In: Grine, F.E., Fleagle, J.G., Leakey, R.E., (Eds.), The First Humans: Origin and Early Evolution of the Genus Homo. pp. 93–98.

Jungers, W.L., Grabowski, M., Hatala, K.G., Richmond, B.G., 2016. The evolution of body size and shape in the human career. Philosophical Transactions of the Royal Society B: Biological Sciences. 371, 20150247.

Jungers, W.L., Stern, J.T., 1983. Body proportions, skeletal allometry and locomotion in the Hadar hominids: a reply to Wolpoff. Journal of Human Evolution. 12, 673–684.

Kimbel, W.H., Johanson, D.C., Rak, Y., 1994. The first skull and other new discoveries of Australopithecus afarensis at Hadar, Ethiopia. Nature. 368, 449–451.

Kimbel, W.H., Rak, Y. and Johanson, D.C., 2004. The skull of *Australopithecus afarensis*. Oxford University Press.

King, J.W., Brelsford, H.J. and Tullos, H.S., 1969. Analysis of the Pitching Arm of the Professional Baseball Pitcher. Clinical Orthopaedics and Related Research. 67, 116–123.

Klingenberg, C.P., 1996a. Individual variation of ontogenies: a longitudinal study of growth and timing. Evolution. 50, 2412–2428.

Klingenberg, C.P., 1996b. Multivariate allometry. In: Marcus, L.F., Corti, M., Loy, A., Naylor, G.J., Slice, D.E., (Eds.), Advances in morphometrics. Springer, Boston, MA. 23–49

Klingenberg, C.P., 2016. Size, shape, and form: concepts of allometry in geometric morphometrics. Development genes and evolution. 226, 113–137.

Klingenberg, C.P. and Spence, J.R., 1993. Heterochrony and allometry: lessons from the water strider genus *Limnoporus*. Evolution. 47, 1834–1853.

Knussman, R. (1967). Humerus, Ulna und Radius der Simiae: vergleichend-morphologische Untersuchungen mit Berücksichtigung der Funktion. Basel, New York, Karger.

Korey, K.A., 1990. Deconstructing reconstruction: The OH 62 humerofemoral index. American Journal of Physical Anthropology. 83, 25–33.

Kowalewski, M., E. Dyreson, J.D. Marcot, J.A. Vargas, K.W. Flessa. D.P. Hallmann. 1997. Phenetic discrimination of biometric simpletons: paleobiological implications of morphospecies in the lingulide brachiopod Glottidia. Paleobiology. 23, 444–469.

Kramer, P.A. and Sylvester, A.D., 2013. Humans, geometric similarity and the Froude number: is ‘‘reasonably close’’ really close enough?. Biology Open. 2, 111–120.

Lague, M.R., 2002. Another look at shape variation in the distal femur of *Australopithecus afarensis:* Implications for taxonomic and functional diversity at Hadar. Journal of Human Evolution. 42, 609–626.

Larson, S.G., 1996. Estimating humeral torsion on incomplete fossil anthropoid humeri. Journal of Human Evolution. 31, 239–257.

Larson, S.G., 2009. Evolution of the hominin shoulder: early Homo. In: Grine, F.E., Fleagle, J.G., Leakey, R.E., (Eds.), The First Humans: Origin and Early Evolution of the Genus Homo. Springer, Dordrecht. 65–75.

Leakey, M.D., Hay, R.L., 1979. Pliocene footprints in the Laetoli Beds at Leatoli, northern Tanzania. Nature. 278, 323–327.

Lordkipanidze, D., Jashashvili, T., Vekua, V., Ponce de Leon, M.S., Zollikofer, C.P.E., Rightmire, G.P., Pontzer, H., Ferring, R., Oms, O., Tappen, M., Bukhsianidze, M., Agusti, J., Kahlke, R., Kiladze, G., Martinez-Navarro, B., Mouskhelishvili, A., Nioradze, M., Rook, L., 2007. Postcranial evidence from early Homo from Dmanisi, Georgia. Nature. 449, 305–310.

Lovejoy, C.O., 1979. Reconstruction of the pelvis of A.L.-288 (Hadar Formation, Ethiopia). American Journal of Physical Anthropology. 460.

Lovejoy, C.O., 2007. The natural history of human gait and posture: Part 3. The knee. Gait & posture. 25, 325–341.

Lovejoy, C.O., Heiple, K.G., 1970. A reconstruction of the femur of *Australopithecus africanus*. American Journal of Physical Anthropology. 32, 33–40.

Lovejoy, C.O. and Heiple, K.G., 1972. Proximal femoral anatomy of *Australopithecus*. Nature. 235, 175.

Lovejoy, C.O., Suwa, G., Simpson, S.W., Matternes, J.H., White, T.D., 2009a. The great divides: *Ardipithecus ramidus* reveals the postcrania of our last common ancestors with African apes. Science. 326, 100–106.

Lovejoy, C.O., Suwa, G., Spurlock, L., Asfaw, B., White, T.D., 2009b. The pelvis and femur of *Ardipithecus ramidus:* The emergence of upright walking. Science. 326, 71e1–71e6.

Marchi, D., 2015. Using the morphology of the hominoid distal fibula to interpret arboreality in *Australopithecus afarensis*. Journal of Human Evolution. 85, 136–148.

Martin, R., 1928. Lehrbuch der Anthropologie: in systematischer Darstellung mit besonderer Berücksichtigung der anthropologischen Methoden; für Studierende, Ärzte und Forschungsreisende. 3. Bibliographie, Literaturverzeichnis, Sachregister, Autorenregister. G. Fischer.

Martin, R., Saller, K., 1928. Lehrbuch der Anthropologie in systematischer Darstellung, Bd. II.

Martin, R., Saller, K., 1959. Lehrbuch der Anthropologie. Stuttgart: Gustav-Fischer.

Masao, F.T., Ichumbaki, E.B., Cherin, M., Barili, A., Boschian, G., Iurino, D.A., Menconero, S., Moggi-Cecchi, J., Manzi, G., 2016. New footprints from Laetoli (Tanzania) provide evidence for marked body size variation in early hominins. eLife. 5, 29.

McHenry, H.M., 1978. Fore-and hindlimb proportions in Plio-Pleistocene hominids. American Journal of Physical Anthropology. 49, 15–22.

McHenry, H.M., 1991. Petite bodies of the “robust” australopithecines. American Journal of Physical Anthropology. 86, 445–454.

McHenry, H.M., 1992. Body size and proportions in early hominids. American Journal of Physical Anthropology. 87, 407–431.

McHenry, H.M., Berger, L.R., 1998a. Body proportions in *Australopithecus afarensis* and *A. africanus* and the origin of the genus *Homo*. Journal of Human Evolution. 35, 1–22.

McHenry, H.M., Berger, L.R., 1998b. Limb lengths in *Australopithecus* and the origin of the genus *Homo*. South African Journal of Science. 94, 447–450.

McHenry, H.M., Brown, C.C., McHenry, L.J., 2007. Fossil hominin ulnae and the forelimb of *Paranthropus*. American Journal of Physical Anthropology. 134, 209–218.

McHenry, H.M., Corruccini, R.S., Howell, F.C., 1976. Analysis of an early hominid ulna from the Omo Basin, Ethiopia. American Journal of Physical Anthropology. 44, 295–304.

McHenry, H.M. and Corruccini, R.S., 1978. The femur in early human evolution. American Journal of Physical Anthropology. 49, 473–487.

McHenry, H.M., Jones, A.L., 2006. Hallucial convergence in early hominids. Journal of Human Evolution. 50, 534–539.

Migliano, A.B., Vinicius, L., Lahr, M.M., 2007. Life history trade-offs explain the evolution of human pygmies. Proceedings of the National Academy of Sciences. 104, 20216–20219.

Miller, R.A., 1932. Evolution of the pectoral girdle and forelimb in the primates. American Journal of Physical Anthropology. 17, 1–56.

Napier, J.R., 1964. The evolution of bipedal walking in the hominids. Arch. Biol. (Liege) 1964;75:673–708.

Ohman, J.C., Wood, C., Wood, B., Crompton, R.H., Günther, M.M., Yu, L., Savage, R., Wang, W., 2002. Stature-at-death of KNM-WT 15000. Human Evolution. 17, 129–141.

Partridge, T.C., Granger, D.E., Caffee, M.W. and Clarke, R.J., 2003. Lower Pliocene hominid remains from Sterkfontein. Science, 300, 607–612.

Pearson, O.M., 1997. Postcranial morphology and the origin of modern humans. Doctoral Dissertation. State University of New York at Stony Brook.

Pearson, O.M., 2000. Postcranial remains and the origin of modern humans. Evolutionary Anthropology. 9, 229–247.

Pickering, T.R., Heaton, J.L., Clarke, R.J., Stratford, D., 2018. Hominin hand bone fossils from Sterkfontein Caves, South Africa (1998 – 2003 excavations). Journal of Human Evolution. 118, 89–102.

Pickering, T.R., Heaton, J.L., Clarke, R.J., Stratford, D., 2019. Hominin vertebrae and upper limb bone fossils from Sterkfontein Caves, South Africa (1998 – 2003 excavations). American Journal of Physical Anthropology (in press).

Pickford, M., Senut, B., Gommery, D., Treil, J., 2002. Bipedalism in *Orrorin tugenensis* revealed by its femora. Comptes Rendus – Palevol. 1, 191–203.

Pieper, H.G., 1998. Humeral torsion in the throwing arm of handball players. The American Journal of Sports Medicine. 26, 247–253.

Pontzer, H., 2007. Predicting the energy cost of terrestrial locomotion: a test of the Limb model in humans and quadrupeds. Journal of Experimental Biology. 210, 484–494.

Pontzer, H., 2012. Ecological Energetics in Early *Homo*. Current Anthropology. 53, S346–S358.

Pontzer, H., Raichlen, D.A., Sockol, M.D., 2009. The metabolic cost of walking in humans, chimpanzees, and early hominins. Journal of Human Evolution. 56, 43–54.

Pontzer, H., Rolian, C., Rightmire, G.P., Jashashvili, T., Ponce de León, M.S., Lordkipanidze, D., Zollikofer, C.P.E., 2010. Locomotor anatomy and biomechanics of the Dmanisi hominins. Journal of Human Evolution. 58, 492–504.

Raichlen, D.A., Gordon, A.D., 2017. Interpretation of footprints from Site S confirms human-like bipedal biomechanics in Laetoli hominins. Journal of Human Evolution. 107, 134–138.

Reed, K.E., Kitching, J.W., Grine, F.E., Jungers, W.L., Sokoloff, L., 1993. Proximal femur of *Australopithecus africanus* from member 4, Makapansgat, South Africa. American Journal of Physical Anthropology. 92, 1–15.

Rein, T.R., Harrison, T., Carlson, K.J. and Harvati, K., 2017. Adaptation to suspensory locomotion in *Australopithecus sediba*. Journal of Human Evolution. 104:1–12.

Reno, P., Degusta, D., Serrat, M., Meindl, R., White, T., Eckhardt, R., Kuperavage, A., Galik, K., Lovejoy, C.O., Reno, P.L., Serrat, M.A., Meindl, R.S., White, T.D., Eckhardt, R.B., Kuperavage, A.J., 2005. Plio-Pleistocene Hominid Limb Proportions Evolutionary Reversals or Estimation Errors? Current Anthropology. 46, 575–588.

Reno, P.L., Lovejoy, C.O., 2015. From Lucy to Kadanuumuu: balanced analyses of *Australopithecus afarensis* assemblages confirm only moderate skeletal dimorphism. PeerJ. 3, e925.

Reno, P.L., Meindl, R.S., McCollum, M.A., Lovejoy, C.O., 2003. Sexual dimorphism in *Australopithecus afarensis* was similar to that of modern humans. Proceedings of the National Academy of Sciences. 100, 9404–9409.

Reno, P.L., Serrat, M.A., Meindl, R.S., Tim, D., Eckhardt, R.B., Kuperavage, A.J., Lovejoy, C.O., 2005. Plio-Pleistocene Hominid Limb Proportions. Current Anthropology. 46, 575–588.

Richmond, B.G., Aiello, L.C., Wood, B.A., 2002. Early hominin limb proportions. Journal of Human Evolution. 43, 529–548.

Richmond, B.G., Jungers, W.L., 2008. Orrorin tugenensis Femoral Morphology and the Evolution of Hominin Bipedalism. Science. 319, 1662–1665.

Richmond, B.G., Strait, D.S., 2000. Evidence that humans evolved from a knuckle-walking ancestor. Nature. 404, 382–385.

Robinson, J.T., 1972. Early hominid posture and locomotion. University of Chicago Press.

Ruff, C.B., 1994. Morphological adaptation to climate in modern and fossil hominids. American Journal of Physical Anthropology. 37, 65–107.

Ruff, C., 2009. Relative limb strength and locomotion in *Homo habilis*. American Journal of Physical Anthropology. 138, 90–100.

Ryan, T.M., Sukhdeo, S., 2016. KSD-VP-1/1: Analysis of the postcranial skeleton using high-resolution computed tomography. In: Haile-Selassie, Y., Su, D.F., (Eds.), The Postcranial Anatomy of Australopithecus Afarensis. Springer, 39–62.

Sarmiento, E.E., 1985. Functional differences in the skeleton of wild and captive orangutans and their adaptive significance. Ph.D. dissertation, New York University, New York.

Schmid, P., 1983. Eine rekonstruktion des skelettes von A.L. 288–1 (Hadar) und deren Konsequenzen. Folia Primatologica. 40, 283–306.

Schultz, A.H., 1937. Proportions, variability and asymmetries of the long bones of the limbs and the clavicles in man and apes. Human Biology. 9, 281–328.

Senut B, Tardieu C., (1985) Functional aspects of Plio-Pleistocene hominid limb bones: implications for taxonomy and phylogeny. In: Delson, E., (Eds), Ancestors: The Hard Evidence. New York: Alan R. Liss, Inc. 193–201.

Skelton, R.R. and McHenry, H.M., 1992. Evolutionary relationships among early hominids. Journal of Human Evolution. 23, 309–349.

Skelton, R.R., McHenry, H.M., Drawhorn, G.M., Bilsborough, A., Chamberlain, A.T., Wood, B.A. and Vančata, V., 1986. Phylogenetic Analysis of Early Hominids [and Comments and Reply]. Current Anthropology. 27, 21–43.

Sockol, M.D., Raichlen, D.A., Pontzer, H., 2007. Chimpanzee locomotor energetics and the origin of human bipedalism. Proceedings of the National Academy of Sciences. 104, 12265–12269.

Stern, J.T., Susman, R.L., 1983. The locomotor anatomy of *Australopithecus afarensis*. American Journal of Physical Anthropology. 60, 279–317.

Steudel-Numbers, K.L., 2006. Energetics in *Homo erectus* and other early hominins: The consequences of increased lower-limb length. Journal of Human Evolution. 51, 445–453.

Steudel-Numbers, K.L., Tilkens, M.J., 2004. The effect of lower limb length on the energetic cost of locomotion: implications for fossil hominins. Journal of Human Evolution. 47, 95–109.

Strait, D.S. and Grine, F.E., 2004. Inferring hominoid and early hominid phylogeny using craniodental characters: the role of fossil taxa. Journal of Human Evolution. 47, 399–452.

Strait, D.S., Grine, F.E. and Moniz, M.A., 1997. A reappraisal of early hominid phylogeny. Journal of Human Evolution. 32, 17–82.

Stock, J.T., Shaw, C.N., 2007. Which Measures of Diaphyseal Robusticity Are Robust? A comparison of external methods of quantifying the strength of long bone diaphyses to cross-sectional geometric properties. American Journal of Physical Anthropology. 134, 412–423.

Susman, R.L., Stern, J.T., Jungers, W.L., 1984. Arboreality and bipedality in the Hadar hominids. Folia Primatologica. 43, 113–156.

Susman, R.L., Stern, J.T., Jungers, W.L., 1985. Locomotor adaptations in the Hadar hominids. In: Delson, E., (Eds), Ancestors: The Hard Evidence. Alan R. Liss New York, 184–192.

Sylvester, A.D., Kramer, P.A. and Jungers, W.L. 2008. Modern humans are not (quite) isometric. American Journal of Physical Anthropology. 137, 371–383.

Tallman, M., Almécija, S., Reber, S.L., Alba, D.M., Moyà-Solà, S., 2013. The distal tibia of *Hispanopithecus laietanus:* More evidence for mosaic evolution in Miocene apes. Journal of Human Evolution. 64, 319–327.

Tardieu, C., 2010. Development of the human hind limb and its importance for the evolution of bipedalism. Evolutionary Anthropology: Issues, News, and Reviews. 19, 174–186.

Trinkaus, E., 1981. Neanderthal limb proportions and cold adaptation. In: Stringer, C.B. (Ed.), Aspects of human evolution. London: Taylor and Francis. 187–224.

Trinkaus, E., 1983. Neandertal postcrania and the adaptive shift to modern humans. The Mousterian Legacy: Human Biocultural Change in the Upper Pleistocene. 164, 165–200.

Trinkaus, E., Stringer, C.B., Ruff, C.B., Hennessy, R.J., Roberts, M.B., Parfitt, S.A., 1999. Diaphyseal cross-sectional geometry of the Boxgrove 1 Middle Pleistocene human tibia. Journal of Human Evolution. 37, 1–25.

Tuttle, R.H., 1985. Ape footprints and Laetoli impressions: a response to the SUNY claims. In: Tuttle, R.H., Tobias, P.V., Liss, A.R. (Eds.), Hominid evolution: Past, Present and Future. 129–133.

Villa, P., Mahieu, E., 1991. Breakage patterns of human long bones. Journal of Human Evolution. 21, 27–48.

Walker, A., Leakey, R.E., 1993. The Nariokotome *Homo erectus* skeleton. Harvard University Press.

Wall-Scheffler, C.M., Myers, M.J., 2013. Reproductive costs for everyone: how female loads impact human mobility strategies. Journal of Human Evolution. 64, 448–456.

Wang, W., Crompton, R.H., Carey, T.S., Günther, M.M., Li, Y., Savage, R., Sellers, W.I., 2004. Comparison of inverse-dynamics musculo-skeletal models of A.L. 288–1 *Australopithecus afarensis* and KNM-WT 15000 *Homo ergaster* to modern humans, with implications for the evolution of bipedalism. Journal of Human Evolution. 47, 453–478.

Wang, W.J., Crompton, R.H., 2003. Size and power required for motion with implication for the evolution of early hominids. Journal of Biomechanics. 36, 1237–1246.

Wang, W.J., Crompton, R.H., Li, Y., Gunther, M.M., 2003. Optimum ratio of upper to lower limb lengths in hand-carrying of a load under the assumption of frequency coordination. Journal of Biomechanics. 36, 249–252.

Ward C.V., 1997. Functional anatomy and phyletic implications of the hominoid trunk and hindlimb. In: Begun D.R., Ward C.V., Rose M.D., (Eds.), Function, phylogeny, and fossils: Miocene Hominoid evolution and adaptations. New York: Plenum Press. 101–130.

Ward, C.V., 2002. Interpreting the posture and locomotion of *Australopithecus afarensis:* where do we stand?. American Journal of Physical Anthropology. 119, 185–215.

Ward, C.V., Leakey, M.G. and Walker, A., 2001. Morphology of Australopithecus anamensis from Kanapoi and Allia Bay, Kenya. Journal of Human Evolution. 41, 255–368.

Watson, J.C., Payne, R.C., Chamberlain, A.T., Jones, R.K., Sellers, W.I., 2008. The energetic costs of load-carrying and the evolution of bipedalism. Journal of Human Evolution. 54, 675–683.

Wilder, H.H., 1920. A laboratory manual of anthropometry. P. Blakiston’s Son and Co., Philadelphia, Pa.

Wolpoff, M.H., 1983. Lucy’s little legs. Journal of Human Evolution. 12, 443–453.

Young, N.M., Wagner, G.P., Hallgrimsson, B., 2010. Development and the evolvability of human limbs. Proceedings of the National Academy of Sciences. 107, 3400–3405.

Zipfel, B., DeSilva, J.M., Kidd, R.S., Carlson, K.J., Churchill, S.E., Berger, L.R., 2011. The foot and ankle of *Australopithecus sediba*. Science, p.1202703.

